# Robust identification of regulatory variants (eQTLs) using a differential expression framework developed for RNA-sequencing

**DOI:** 10.1101/2022.11.18.517114

**Authors:** Mackenzie A. Marrella, Fernando H. Biase

## Abstract

**Background:** A gap currently exists between genetic variants and the underlying cell and tissue biology of a trait, and expression quantitative trait loci (eQTL) studies provide important information to help close that gap. However, two concerns that arise with eQTL analyses using RNA-sequencing data are normalization of data across samples and the data not following a normal distribution. Multiple pipelines have been suggested to address this. For instance, the most recent analysis of the human and farm Genotype-Tissue Expression (GTEx) project proposes using trimmed means of M-values (TMM) to normalize the data followed by an inverse normal transformation.

**Results:** In this study, we reasoned that eQTL analysis could be carried out using the same framework used for differential gene expression (DGE), which uses a negative binomial model, a statistical test feasible for count data. Using the GTEx framework, we identified 38 significant eQTLs (P<5×10^-8^) following the ANOVA model and 15 significant eQTLs (P<5×10^-8^) following the additive model. Using a differential gene expression framework, we identified 2,471 and nine significant eQTLs (P<5×10^-8^) following an analytical framework equivalent to the ANOVA and additive model, respectively. When we compared the two approaches, there was no overlap of significant eQTLs between the two frameworks. Because we defined specific contrasts, we identified trans eQTLs that more closely resembled what we expect from genetic variants showing complete dominance between alleles. Yet, these were not identified by the GTEx framework.

**Conclusions:** Our results show that transforming RNA-sequencing data to fit a normal distribution prior to eQTL analysis is not required when the DGE framework is employed, thus this may be more suitable for finding genes whose expression are impacted by genetic variants. Our approach detected biologically relevant variants that otherwise would not have been identified due to data transformation to fit a normal distribution.

## Background

Recent publications from the human Genotype-Tissue Expression (GTEx) (1, 2) and the cattle GTEx (3, 4) projects have shed light on the genetic control of gene expression in large mammals. The recent findings indicate that genomic variants have a greater impact on gene expression than previously anticipated (5). These studies have provided valuable information which will help close the critical gap between genomic variants and phenotypic variation (6, 7), especially those associated with health in humans and livestock.

Given the importance of identifying expression quantitative trait loci (eQTL) (8) to understand cell or tissue biology, several statistical approaches have emerged to allow the coordinated analysis of genomic variants and transcript abundance (reviewed by Nica and Dermitzakis (8)). While the first eQTL studies used microarray data (9), most of the analyses carried out in recent years use RNA-sequencing data. One emerging concern is the normalization of the data across samples. To that end, several methods have been used for data normalization across samples such as the trimmed mean of M-values (TMM) (10), fragments per kilobase per million reads (FPKM) (11), and transcript per million reads (TPM) (12). These and other methods have been evaluated, and TMM might have an advantage over other methods (13). Another concern related to eQTL analysis is that RNA-sequencing data do not follow a normal distribution, however, all statistical approaches currently employed assume that the inputted data will follow a normal distribution. Researchers have addressed this by transforming the data using the variance stabilization (14–16), Log_2_ transformation (17, 18), or the inverse normal transformation (1, 3, 4, 19, 20).

Because the principle of eQTL analysis is to identify differences in transcript abundance between genotypes (9), we reasoned that the analysis of eQTLs using transcript abundance estimated from RNA-sequencing could be carried out using the same framework used for differential gene expression. A major benefit of using such a framework is that DGE is estimated using a negative binomial model (14, 21, 22), which is suitable for sequence count data (23, 24). Here, our objective was to identify eQTLs in cattle peripheral white blood cells (PWBCs) using RNA-sequencing data and the Bioconductor (25) package “edgeR” (21, 26), which was designed for DGE analysis using the general linear model framework.

## Methods

All bioinformatics and analytical procedures are presented in Additional file 1.

### Data processing for variant detection, and variant filtering

We analyzed RNA-sequencing data from 42 heifers (*Bos taurus*, Angus x Simmental) publicly available in the GEO database: GSE103628 (27, 28) and GSE146041 (29). Hisat2 (v.2.2.0) (30) was used to align the pair-end short reads to the cattle genome (31, 32) (Bos_taurus.ARS-UCD1.2.99), obtained from the Ensembl database (33). Next, we used Samtools (v.1.10) (34) to filter reads that did not map, secondary alignments, alignments from reads that failed platform/vendor quality checks, and were PCR or optical duplicates. Duplicates were removed using the function “bammarkduplicates” from biobambam2 (2.0.95) (35). The function “SplitNCigarReads” from GATK (v.4.2.2.0) (36) was then used to separate sequences with a CIGAR string, which resulted from sequencing exon-exon boundaries. Variants were then called in our data by using the functions “bcftools mpileup” and “bcftools call” from Samtools (34).

We filtered the variants with the function “bcftools view” from Samtools to select sites where 20 or more reads were used to identify a variant. Next, in R software (4.0.3) (37), we retained variant sites that were identified as single nucleotide polymorphisms and retained variants with genotypes called in at least 20 samples.

### Variant annotation

After the list of significant SNP-gene pairs was generated from the eQTL analysis, attributes were read in from the Ensembl genome database. The attribute list was merged with the output from the eQTL analysis as well as the nucleotide genotypic data for all samples. Ensembl Variant Effect Predictor (38) was used to compare our data to the cattle genome (Bos taurus, ARS-UCD1.2) to identify the functional consequences of the SNPs.

### Quantification of transcript abundance

For the expression dataset, we obtained the raw read counts from our previous work (29). First, we eliminated one sample that had less than a million reads mapped to the annotation; second, we calculated counts per million reads (CPM) (21); third, we retained protein-coding genes that had CPM greater than two in five or more samples. Next, we calculated TPM (39), which was used in all plots with transcript abundance.

### eQTL analysis

First, we tested whether the samples presented a genetic stratification using plink (40) to calculate the eigenvectors (41). Given the sample elimination due to low mapping to the annotation, we carried out an eQTL analysis with 41 samples. To prevent overinflation of effects when working with variants with low allelic frequencies (42) and conduct a robust analysis with enough samples in each group of genotypes, we further retained those single nucleotide polymorphisms that had at least five animals in each of the two homozygotes and heterozygote genotypes, had a minor allelic frequency > 0.15, and followed Hardy-Weinberg equilibrium (false discovery rate=0.05), which was tested with the R package “HardyWeinberg” (43). In both approaches described below, eQTLs that overlapped between the ANOVA and additive model are only reported in the ANOVA model.

#### Approach 1: TMM normalized and normal-transformed RNA-seq data

In line with standard procedures adopted for eQTL analysis (1, 19, 20), we normalized expression abundance for 10,332 genes using the TMM method (10). First, we used the function “calcNormFactors” from the R package “edgeR” (21, 26) to calculate the normalization factors then we multiplied the normalization factors by the respective library size. Next, we used the function “cpm” with the normalized library size to obtain TMM normalized counts per million. Next, we carried out an inverse normal transformation (1, 19, 20) using the “RankNorm” function from the R package “RNOmni”. Additive and ANOVA analyses were carried out independently for eQTL analysis with the R package “MatrixEQTL” (44) using 6,216 SNPs. We inferred a significant eQTL when the nominal P-value was less than 5 x 10^-8^, which is a threshold commonly applied to genome-wide association studies (45–49), and corresponded to a false discovery rate (50) of 4% and 12% for the ANOVA and additive model, respectively.

#### Approach 2: Using a differential gene expression framework

We analyzed the RNA-sequencing data with a general linear model in “edgeR” and tested for differential gene expression using the quasi-likelihood F-test (51, 52). We note that the normalization adopted by default in “edgeR” adjusts for library sequencing depth, but we added the TMM normalization factors calculated by the function “calcNormFactors” to the procedure for identification of eQTL.

For additive analysis, the genotypes were input as numerical independent variables, and the gene expression data was used as a dependent variable. As part of our proposed approach, we also eliminated genes that had outlier values of transcript abundance, which reduced the transcriptome data to 4,149 genes. For ANOVA-like analysis, we carried out a two-tier analysis. First, we tested the association between SNP and gene transcript abundance using all three genotypes. Next, we subset SNPs that were significantly associated with gene transcript abundance and pseudo-coded the genotypes to establish two contrasts (53). The first contrast compared the homozygote genotype from the reference allele versus the heterozygote and the homozygote genotype from the alternate allele (i.e. AA versus AB, BB). The second contrast compared the homozygote genotype from the alternate allele versus the heterozygote and the homozygote genotype from the reference allele (i.e. AA, AB versus BB). We also inferred a significant eQTL when the nominal P-value was less than 5 x 10^-8^ (45–49).

### Visualization of the results

We used the R packages “ggplot2”, “cowplot” (54), or “plotly” (55) for plotting (56) and used Cytoscape (57) to visualize eQTLs in network style.

### Analysis of gene ontology enrichment

We tested several lists of genes for the enrichment of gene ontology using the R package “GOseq”(58). In order to account for multiple hypothesis testing, P-values were adjusted by family wise error rate (FWER) (59). Results were maintained if they had FWER <0.07.

## Results

### Overview of SNP identification

Our pipeline for SNP identification started with the alignment of reads to the reference genome with HISAT2 (30), which was followed by the filtering of secondary alignments using Samtools (60) and removal of duplicates using biobambam (35). Next, we used the function “SplitNCigarReads” from the GATK (61, 62) framework. Then, bcftools (63, 64) was used for variant calling and filtering. We opted for bcftools because it calls genotypes at each genomic position for all samples with sequencing coverage. To increase the confidence in our analysis, we filtered out the genotypes with coverage of fewer than 20 reads, genotypes that presented secondary alternate alleles, and those positions that were genotyped in fewer than 26 samples (Figure 1A). After filtering, we compiled genotypes at 23,506,613 nucleotide positions. Not surprisingly, 99.6% of the genomic positions were homozygous for the reference allele and 2,167 positions were homozygous for the alternate allele. Our pipeline identified 91,006 nucleotide positions showing polymorphisms in our samples.

**Figure 1.**
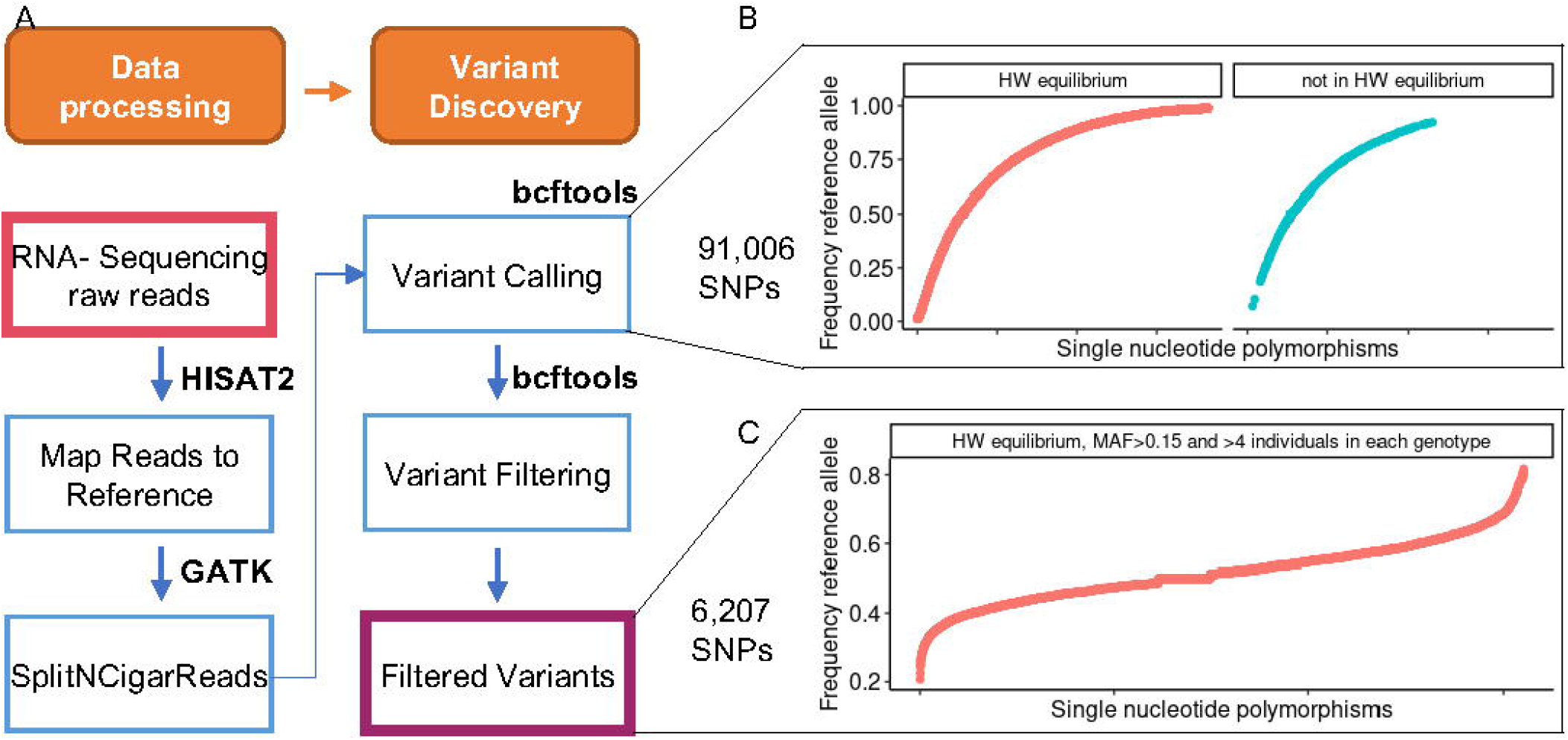
Overview of genotyping and variant discovery using RNA-sequencing data from PWBCs. **(A)** Schematics of bioinformatics procedures. **(B)** Distribution of allelic frequency of all variants genotyped in at least 26 samples. **(C)** Distribution of allelic frequency of all variants genotyped in at least 26 samples followed by filtering to retain 6,207 SNPs. (HW: Hardy-Weinberg; MAF: minimum allele frequency).

We then calculated the frequency of the reference allele and tested the deviation from Hardy-Weinberg equilibrium (Figure 1B). To prevent common pitfalls in the eQTL analysis (42), we removed SNPs from our initial pool of 91,006 that: (i) showed significant deviation from Hardy-Weinberg equilibrium (FDR<0.01); (ii) presented a minimum allelic frequency < 0.15; and (iii) had less than 5 individuals in each of the three genotypes. There were 6,207 SNPs retained for further analysis. Notably, 96% (n=5,964) of the SNPs have been previously identified and are recorded in the Ensembl variant database (65, 66), which includes the dbSNP ((67) version 150), while 243 SNPs were not identified in Ensembl variant database (Additional file 2). Most of the SNPs are in 3 prime UTRs (n=1,553), and a smaller proportion (n=483) were annotated as missense variants (Additional file 2). We observed no genetic substructure of the individuals based on the SNPs analyzed here (Figure S1, Additional file 3).

### eQTL analyses

For eQTL analysis, we obtained the matrix with raw counts from a previous study (29) from our group. After filtering for lowly expressed genes, we quantified the transcript abundance for 10,332 protein-coding genes. We then analyzed the transcriptome and the SNP data following the two frameworks.

#### Approach 1: TMM normalized and normal-transformed RNA-seq data

We analyzed the data using the normalization indicated in the GTEx project (1), which is the normalization of the counts using the TMM approach (10) followed by the inverse normal transformation within a gene and across samples (20). The transformation indeed normalized the RNA-sequencing data (Figure S2, Additional file 3). Using the R package “MatrixEQTL” (68), the ANOVA and additive analysis concluded in 4.699 and 2.473 seconds respectively using one core processor (2.60GHz).

We identified 35 significant eQTLs (P < 5×10^-8^) following the ANOVA model (Figure 2). Annotated SNPs mapped to the genes: *ASCC1, BOLA-DQB, FAF2, IARS2, MGST2, MRPS9, NECAP2, TRIP11* (Additional file 4). We also identified 39 significant eQTLs (P < 5×10^-8^) following the additive model (Figure 3). Annotated SNPs mapped to the genes *AHNAK, GLB1, TRIP11*, Additional file 5), and most of the SNPs on the gene *TRIP11* composed the majority of the eQTLs.

**Figure 2.**
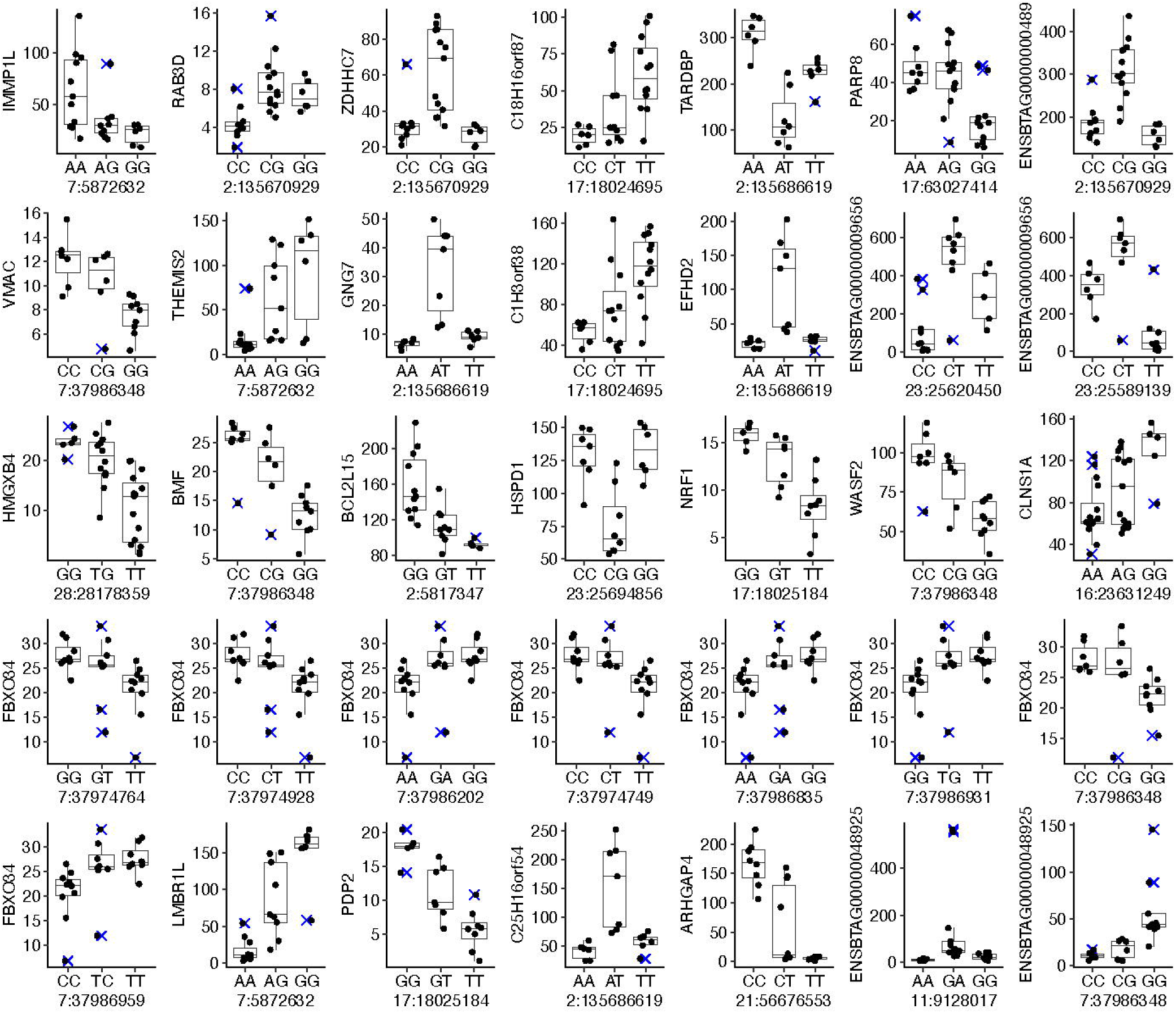
eQTLs identified using ANOVA model on TMM normalized counts per million and normal-transformed RNA-seq data. Y axis for all graphs is TMM normalized counts per million.

**Figure 3.**
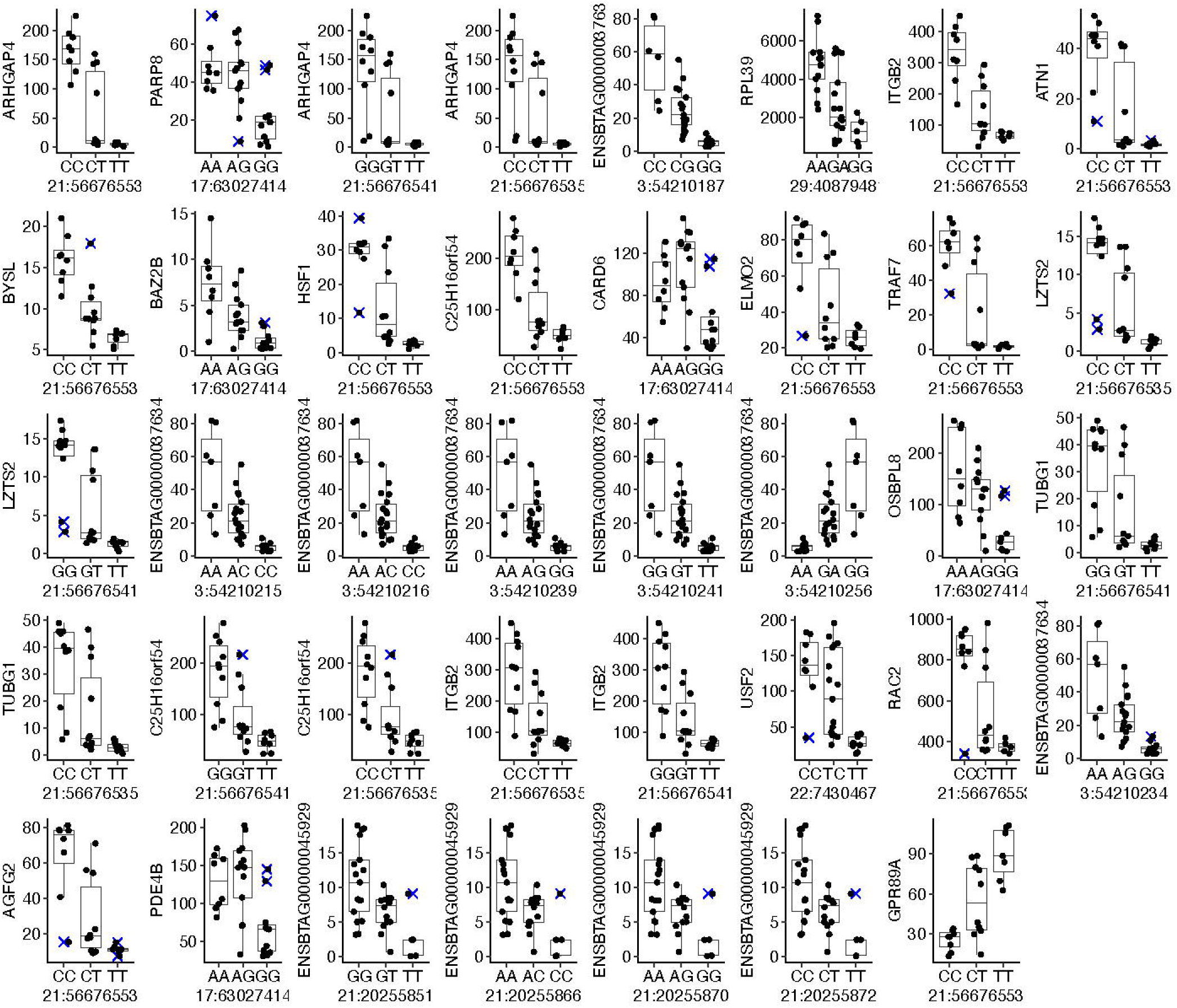
eQTLs identified using additive model on TMM normalized counts per million and normal-transformed RNA-seq data. Y axis for all graphs is TMM normalized counts per million.

#### Approach 2: Using a differential gene expression framework

We propose further filtering of the transcriptome data to remove outliers that resulted in 4,149 genes utilized for downstream analysis. Using the R package “edgeR” (21), all tests to determine dominance and additive models were completed in 36 and nine minutes respectively using 34 core processors (2.60GHz).

We identified 2,540 significant eQTLs (P < 5×10^-8^). These eQTLs were formed by 32 SNPs present in the dbSNP and one SNP that is a putatively new variant (Additional file 3) influencing the transcript abundance of 928 genes (Figure 4A). The majority (99.2%) of the eQTLs were formed by SNPs on the genes TATA-Box binding protein associated factor 15 (*TAF15*) and SMG6 nonsense-mediated mRNA decay factor (*SMG6*) (Figure 4B). There was no overlap of significant eQTL between both approaches (Additional file 6, Figures S3 and S4, Additional file 3).

**Figure 4.**
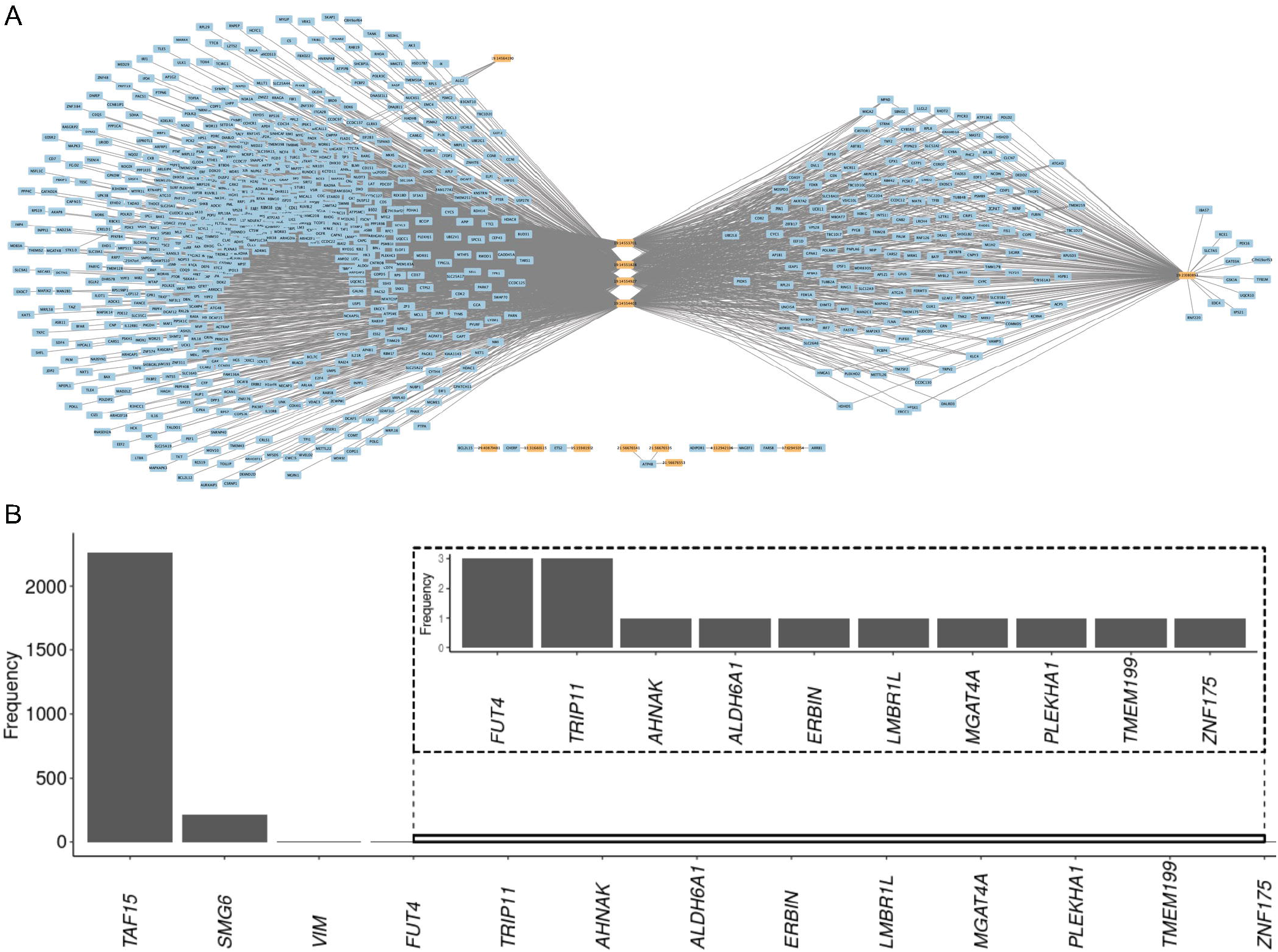
Significant eQTLs were identified using the differential gene expression framework. **(A)** Network depicting the connectivity between SNPs and the genes whose genotypes are influencing their transcript abundance. **(B)** Bar plot of the frequency of genes containing SNPs forming eQTLs. Only SNPs that were annotated to genes with a symbol (within a gene model, or within 1,000 nucleotides on each side) are depicted in this figure.

It was also possible to separate the eQTLs into dominance or additive allelic interaction. We determined that eight of the eQTLs followed the pattern of an additive allelic relationship (Figure 5A, Additional file 7). Two SNPs (rs41892216 and rs135008768) impacting the expression of the gene sialic acid-binding Ig-like lectin 14 are also present in the region containing the sialic acid-binding Ig-like lectin gene family on chromosome 18. One SNP is a missense mutation (18:57565792, Figure 5B) on the gene *SIGLEC5* and the SNP on nucleotide 18:57498163 is a variant downstream to *SIGLEC6*. Two other SNPs were annotated to the genes *AHNAK*, 29:40879481 and *MGAT4A*, 11:3860402.

**Figure 5.**
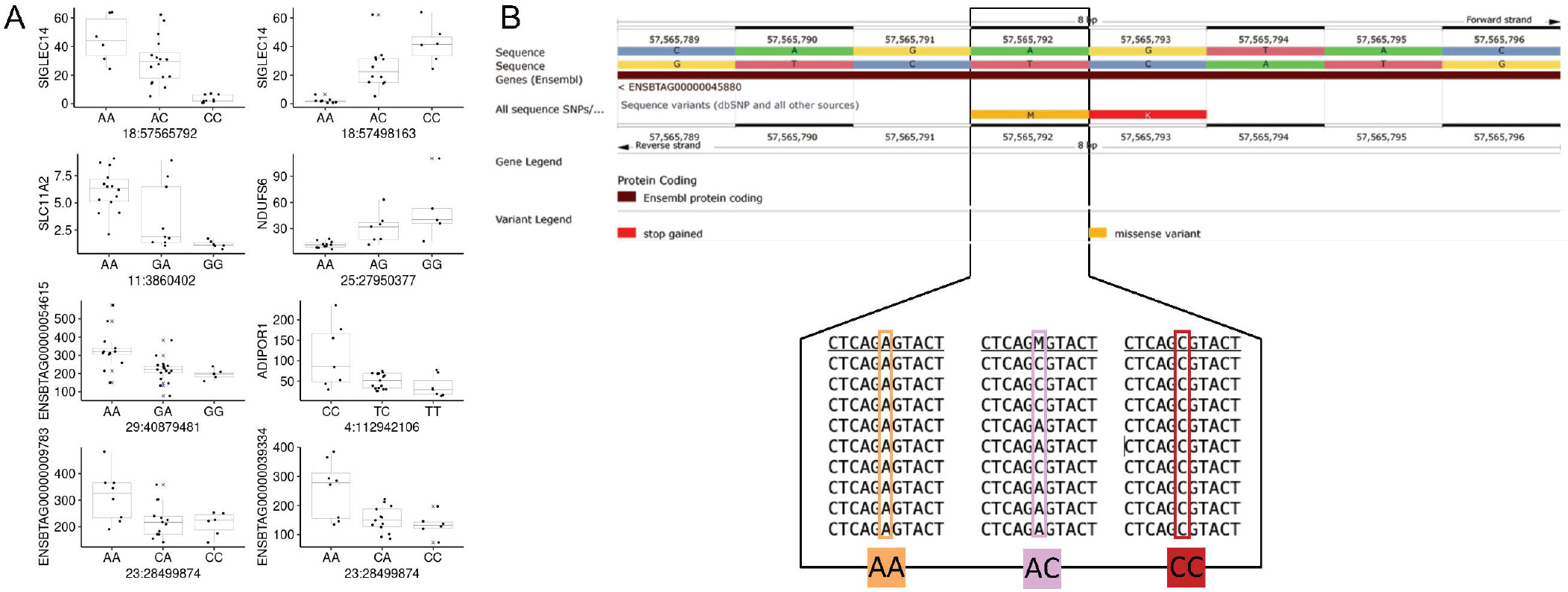
Significant eQTLs were inferred using the differential gene expression framework following the additive relationship between alleles. **(A)** Nine eQTLs following the additive model determined by edgeR. **(B)** Ensembl genome browser indicating the SNP position and examples of raw data used for the SNP’s identification.

We also identified 2533 significant eQTLs following a dominance allelic relationship (Additional file 8). Nine annotated SNPs mapped to the genes (*ALDH6A1, ERBIN*, FUT4, *LMBR1L, PLEKHA1, SMG6, TAF15, TMEM199, VIM, ZNF175*). Of notice, four intronic variants and one missense variant on the gene *TAF15* (Figure 6A) were collectively associated with the expression of 871 genes, with some examples depicted in Figure 6B.

**Figure 6.**
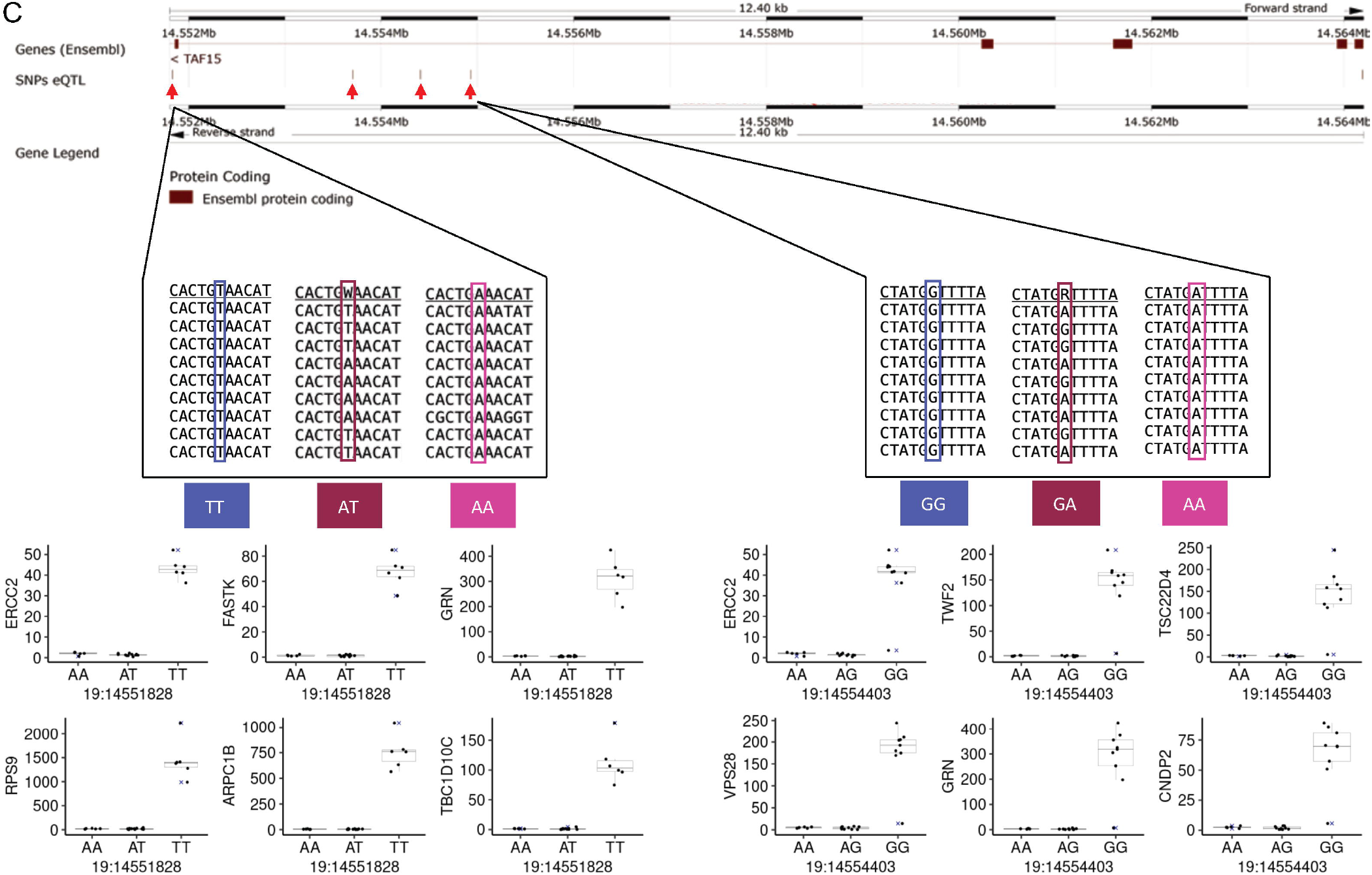
Significant eQTLs were inferred using the differential gene expression framework following the dominance relationship between alleles. **(A)** Ensembl genome browser indicating the SNP positions and examples of raw data used for the SNP’s identification on *TAF15*. **(B)** Examples of eQTLs determined by “edgeR” using the ANOVA-like analysis.

Given the number of genes expressed in PWBCs that were influenced by SNPs, we asked if there would be an enrichment of gene ontology (69) biological processes among these 871 genes. We observed that by setting a more stringent threshold of significance for the eQTLs (P < 5×10^-12^), we subset 132 genes, which are enriched for one biological process (FWER < 0.1): regulation of catalytic activity (fold-enrichment: 4.2; genes: *APBA3, ARHGDIA, ARHGEF1, CAPN1, DENND1C, EEF1D, RALGDS, RING1*, Additional file 9).

## Discussion

The major goal of our work was to identify genes expressed in PWBCs of crossbred beef heifers whose transcript abundance is impacted by genetic variants. We used a gold standard approach presented by the GTEx consortium, but also analyzed the RNA-sequencing data without a transformation to force a Gaussian distribution of the counts. The framework for eQTL analysis presented here is motivated by the following rationale: (i) the vast majority of eQTL analyses carried out currently use RNA-sequencing data; (ii) by the nature of the procedures, RNA-sequencing data is count data, which is not normally distributed (70, 71); and (iii) in principle, an eQTL analysis is an expansion of a differential gene expression (DGE) analysis, where samples are grouped by their genotypes, which is analogous to groups or treatments typically used in DGE analysis. Compared to the latest GTEx framework, our analysis of RNA-sequencing data from cattle PWBCs using the DGE framework identified more (100-fold difference) eQTLs under the dominance model and an equivalent number of eQTLs under the additive model of allele interaction when compared to the framework used in the human or farm GTEx consortia.

Our study has a few limitations, but they do not hinder the validity of our findings. First, we identified SNPs using the RNA-sequencing data, thus we are not accounting for genomic variants in promoters or distal cis-regulatory elements. This is likely to have impacted the limited number of cis-eQTLs reported here. Second, our transcriptome data represents a mixture of white cells identified in the blood. The proportion of different cells that compose the mixture of white cells was not accounted in our model. A genetic factor contributing to a potential greater abundance of one specific cell type (72) is thus a confounding factor in our study. However, these two limitations do not directly impact our main take home message that there is no need for researchers to normalize RNA-sequencing data in eQTL studies.

### Variant genotyping using RNA-sequencing data

RNA-sequencing data is feasible for the identification of genomic variants in a wide range of organisms, including livestock (73–76), and multiple pipelines have been developed for variant discovery and genotype calling (73–76). Here we opted for a hybrid approach, which utilized the “SplitNCigarReads” function of GATK (61) followed by the functions “mpileup” and “call” from BCFtools (64). The reason for using BCFtools was that it calls genotypes at every nucleotide position by default so that individuals were genotyped regardless of the homozygote or heterozygote makeup.

Prior research showed that the efficacy of genotype calling using RNA-sequencing data is high (77). Although we did not assess the specificity of genotype calling with an orthogonal method, we employed a stringent requirement for coverage equal to or greater than 20x, which is higher than the previously suggested 10x (74, 77) for high confidence genotype calling. In addition, 96% of the variants identified in our pipeline are present in the dbSNP ((67) version 150), and the variants have the same allelic composition reported in the dbSNP. Our hybrid pipeline efficiently genotyped individuals at homozygote and heterozygote genomic positions, although further confirmation is required for the variants called in our work that are not reported in the dbSNP.

### eQTL analysis using RNA-sequencing with and without forcing the data into a Gaussian distribution

Current statistical approaches employed for eQTL analysis (78) assume that the data is normally distributed, and the transformation of RNA-sequencing data to enforce a normal distribution is employed in nearly all major eQTL studies. Our comparison of the RNA-sequencing data prior to and after transforming the data (Supplementary Figure 1) does confirm that the inverse normal transformation (20) is highly effective in reducing skewness and shrinking the variance to reduce the impact of extreme values in the analysis (71), and thus making the data suitable for statistics tests requiring normally distributed data.

We first analyzed our data following the GTEx framework (1), transforming the data to achieve a normal distribution. Our analysis yielded less significant associations between genotype and gene transcript abundance relative to previously published studies that worked with genes expressed in blood samples (79–82) and the recent results from the cattle GTEx consortium (83). This large difference was expected because we only utilized 6,207 SNPs in our analysis, which yields less genotypic data as compared to high-throughput genotyping platforms or imputation of SNPs from reference populations. Another difference between our procedure and other reports was the stringent threshold to infer significance (P=5×10^-8^, -Log_10_(5×10^-8^)=7.3).

We noted, however, that visual inspection of the data with significant eQTLs identified with the ANOVA model (see examples in Figure 2C) does not clearly indicate patterns of data distribution that resemble the definition of allelic interaction characterized as complete dominance (84, 85). The dispersion of the data with significant eQTLs identified with the additive model (see examples in Figure 3C) does indicate patterns of data distribution that resemble alleles interacting in additive mode (84, 85). However, the distribution of heterozygotes showed two groups of samples with district profiles.

The graph profiles obtained from significant eQTLs using the GTEx framework prompted us to analyze the data using a DGE framework. To that end, we carried out an analysis using one of the commonly used statistical algorithms coded in the R package “edgeR” (21, 26, 86). The comparison of our eQTL analysis using “edgeR” showed a striking contrast with the analysis using the GTEx framework and “MatrixEQTL” in many important aspects. First, there was no overlap of significant eQTLs obtained between the two approaches within this study. Here, we point out that identifying which eQTL is true is virtually impossible without further mechanistic experiments that confirm the influence of allelic variants on gene expression (87, 88). Our findings add to previous observations that the type of statistical analysis carried out is a critical contributor to the lack of replicability observed across eQTL studies (89, 90). Second, working with specific contrasts, we were able to identify trans eQTLs that more closely resemble complete dominance, which were not identified by the standard framework. Our results are evidence that the number of genes whose expression are under genetic control and follow patterns of complete dominance (91, 92) is probably more common than previously expected (1). The identification of groups of genes enriched for specific biological processes strongly supports that this genetic control under the dominance model may have a biological role in the function of PWBCs.

We identified two important aspects that show a contrast between the ANOVA framework and the DEG framework we propose here. First, the functions in “MatrixEQTL” require less computational resources and time to conclude the analysis relative to the calculations carried out using the DGE framework in “edgeR”. Our proposed approach is inherently more complex, as we carried out multiple tests to provide robust and valuable information about dominance interaction between alleles. It is also very important to note that our study is not about the tools (“MatrixEQTL” or “edgeR”), because researchers can use other tools for the standard analysis of eQTL such as “FastQTL” (93) or DESeq2 (14) for the DGE framework. Second, the transformation of the data to fit a normal distribution clearly shrinks the variance (Supplementary Figure 3) reducing the differences in transcript abundance among genotypes thus reducing the likelihood of these eQTLs to be inferred as significant. At the end, the most critical choice researchers need to make is between (i) forcing data that is not normally distributed and has many outlier data points (70, 71) into normality or (ii) utilizing a framework that employs a statistical test appropriate for count data.

## Conclusions

In summary, different types of data normalization and analytical procedures lead to a variety of combinations that can be used for eQTL analysis using RNA-sequencing. Most of these approaches also transform the data to fit a normal distribution. Our analysis showed that it is possible to carry out eQTL studies using basic concepts and the analytical framework developed for differential gene expression that does not require data transformation to fit a normal distribution, thus it is likely more suitable for RNA-sequencing. The approach proposed here can uncover genetic control of gene expression that is biologically relevant for the tissue studied that otherwise may not be detected through data transformation and linear models.

## List of abbreviations

ANOVA: analysis of variance
CPM: counts per million reads
DGE: differential gene expression
eQTL: expression quantitative trait loci
FPKM: fragments per kilobase per million reads
FWER: family wise error rate
GTEx: genotype-tissue expression
Log_2_: Logarithm base 2
n: number
P: probability
PWBCs: peripheral white blood cells
RNA: ribonucleic acid
SNP: single nucleotide polymorphism
TMM: trimmed means of M-values
TPM: transcript per million reads
UTRs: untraslated regions

## Declarations

### Ethics approval and consent to participate

Not applicable

### Consent for publication

Not applicable

### Availability of data and material

The datasets generated and/or analysed during the current study are available in the GEO repository, session identifiers: GSE103628 and GSE146041.

### Author Contributions

Fernando Biase conceptualized the study and supervised the research. Fernando Biase and Mackenzie Marella developed the bioinformatic pipeline, wrote the code for analytical procedures, and wrote the paper.

## Acknowledgments

We appreciate the members of our team (Jada Nix and Chace Wilson) for their input and suggestions that improved our manuscript.

## Additional Files

File name: Additional file 1

File format: pdf

Title: Supplementary code to Robust identification of regulatory variants (eQTLs) using a differential expression framework developed for RNA-sequencing

Description: All codes utilized for in our work to produce the results described in the paper

File name: Additional file 2

File format: .xlsx

Title: Distribution of the SNPs based on presence in the database and predicted consequence of the variant.

Description: Distribution of the SNPs based on presence in the database and predicted consequence of the variant.

File name: Additional file 3

File format: .pdf

Title: Supplementary figures

Description: Supplementary figures

File name: Additional file 4

File format: .xlsx

Title: Annotated results of eQTL analysis using the GTEx framework and ANOVA model.

Description: Annotated results of eQTL analysis using the GTEx framework and ANOVA model.

File name: Additional file 5

File format: .xlsx

Title: Annotated results of eQTL analysis using the GTEx framework and additive model.

Description: Annotated results of eQTL analysis using the GTEx framework and additive model.

File name: Additional file 6

File format: .xlsx

Title: eQTL results with DGE framework and the standard approach using MatrixEQTL.

Description: eQTL results with DGE framework and the standard approach using MatrixEQTL.

File name: Additional file 7

File format: .xlsx

Title: Annotated eQTL results using edgeR framework and additive model.

Description: Annotated eQTL results using edgeR framework and additive model.

File name: Additional file 8

File format: .xlsx

Title: Annotated eQTL results with edgeR framework and ANOVA model.

Description: Annotated eQTL results with edgeR framework and ANOVA model.

File name: Additional file 9

File format: .xlsx

Title: Gene ontology analysis of genes whose expression are influenced by SNPs.

Description: Gene ontology analysis of genes whose expression are influenced by SNPs.

