## Additional file 1 for "Robust identification of regulatory variants (eQTLs) using a differential expression framework developed for RNA-sequencing"

Mackenzie Marrela and Fernando H. Biase

2022-11-17

#### Overview

Code produced by Mackenzie Marrela and Fernando Biase. We created this file to permit reproducibility of the findings described in the paper. Please direct questions to Fernando Biase: ***fbiase*** at ***vt.edu***

```
library('vcfR', lib.loc="/usr/lib/R/site-library")
library('R6', lib.loc="/usr/lib/R/site-library")
library('reshape2', lib.loc="/usr/lib/R/site-library")
library('stringr', lib.loc="/usr/lib/R/site-library")
library('qvalue', lib.loc="/usr/lib/R/site-library")
library('foreach', lib.loc="/usr/lib/R/site-library")
library('doParallel', lib.loc="/usr/lib/R/site-library")
library('parallel', lib.loc="/usr/lib/R/site-library")
library('Biobase', lib.loc="/usr/lib/R/site-library")
library('MatrixEQTL', lib.loc="/usr/lib/R/site-library")
library('GenomicTools', lib.loc="/usr/lib/R/site-library")
library("DESeq2", lib.loc="/usr/lib/R/site-library")
library('edgeR', lib.loc="/usr/lib/R/site-library")
library("ggplot2", lib.loc="/usr/lib/R/site-library")
library('sjmisc', lib.loc="/usr/lib/R/site-library")
library('readxl', lib.loc="/usr/lib/R/site-library")
library('cowplot', lib.loc="/usr/lib/R/site-library")
library("goseq" ,quietly = TRUE, lib.loc="/usr/lib/R/site-library")
library('ggforce',lib.loc="/usr/lib/R/site-library")
library('tidyr',lib.loc="/usr/lib/R/site-library")
library('dplyr',lib.loc="/usr/lib/R/site-library")
library('RNOmni',lib.loc="/usr/lib/R/site-library")
library('HardyWeinberg', lib.loc="/usr/lib/R/site-library")
library("doParallel",quietly = TRUE , lib.loc="/usr/lib/R/site-library")
library("grid",quietly = TRUE , lib.loc="/usr/lib/R/site-library")
library("gridExtra",quietly = TRUE , lib.loc="/usr/lib/R/site-library")
```

#read in files

```
snp_data_vcf_genotype<-foreach(i= 1:NROW(list_of_files),.combine='rbind',.inorder = FALSE,.packages = c('vcfR','R6','reshape2','stringr'))%dopar%{
  snp_data_vcf <- read.vcfR( list_of_files[i], verbose = FALSE)
  snp_data_vcf <- extract.indels(snp_data_vcf, return.indels = FALSE )
  snp_data_vcf <- vcfR2tidy(snp_data_vcf, info_only = FALSE, single_frame = TRUE, toss_INFO_column = TRUE)
  snp_data_vcf<-as.data.frame(snp_data_vcf$dat)
  snp_data_vcf$gt_GT <- ifelse(snp_data_vcf$gt_DP > 10, snp_data_vcf$gt_GT,  '<NA>' )
  snp_data_vcf$gt_GT_alleles <- ifelse(snp_data_vcf$gt_DP > 10, snp_data_vcf$gt_GT_alleles,  '<NA>' )
  snp_data_vcf$gt_GT <- ifelse(snp_data_vcf$gt_GT == "0/0" | snp_data_vcf$gt_GT == "0/1" | snp_data_vcf$gt_GT == "1/1" ,snp_data_vcf$gt_GT,  '<NA>')
  snp_data_vcf$gt_GT_alleles <- ifelse(snp_data_vcf$gt_GT == "0/0" | snp_data_vcf$gt_GT == "0/1" | snp_data_vcf$gt_GT == "1/1" ,snp_data_vcf$gt_GT_alleles,  '<NA>')
  snp_data_vcf_genotype<-reshape2::dcast( snp_data_vcf , CHROM + POS ~ Indiv, value.var="gt_GT")
  #snp_data_vcf_nucleotide<-reshape2::dcast( snp_data_vcf , CHROM + POS ~ Indiv, value.var="gt_GT_alleles")

  snp_data_vcf_genotype<-snp_data_vcf_genotype[rowSums(snp_data_vcf_genotype== "<NA>") < 26,]
  #snp_data_vcf_nucleotide<-snp_data_vcf_nucleotide[rowSums(snp_data_vcf_nucleotide== "<NA>") < 26,]

}

parallel::stopCluster(myCluster)



snp_data_vcf <- read.vcfR("/mnt/storage/lab_folder/heifer_infertility/alignment_SNP/2021_11_3_samtools_variant_filtering_chr29.vcf.gz", verbose = FALSE)
snp_data_vcf <- extract.indels(snp_data_vcf, return.indels = FALSE )
snp_data_vcf <- vcfR2tidy(snp_data_vcf, info_only = FALSE, single_frame = TRUE, toss_INFO_column = TRUE)
snp_data_vcf<-as.data.frame(snp_data_vcf$dat)
snp_data_vcf$gt_GT <- ifelse(snp_data_vcf$gt_DP > 10, snp_data_vcf$gt_GT,  '<NA>' )
snp_data_vcf$gt_GT_alleles <- ifelse(snp_data_vcf$gt_DP > 10, snp_data_vcf$gt_GT_alleles,  '<NA>' )
snp_data_vcf$gt_GT <- ifelse(snp_data_vcf$gt_GT == "0/0" | snp_data_vcf$gt_GT == "0/1" | snp_data_vcf$gt_GT == "1/1" ,snp_data_vcf$gt_GT,  '<NA>')
snp_data_vcf$gt_GT_alleles <- ifelse(snp_data_vcf$gt_GT == "0/0" | snp_data_vcf$gt_GT == "0/1" | snp_data_vcf$gt_GT == "1/1" ,snp_data_vcf$gt_GT_alleles,  '<NA>')
snp_data_vcf_genotype<-reshape2::dcast( snp_data_vcf , CHROM + POS ~ Indiv, value.var="gt_GT")
snp_data_vcf_nucleotide<-reshape2::dcast( snp_data_vcf , CHROM + POS ~ Indiv, value.var="gt_GT_alleles")

snp_data_vcf_genotype_chr29<-snp_data_vcf_genotype[rowSums(snp_data_vcf_genotype== "<NA>") < 26,]
snp_data_vcf_nucleotide_chr29<-snp_data_vcf_nucleotide[rowSums(snp_data_vcf_nucleotide== "<NA>") < 26,]
```

```
rm(snp_data_vcf,snp_data_vcf_genotype,snp_data_vcf_nucleotide)

snp_data_vcf <- read.vcfR("/mnt/storage/lab_folder/heifer_infertility/alignment_SNP/2021
_11_3_samtools_variant_filtering_chr1.vcf.gz", verbose = FALSE)
snp_data_vcf <- extract.indels(snp_data_vcf, return.indels = FALSE )
snp_data_vcf <- vcfR2tidy(snp_data_vcf, info_only = FALSE, single_frame = TRUE, toss_INF
O_column = TRUE)
snp_data_vcf<-as.data.frame(snp_data_vcf$dat)
snp_data_vcf$gt_GT <- ifelse(snp_data_vcf$gt_DP > 10, snp_data_vcf$gt_GT,  '<NA>' )
snp_data_vcf$gt_GT_alleles <- ifelse(snp_data_vcf$gt_DP > 10, snp_data_vcf$gt_GT_allele
s,  '<NA>' )
snp_data_vcf$gt_GT <- ifelse(snp_data_vcf$gt_GT == "0/0" | snp_data_vcf$gt_GT == "0/1" |
snp_data_vcf$gt_GT == "1/1" ,snp_data_vcf$gt_GT,  '<NA>')
snp_data_vcf$gt_GT_alleles <- ifelse(snp_data_vcf$gt_GT == "0/0" | snp_data_vcf$gt_GT ==
"0/1" | snp_data_vcf$gt_GT == "1/1" ,snp_data_vcf$gt_GT_alleles,  '<NA>')
snp_data_vcf_genotype<-reshape2::dcast( snp_data_vcf , CHROM + POS ~ Indiv, value.var="g
t_GT")
snp_data_vcf_nucleotide<-reshape2::dcast( snp_data_vcf , CHROM + POS ~ Indiv, value.var=
"gt_GT_alleles")

snp_data_vcf_genotype_chr1<-snp_data_vcf_genotype[rowSums(snp_data_vcf_genotype== "<NA>"
) < 26,]
snp_data_vcf_nucleotide_chr1<-snp_data_vcf_nucleotide[rowSums(snp_data_vcf_nucleotide==
"<NA>") < 26,]

rm(snp_data_vcf,snp_data_vcf_genotype,snp_data_vcf_nucleotide)


snp_data_vcf <- read.vcfR("/mnt/storage/lab_folder/heifer_infertility/alignment_SNP/2021
_11_3_samtools_variant_filtering_chr2.vcf.gz", verbose = FALSE)
snp_data_vcf <- extract.indels(snp_data_vcf, return.indels = FALSE )
snp_data_vcf <- vcfR2tidy(snp_data_vcf, info_only = FALSE, single_frame = TRUE, toss_INF
O_column = TRUE)
snp_data_vcf<-as.data.frame(snp_data_vcf$dat)
snp_data_vcf$gt_GT <- ifelse(snp_data_vcf$gt_DP > 10, snp_data_vcf$gt_GT,  '<NA>' )
snp_data_vcf$gt_GT_alleles <- ifelse(snp_data_vcf$gt_DP > 10, snp_data_vcf$gt_GT_allele
s,  '<NA>' )
snp_data_vcf$gt_GT <- ifelse(snp_data_vcf$gt_GT == "0/0" | snp_data_vcf$gt_GT == "0/1" |
snp_data_vcf$gt_GT == "1/1" ,snp_data_vcf$gt_GT,  '<NA>')
snp_data_vcf$gt_GT_alleles <- ifelse(snp_data_vcf$gt_GT == "0/0" | snp_data_vcf$gt_GT ==
"0/1" | snp_data_vcf$gt_GT == "1/1" ,snp_data_vcf$gt_GT_alleles,  '<NA>')
snp_data_vcf_genotype<-reshape2::dcast( snp_data_vcf , CHROM + POS ~ Indiv, value.var="g
t_GT")
snp_data_vcf_nucleotide<-reshape2::dcast( snp_data_vcf , CHROM + POS ~ Indiv, value.var=
"gt_GT_alleles")

snp_data_vcf_genotype_chr2<-snp_data_vcf_genotype[rowSums(snp_data_vcf_genotype== "<NA>"
) < 26,]
snp_data_vcf_nucleotide_chr2<-snp_data_vcf_nucleotide[rowSums(snp_data_vcf_nucleotide==
"<NA>") < 26,]
```

```
rm(snp_data_vcf,snp_data_vcf_genotype,snp_data_vcf_nucleotide)


snp_data_vcf <- read.vcfR("/mnt/storage/lab_folder/heifer_infertility/alignment_SNP/2021_11_3_samtools_variant_filtering_chr3.vcf.gz", verbose = FALSE)
snp_data_vcf <- extract.indels(snp_data_vcf, return.indels = FALSE )
snp_data_vcf <- vcfR2tidy(snp_data_vcf, info_only = FALSE, single_frame = TRUE, toss_INFO_column = TRUE)
snp_data_vcf<-as.data.frame(snp_data_vcf$dat)
snp_data_vcf$gt_GT <- ifelse(snp_data_vcf$gt_DP > 10, snp_data_vcf$gt_GT,  '<NA>' )
snp_data_vcf$gt_GT_alleles <- ifelse(snp_data_vcf$gt_DP > 10, snp_data_vcf$gt_GT_alleles,  '<NA>' )
snp_data_vcf$gt_GT <- ifelse(snp_data_vcf$gt_GT == "0/0" | snp_data_vcf$gt_GT == "0/1" | snp_data_vcf$gt_GT == "1/1" ,snp_data_vcf$gt_GT,  '<NA>')
snp_data_vcf$gt_GT_alleles <- ifelse(snp_data_vcf$gt_GT == "0/0" | snp_data_vcf$gt_GT == "0/1" | snp_data_vcf$gt_GT == "1/1" ,snp_data_vcf$gt_GT_alleles,  '<NA>')
snp_data_vcf_genotype<-reshape2::dcast( snp_data_vcf , CHROM + POS ~ Indiv, value.var="gt_GT")
snp_data_vcf_nucleotide<-reshape2::dcast( snp_data_vcf , CHROM + POS ~ Indiv, value.var="gt_GT_alleles")

snp_data_vcf_genotype_chr3<-snp_data_vcf_genotype[rowSums(snp_data_vcf_genotype== "<NA>") < 26,]
snp_data_vcf_nucleotide_chr3<-snp_data_vcf_nucleotide[rowSums(snp_data_vcf_nucleotide=="<NA>") < 26,]

rm(snp_data_vcf,snp_data_vcf_genotype,snp_data_vcf_nucleotide)

snp_data_vcf <- read.vcfR("/mnt/storage/lab_folder/heifer_infertility/alignment_SNP/2021_11_3_samtools_variant_filtering_chr4.vcf.gz", verbose = FALSE)
snp_data_vcf <- extract.indels(snp_data_vcf, return.indels = FALSE )
snp_data_vcf <- vcfR2tidy(snp_data_vcf, info_only = FALSE, single_frame = TRUE, toss_INFO_column = TRUE)
snp_data_vcf<-as.data.frame(snp_data_vcf$dat)
snp_data_vcf$gt_GT <- ifelse(snp_data_vcf$gt_DP > 10, snp_data_vcf$gt_GT,  '<NA>' )
snp_data_vcf$gt_GT_alleles <- ifelse(snp_data_vcf$gt_DP > 10, snp_data_vcf$gt_GT_alleles,  '<NA>' )
snp_data_vcf$gt_GT <- ifelse(snp_data_vcf$gt_GT == "0/0" | snp_data_vcf$gt_GT == "0/1" | snp_data_vcf$gt_GT == "1/1" ,snp_data_vcf$gt_GT,  '<NA>')
snp_data_vcf$gt_GT_alleles <- ifelse(snp_data_vcf$gt_GT == "0/0" | snp_data_vcf$gt_GT == "0/1" | snp_data_vcf$gt_GT == "1/1" ,snp_data_vcf$gt_GT_alleles,  '<NA>')
snp_data_vcf_genotype<-reshape2::dcast( snp_data_vcf , CHROM + POS ~ Indiv, value.var="gt_GT")
snp_data_vcf_nucleotide<-reshape2::dcast( snp_data_vcf , CHROM + POS ~ Indiv, value.var="gt_GT_alleles")

snp_data_vcf_genotype_chr4<-snp_data_vcf_genotype[rowSums(snp_data_vcf_genotype== "<NA>") < 26,]
snp_data_vcf_nucleotide_chr4<-snp_data_vcf_nucleotide[rowSums(snp_data_vcf_nucleotide=="<NA>") < 26,]

rm(snp_data_vcf,snp_data_vcf_genotype,snp_data_vcf_nucleotide)
```

```
snp_data_vcf <- read.vcfR("/mnt/storage/lab_folder/heifer_infertility/alignment_SNP/2021
_11_3_samtools_variant_filtering_chr5.vcf.gz", verbose = FALSE)
snp_data_vcf <- extract.indels(snp_data_vcf, return.indels = FALSE )
snp_data_vcf <- vcfR2tidy(snp_data_vcf, info_only = FALSE, single_frame = TRUE, toss_INF
O_column = TRUE)
snp_data_vcf<-as.data.frame(snp_data_vcf$dat)
snp_data_vcf$gt_GT <- ifelse(snp_data_vcf$gt_DP > 10, snp_data_vcf$gt_GT,  '<NA>' )
snp_data_vcf$gt_GT_alleles <- ifelse(snp_data_vcf$gt_DP > 10, snp_data_vcf$gt_GT_allele
s,  '<NA>' )
snp_data_vcf$gt_GT <- ifelse(snp_data_vcf$gt_GT == "0/0" | snp_data_vcf$gt_GT == "0/1" |
snp_data_vcf$gt_GT == "1/1" ,snp_data_vcf$gt_GT,  '<NA>')
snp_data_vcf$gt_GT_alleles <- ifelse(snp_data_vcf$gt_GT == "0/0" | snp_data_vcf$gt_GT ==
"0/1" | snp_data_vcf$gt_GT == "1/1" ,snp_data_vcf$gt_GT_alleles,  '<NA>')
snp_data_vcf_genotype<-reshape2::dcast( snp_data_vcf , CHROM + POS ~ Indiv, value.var="g
t_GT")
snp_data_vcf_nucleotide<-reshape2::dcast( snp_data_vcf , CHROM + POS ~ Indiv, value.var=
"gt_GT_alleles")

snp_data_vcf_genotype_chr5<-snp_data_vcf_genotype[rowSums(snp_data_vcf_genotype== "<NA>"
) < 26,]
snp_data_vcf_nucleotide_chr5<-snp_data_vcf_nucleotide[rowSums(snp_data_vcf_nucleotide==
"<NA>") < 26,]

rm(snp_data_vcf,snp_data_vcf_genotype,snp_data_vcf_nucleotide)

snp_data_vcf <- read.vcfR("/mnt/storage/lab_folder/heifer_infertility/alignment_SNP/2021
_11_3_samtools_variant_filtering_chr6.vcf.gz", verbose = FALSE)
snp_data_vcf <- extract.indels(snp_data_vcf, return.indels = FALSE )
snp_data_vcf <- vcfR2tidy(snp_data_vcf, info_only = FALSE, single_frame = TRUE, toss_INF
O_column = TRUE)
snp_data_vcf<-as.data.frame(snp_data_vcf$dat)
snp_data_vcf$gt_GT <- ifelse(snp_data_vcf$gt_DP > 10, snp_data_vcf$gt_GT,  '<NA>' )
snp_data_vcf$gt_GT_alleles <- ifelse(snp_data_vcf$gt_DP > 10, snp_data_vcf$gt_GT_allele
s,  '<NA>' )
snp_data_vcf$gt_GT <- ifelse(snp_data_vcf$gt_GT == "0/0" | snp_data_vcf$gt_GT == "0/1" |
snp_data_vcf$gt_GT == "1/1" ,snp_data_vcf$gt_GT,  '<NA>')
snp_data_vcf$gt_GT_alleles <- ifelse(snp_data_vcf$gt_GT == "0/0" | snp_data_vcf$gt_GT ==
"0/1" | snp_data_vcf$gt_GT == "1/1" ,snp_data_vcf$gt_GT_alleles,  '<NA>')
snp_data_vcf_genotype<-reshape2::dcast( snp_data_vcf , CHROM + POS ~ Indiv, value.var="g
t_GT")
snp_data_vcf_nucleotide<-reshape2::dcast( snp_data_vcf , CHROM + POS ~ Indiv, value.var=
"gt_GT_alleles")

snp_data_vcf_genotype_chr6<-snp_data_vcf_genotype[rowSums(snp_data_vcf_genotype== "<NA>"
) < 26,]
snp_data_vcf_nucleotide_chr6<-snp_data_vcf_nucleotide[rowSums(snp_data_vcf_nucleotide==
"<NA>") < 26,]

rm(snp_data_vcf,snp_data_vcf_genotype,snp_data_vcf_nucleotide)

snp_data_vcf <- read.vcfR("/mnt/storage/lab_folder/heifer_infertility/alignment_SNP/2021
```

```
_11_3_samtools_variant_filtering_chr7.vcf.gz", verbose = FALSE)
snp_data_vcf <- extract.indels(snp_data_vcf, return.indels = FALSE )
snp_data_vcf <- vcfR2tidy(snp_data_vcf, info_only = FALSE, single_frame = TRUE, toss_INF
O_column = TRUE)
snp_data_vcf<-as.data.frame(snp_data_vcf$dat)
snp_data_vcf$gt_GT <- ifelse(snp_data_vcf$gt_DP > 10, snp_data_vcf$gt_GT,  '<NA>' )
snp_data_vcf$gt_GT_alleles <- ifelse(snp_data_vcf$gt_DP > 10, snp_data_vcf$gt_GT_allele
s,  '<NA>' )
snp_data_vcf$gt_GT <- ifelse(snp_data_vcf$gt_GT == "0/0" | snp_data_vcf$gt_GT == "0/1" |
snp_data_vcf$gt_GT == "1/1" ,snp_data_vcf$gt_GT,  '<NA>')
snp_data_vcf$gt_GT_alleles <- ifelse(snp_data_vcf$gt_GT == "0/0" | snp_data_vcf$gt_GT ==
"0/1" | snp_data_vcf$gt_GT == "1/1" ,snp_data_vcf$gt_GT_alleles,  '<NA>')
snp_data_vcf_genotype<-reshape2::dcast( snp_data_vcf , CHROM + POS ~ Indiv, value.var="g
t_GT")
snp_data_vcf_nucleotide<-reshape2::dcast( snp_data_vcf , CHROM + POS ~ Indiv, value.var=
"gt_GT_alleles")

snp_data_vcf_genotype_chr7<-snp_data_vcf_genotype[rowSums(snp_data_vcf_genotype== "<NA>"
) < 26,]
snp_data_vcf_nucleotide_chr7<-snp_data_vcf_nucleotide[rowSums(snp_data_vcf_nucleotide==
"<NA>") < 26,]

rm(snp_data_vcf,snp_data_vcf_genotype,snp_data_vcf_nucleotide)

snp_data_vcf <- read.vcfR("/mnt/storage/lab_folder/heifer_infertility/alignment_SNP/2021
_11_3_samtools_variant_filtering_chr8.vcf.gz", verbose = FALSE)
snp_data_vcf <- extract.indels(snp_data_vcf, return.indels = FALSE )
snp_data_vcf <- vcfR2tidy(snp_data_vcf, info_only = FALSE, single_frame = TRUE, toss_INF
O_column = TRUE)
snp_data_vcf<-as.data.frame(snp_data_vcf$dat)
snp_data_vcf$gt_GT <- ifelse(snp_data_vcf$gt_DP > 10, snp_data_vcf$gt_GT,  '<NA>' )
snp_data_vcf$gt_GT_alleles <- ifelse(snp_data_vcf$gt_DP > 10, snp_data_vcf$gt_GT_allele
s,  '<NA>' )
snp_data_vcf$gt_GT <- ifelse(snp_data_vcf$gt_GT == "0/0" | snp_data_vcf$gt_GT == "0/1" |
snp_data_vcf$gt_GT == "1/1" ,snp_data_vcf$gt_GT,  '<NA>')
snp_data_vcf$gt_GT_alleles <- ifelse(snp_data_vcf$gt_GT == "0/0" | snp_data_vcf$gt_GT ==
"0/1" | snp_data_vcf$gt_GT == "1/1" ,snp_data_vcf$gt_GT_alleles,  '<NA>')
snp_data_vcf_genotype<-reshape2::dcast( snp_data_vcf , CHROM + POS ~ Indiv, value.var="g
t_GT")
snp_data_vcf_nucleotide<-reshape2::dcast( snp_data_vcf , CHROM + POS ~ Indiv, value.var=
"gt_GT_alleles")

snp_data_vcf_genotype_chr8<-snp_data_vcf_genotype[rowSums(snp_data_vcf_genotype== "<NA>"
) < 26,]
snp_data_vcf_nucleotide_chr8<-snp_data_vcf_nucleotide[rowSums(snp_data_vcf_nucleotide==
"<NA>") < 26,]

rm(snp_data_vcf,snp_data_vcf_genotype,snp_data_vcf_nucleotide)

snp_data_vcf <- read.vcfR("/mnt/storage/lab_folder/heifer_infertility/alignment_SNP/2021
_11_3_samtools_variant_filtering_chr9.vcf.gz", verbose = FALSE)
snp_data_vcf <- extract.indels(snp_data_vcf, return.indels = FALSE )
```

```
snp_data_vcf <- vcfR2tidy(snp_data_vcf, info_only = FALSE, single_frame = TRUE, toss_INF
O_column = TRUE)
snp_data_vcf<-as.data.frame(snp_data_vcf$dat)
snp_data_vcf$gt_GT <- ifelse(snp_data_vcf$gt_DP > 10, snp_data_vcf$gt_GT,  '<NA>' )
snp_data_vcf$gt_GT_alleles <- ifelse(snp_data_vcf$gt_DP > 10, snp_data_vcf$gt_GT_allele
s,  '<NA>' )
snp_data_vcf$gt_GT <- ifelse(snp_data_vcf$gt_GT == "0/0" | snp_data_vcf$gt_GT == "0/1" |
snp_data_vcf$gt_GT == "1/1" ,snp_data_vcf$gt_GT,  '<NA>')
snp_data_vcf$gt_GT_alleles <- ifelse(snp_data_vcf$gt_GT == "0/0" | snp_data_vcf$gt_GT ==
"0/1" | snp_data_vcf$gt_GT == "1/1" ,snp_data_vcf$gt_GT_alleles,  '<NA>')
snp_data_vcf_genotype<-reshape2::dcast( snp_data_vcf , CHROM + POS ~ Indiv, value.var="g
t_GT")
snp_data_vcf_nucleotide<-reshape2::dcast( snp_data_vcf , CHROM + POS ~ Indiv, value.var=
"gt_GT_alleles")

snp_data_vcf_genotype_chr9<-snp_data_vcf_genotype[rowSums(snp_data_vcf_genotype== "<NA>"
) < 26,]
snp_data_vcf_nucleotide_chr9<-snp_data_vcf_nucleotide[rowSums(snp_data_vcf_nucleotide==
"<NA>") < 26,]

rm(snp_data_vcf,snp_data_vcf_genotype,snp_data_vcf_nucleotide)

snp_data_vcf <- read.vcfR("/mnt/storage/lab_folder/heifer_infertility/alignment_SNP/2021
_11_3_samtools_variant_filtering_chr10.vcf.gz", verbose = FALSE)
snp_data_vcf <- extract.indels(snp_data_vcf, return.indels = FALSE )
snp_data_vcf <- vcfR2tidy(snp_data_vcf, info_only = FALSE, single_frame = TRUE, toss_INF
O_column = TRUE)
snp_data_vcf<-as.data.frame(snp_data_vcf$dat)
snp_data_vcf$gt_GT <- ifelse(snp_data_vcf$gt_DP > 10, snp_data_vcf$gt_GT,  '<NA>' )
snp_data_vcf$gt_GT_alleles <- ifelse(snp_data_vcf$gt_DP > 10, snp_data_vcf$gt_GT_allele
s,  '<NA>' )
snp_data_vcf$gt_GT <- ifelse(snp_data_vcf$gt_GT == "0/0" | snp_data_vcf$gt_GT == "0/1" |
snp_data_vcf$gt_GT == "1/1" ,snp_data_vcf$gt_GT,  '<NA>')
snp_data_vcf$gt_GT_alleles <- ifelse(snp_data_vcf$gt_GT == "0/0" | snp_data_vcf$gt_GT ==
"0/1" | snp_data_vcf$gt_GT == "1/1" ,snp_data_vcf$gt_GT_alleles,  '<NA>')
snp_data_vcf_genotype<-reshape2::dcast( snp_data_vcf , CHROM + POS ~ Indiv, value.var="g
t_GT")
snp_data_vcf_nucleotide<-reshape2::dcast( snp_data_vcf , CHROM + POS ~ Indiv, value.var=
"gt_GT_alleles")

snp_data_vcf_genotype_chr10<-snp_data_vcf_genotype[rowSums(snp_data_vcf_genotype== "<NA
>") < 26,]
snp_data_vcf_nucleotide_chr10<-snp_data_vcf_nucleotide[rowSums(snp_data_vcf_nucleotide==
"<NA>") < 26,]

rm(snp_data_vcf,snp_data_vcf_genotype,snp_data_vcf_nucleotide)

snp_data_vcf <- read.vcfR("/mnt/storage/lab_folder/heifer_infertility/alignment_SNP/2021
_11_3_samtools_variant_filtering_chr11.vcf.gz", verbose = FALSE)
snp_data_vcf <- extract.indels(snp_data_vcf, return.indels = FALSE )
snp_data_vcf <- vcfR2tidy(snp_data_vcf, info_only = FALSE, single_frame = TRUE, toss_INF
O_column = TRUE)
```

```
snp_data_vcf<-as.data.frame(snp_data_vcf$dat)
snp_data_vcf$gt_GT <- ifelse(snp_data_vcf$gt_DP > 10, snp_data_vcf$gt_GT,  '<NA>' )
snp_data_vcf$gt_GT_alleles <- ifelse(snp_data_vcf$gt_DP > 10, snp_data_vcf$gt_GT_allele
s,  '<NA>' )
snp_data_vcf$gt_GT <- ifelse(snp_data_vcf$gt_GT == "0/0" | snp_data_vcf$gt_GT == "0/1" |
snp_data_vcf$gt_GT == "1/1" ,snp_data_vcf$gt_GT,  '<NA>')
snp_data_vcf$gt_GT_alleles <- ifelse(snp_data_vcf$gt_GT == "0/0" | snp_data_vcf$gt_GT ==
"0/1" | snp_data_vcf$gt_GT == "1/1" ,snp_data_vcf$gt_GT_alleles,  '<NA>')
snp_data_vcf_genotype<-reshape2::dcast( snp_data_vcf , CHROM + POS ~ Indiv, value.var="g
t_GT")
snp_data_vcf_nucleotide<-reshape2::dcast( snp_data_vcf , CHROM + POS ~ Indiv, value.var=
"gt_GT_alleles")

snp_data_vcf_genotype_chr11<-snp_data_vcf_genotype[rowSums(snp_data_vcf_genotype== "<NA
>") < 26,]
snp_data_vcf_nucleotide_chr11<-snp_data_vcf_nucleotide[rowSums(snp_data_vcf_nucleotide==
"<NA>") < 26,]

rm(snp_data_vcf,snp_data_vcf_genotype,snp_data_vcf_nucleotide)

snp_data_vcf <- read.vcfR("/mnt/storage/lab_folder/heifer_infertility/alignment_SNP/2021
_11_3_samtools_variant_filtering_chr12.vcf.gz", verbose = FALSE)
snp_data_vcf <- extract.indels(snp_data_vcf, return.indels = FALSE )
snp_data_vcf <- vcfR2tidy(snp_data_vcf, info_only = FALSE, single_frame = TRUE, toss_INF
O_column = TRUE)
snp_data_vcf<-as.data.frame(snp_data_vcf$dat)
snp_data_vcf$gt_GT <- ifelse(snp_data_vcf$gt_DP > 10, snp_data_vcf$gt_GT,  '<NA>' )
snp_data_vcf$gt_GT_alleles <- ifelse(snp_data_vcf$gt_DP > 10, snp_data_vcf$gt_GT_allele
s,  '<NA>' )
snp_data_vcf$gt_GT <- ifelse(snp_data_vcf$gt_GT == "0/0" | snp_data_vcf$gt_GT == "0/1" |
snp_data_vcf$gt_GT == "1/1" ,snp_data_vcf$gt_GT,  '<NA>')
snp_data_vcf$gt_GT_alleles <- ifelse(snp_data_vcf$gt_GT == "0/0" | snp_data_vcf$gt_GT ==
"0/1" | snp_data_vcf$gt_GT == "1/1" ,snp_data_vcf$gt_GT_alleles,  '<NA>')
snp_data_vcf_genotype<-reshape2::dcast( snp_data_vcf , CHROM + POS ~ Indiv, value.var="g
t_GT")
snp_data_vcf_nucleotide<-reshape2::dcast( snp_data_vcf , CHROM + POS ~ Indiv, value.var=
"gt_GT_alleles")

snp_data_vcf_genotype_chr12<-snp_data_vcf_genotype[rowSums(snp_data_vcf_genotype== "<NA
>") < 26,]
snp_data_vcf_nucleotide_chr12<-snp_data_vcf_nucleotide[rowSums(snp_data_vcf_nucleotide==
"<NA>") < 26,]

rm(snp_data_vcf,snp_data_vcf_genotype,snp_data_vcf_nucleotide)

snp_data_vcf <- read.vcfR("/mnt/storage/lab_folder/heifer_infertility/alignment_SNP/2021
_11_3_samtools_variant_filtering_chr13.vcf.gz", verbose = FALSE)
snp_data_vcf <- extract.indels(snp_data_vcf, return.indels = FALSE )
snp_data_vcf <- vcfR2tidy(snp_data_vcf, info_only = FALSE, single_frame = TRUE, toss_INF
O_column = TRUE)
snp_data_vcf<-as.data.frame(snp_data_vcf$dat)
snp_data_vcf$gt_GT <- ifelse(snp_data_vcf$gt_DP > 10, snp_data_vcf$gt_GT,  '<NA>' )
```

```
snp_data_vcf$gt_GT_alleles <- ifelse(snp_data_vcf$gt_DP > 10, snp_data_vcf$gt_GT_allele
s,  '<NA>' )
snp_data_vcf$gt_GT <- ifelse(snp_data_vcf$gt_GT == "0/0" | snp_data_vcf$gt_GT == "0/1" |
snp_data_vcf$gt_GT == "1/1" ,snp_data_vcf$gt_GT,  '<NA>')
snp_data_vcf$gt_GT_alleles <- ifelse(snp_data_vcf$gt_GT == "0/0" | snp_data_vcf$gt_GT ==
"0/1" | snp_data_vcf$gt_GT == "1/1" ,snp_data_vcf$gt_GT_alleles,  '<NA>')
snp_data_vcf_genotype<-reshape2::dcast( snp_data_vcf , CHROM + POS ~ Indiv, value.var="g
t_GT")
snp_data_vcf_nucleotide<-reshape2::dcast( snp_data_vcf , CHROM + POS ~ Indiv, value.var=
"gt_GT_alleles")

snp_data_vcf_genotype_chr13<-snp_data_vcf_genotype[rowSums(snp_data_vcf_genotype== "<NA
>") < 26,]
snp_data_vcf_nucleotide_chr13<-snp_data_vcf_nucleotide[rowSums(snp_data_vcf_nucleotide==
"<NA>") < 26,]

rm(snp_data_vcf,snp_data_vcf_genotype,snp_data_vcf_nucleotide)

snp_data_vcf <- read.vcfR("/mnt/storage/lab_folder/heifer_infertility/alignment_SNP/2021
_11_3_samtools_variant_filtering_chr14.vcf.gz", verbose = FALSE)
snp_data_vcf <- extract.indels(snp_data_vcf, return.indels = FALSE )
snp_data_vcf <- vcfR2tidy(snp_data_vcf, info_only = FALSE, single_frame = TRUE, toss_INF
O_column = TRUE)
snp_data_vcf<-as.data.frame(snp_data_vcf$dat)
snp_data_vcf$gt_GT <- ifelse(snp_data_vcf$gt_DP > 10, snp_data_vcf$gt_GT,  '<NA>' )
snp_data_vcf$gt_GT_alleles <- ifelse(snp_data_vcf$gt_DP > 10, snp_data_vcf$gt_GT_allele
s,  '<NA>' )
snp_data_vcf$gt_GT <- ifelse(snp_data_vcf$gt_GT == "0/0" | snp_data_vcf$gt_GT == "0/1" |
snp_data_vcf$gt_GT == "1/1" ,snp_data_vcf$gt_GT,  '<NA>')
snp_data_vcf$gt_GT_alleles <- ifelse(snp_data_vcf$gt_GT == "0/0" | snp_data_vcf$gt_GT ==
"0/1" | snp_data_vcf$gt_GT == "1/1" ,snp_data_vcf$gt_GT_alleles,  '<NA>')
snp_data_vcf_genotype<-reshape2::dcast( snp_data_vcf , CHROM + POS ~ Indiv, value.var="g
t_GT")
snp_data_vcf_nucleotide<-reshape2::dcast( snp_data_vcf , CHROM + POS ~ Indiv, value.var=
"gt_GT_alleles")

snp_data_vcf_genotype_chr14<-snp_data_vcf_genotype[rowSums(snp_data_vcf_genotype== "<NA
>") < 26,]
snp_data_vcf_nucleotide_chr14<-snp_data_vcf_nucleotide[rowSums(snp_data_vcf_nucleotide==
"<NA>") < 26,]

rm(snp_data_vcf,snp_data_vcf_genotype,snp_data_vcf_nucleotide)

snp_data_vcf <- read.vcfR("/mnt/storage/lab_folder/heifer_infertility/alignment_SNP/2021
_11_3_samtools_variant_filtering_chr15.vcf.gz", verbose = FALSE)
snp_data_vcf <- extract.indels(snp_data_vcf, return.indels = FALSE )
snp_data_vcf <- vcfR2tidy(snp_data_vcf, info_only = FALSE, single_frame = TRUE, toss_INF
O_column = TRUE)
snp_data_vcf<-as.data.frame(snp_data_vcf$dat)
snp_data_vcf$gt_GT <- ifelse(snp_data_vcf$gt_DP > 10, snp_data_vcf$gt_GT,  '<NA>' )
snp_data_vcf$gt_GT_alleles <- ifelse(snp_data_vcf$gt_DP > 10, snp_data_vcf$gt_GT_allele
s,  '<NA>' )
```

```
snp_data_vcf$gt_GT <- ifelse(snp_data_vcf$gt_GT == "0/0" | snp_data_vcf$gt_GT == "0/1" |
snp_data_vcf$gt_GT == "1/1" ,snp_data_vcf$gt_GT,  '<NA>')
snp_data_vcf$gt_GT_alleles <- ifelse(snp_data_vcf$gt_GT == "0/0" | snp_data_vcf$gt_GT ==
"0/1" | snp_data_vcf$gt_GT == "1/1" ,snp_data_vcf$gt_GT_alleles,  '<NA>')
snp_data_vcf_genotype<-reshape2::dcast( snp_data_vcf , CHROM + POS ~ Indiv, value.var="g
t_GT")
snp_data_vcf_nucleotide<-reshape2::dcast( snp_data_vcf , CHROM + POS ~ Indiv, value.var=
"gt_GT_alleles")

snp_data_vcf_genotype_chr15<-snp_data_vcf_genotype[rowSums(snp_data_vcf_genotype== "<NA
>") < 26,]
snp_data_vcf_nucleotide_chr15<-snp_data_vcf_nucleotide[rowSums(snp_data_vcf_nucleotide==
"<NA>") < 26,]

rm(snp_data_vcf,snp_data_vcf_genotype,snp_data_vcf_nucleotide)

snp_data_vcf <- read.vcfR("/mnt/storage/lab_folder/heifer_infertility/alignment_SNP/2021
_11_3_samtools_variant_filtering_chr16.vcf.gz", verbose = FALSE)
snp_data_vcf <- extract.indels(snp_data_vcf, return.indels = FALSE )
snp_data_vcf <- vcfR2tidy(snp_data_vcf, info_only = FALSE, single_frame = TRUE, toss_INF
O_column = TRUE)
snp_data_vcf<-as.data.frame(snp_data_vcf$dat)
snp_data_vcf$gt_GT <- ifelse(snp_data_vcf$gt_DP > 10, snp_data_vcf$gt_GT,  '<NA>' )
snp_data_vcf$gt_GT_alleles <- ifelse(snp_data_vcf$gt_DP > 10, snp_data_vcf$gt_GT_allele
s,  '<NA>' )
snp_data_vcf$gt_GT <- ifelse(snp_data_vcf$gt_GT == "0/0" | snp_data_vcf$gt_GT == "0/1" |
snp_data_vcf$gt_GT == "1/1" ,snp_data_vcf$gt_GT,  '<NA>')
snp_data_vcf$gt_GT_alleles <- ifelse(snp_data_vcf$gt_GT == "0/0" | snp_data_vcf$gt_GT ==
"0/1" | snp_data_vcf$gt_GT == "1/1" ,snp_data_vcf$gt_GT_alleles,  '<NA>')
snp_data_vcf_genotype<-reshape2::dcast( snp_data_vcf , CHROM + POS ~ Indiv, value.var="g
t_GT")
snp_data_vcf_nucleotide<-reshape2::dcast( snp_data_vcf , CHROM + POS ~ Indiv, value.var=
"gt_GT_alleles")

snp_data_vcf_genotype_chr16<-snp_data_vcf_genotype[rowSums(snp_data_vcf_genotype== "<NA
>") < 26,]
snp_data_vcf_nucleotide_chr16<-snp_data_vcf_nucleotide[rowSums(snp_data_vcf_nucleotide==
"<NA>") < 26,]

rm(snp_data_vcf,snp_data_vcf_genotype,snp_data_vcf_nucleotide)

snp_data_vcf <- read.vcfR("/mnt/storage/lab_folder/heifer_infertility/alignment_SNP/2021
_11_3_samtools_variant_filtering_chr17.vcf.gz", verbose = FALSE)
snp_data_vcf <- extract.indels(snp_data_vcf, return.indels = FALSE )
snp_data_vcf <- vcfR2tidy(snp_data_vcf, info_only = FALSE, single_frame = TRUE, toss_INF
O_column = TRUE)
snp_data_vcf<-as.data.frame(snp_data_vcf$dat)
snp_data_vcf$gt_GT <- ifelse(snp_data_vcf$gt_DP > 10, snp_data_vcf$gt_GT,  '<NA>' )
snp_data_vcf$gt_GT_alleles <- ifelse(snp_data_vcf$gt_DP > 10, snp_data_vcf$gt_GT_allele
s,  '<NA>' )
snp_data_vcf$gt_GT <- ifelse(snp_data_vcf$gt_GT == "0/0" | snp_data_vcf$gt_GT == "0/1" |
snp_data_vcf$gt_GT == "1/1" ,snp_data_vcf$gt_GT,  '<NA>')
```

```
snp_data_vcf$gt_GT_alleles <- ifelse(snp_data_vcf$gt_GT == "0/0" | snp_data_vcf$gt_GT ==
"0/1" | snp_data_vcf$gt_GT == "1/1" ,snp_data_vcf$gt_GT_alleles,  '<NA>')
snp_data_vcf_genotype<-reshape2::dcast( snp_data_vcf , CHROM + POS ~ Indiv, value.var="g
t_GT")
snp_data_vcf_nucleotide<-reshape2::dcast( snp_data_vcf , CHROM + POS ~ Indiv, value.var=
"gt_GT_alleles")

snp_data_vcf_genotype_chr17<-snp_data_vcf_genotype[rowSums(snp_data_vcf_genotype== "<NA
>") < 26,]
snp_data_vcf_nucleotide_chr17<-snp_data_vcf_nucleotide[rowSums(snp_data_vcf_nucleotide==
"<NA>") < 26,]

rm(snp_data_vcf,snp_data_vcf_genotype,snp_data_vcf_nucleotide)

snp_data_vcf <- read.vcfR("/mnt/storage/lab_folder/heifer_infertility/alignment_SNP/2021
_11_3_samtools_variant_filtering_chr18.vcf.gz", verbose = FALSE)
snp_data_vcf <- extract.indels(snp_data_vcf, return.indels = FALSE )
snp_data_vcf <- vcfR2tidy(snp_data_vcf, info_only = FALSE, single_frame = TRUE, toss_INF
O_column = TRUE)
snp_data_vcf<-as.data.frame(snp_data_vcf$dat)
snp_data_vcf$gt_GT <- ifelse(snp_data_vcf$gt_DP > 10, snp_data_vcf$gt_GT,  '<NA>' )
snp_data_vcf$gt_GT_alleles <- ifelse(snp_data_vcf$gt_DP > 10, snp_data_vcf$gt_GT_allele
s,  '<NA>' )
snp_data_vcf$gt_GT <- ifelse(snp_data_vcf$gt_GT == "0/0" | snp_data_vcf$gt_GT == "0/1" |
snp_data_vcf$gt_GT == "1/1" ,snp_data_vcf$gt_GT,  '<NA>')
snp_data_vcf$gt_GT_alleles <- ifelse(snp_data_vcf$gt_GT == "0/0" | snp_data_vcf$gt_GT ==
"0/1" | snp_data_vcf$gt_GT == "1/1" ,snp_data_vcf$gt_GT_alleles,  '<NA>')
snp_data_vcf_genotype<-reshape2::dcast( snp_data_vcf , CHROM + POS ~ Indiv, value.var="g
t_GT")
snp_data_vcf_nucleotide<-reshape2::dcast( snp_data_vcf , CHROM + POS ~ Indiv, value.var=
"gt_GT_alleles")

snp_data_vcf_genotype_chr18<-snp_data_vcf_genotype[rowSums(snp_data_vcf_genotype== "<NA
>") < 26,]
snp_data_vcf_nucleotide_chr18<-snp_data_vcf_nucleotide[rowSums(snp_data_vcf_nucleotide==
"<NA>") < 26,]

rm(snp_data_vcf,snp_data_vcf_genotype,snp_data_vcf_nucleotide)

snp_data_vcf <- read.vcfR("/mnt/storage/lab_folder/heifer_infertility/alignment_SNP/2021
_11_3_samtools_variant_filtering_chr19.vcf.gz", verbose = FALSE)
snp_data_vcf <- extract.indels(snp_data_vcf, return.indels = FALSE )
snp_data_vcf <- vcfR2tidy(snp_data_vcf, info_only = FALSE, single_frame = TRUE, toss_INF
O_column = TRUE)
snp_data_vcf<-as.data.frame(snp_data_vcf$dat)
snp_data_vcf$gt_GT <- ifelse(snp_data_vcf$gt_DP > 10, snp_data_vcf$gt_GT,  '<NA>' )
snp_data_vcf$gt_GT_alleles <- ifelse(snp_data_vcf$gt_DP > 10, snp_data_vcf$gt_GT_allele
s,  '<NA>' )
snp_data_vcf$gt_GT <- ifelse(snp_data_vcf$gt_GT == "0/0" | snp_data_vcf$gt_GT == "0/1" |
snp_data_vcf$gt_GT == "1/1" ,snp_data_vcf$gt_GT,  '<NA>')
snp_data_vcf$gt_GT_alleles <- ifelse(snp_data_vcf$gt_GT == "0/0" | snp_data_vcf$gt_GT ==
"0/1" | snp_data_vcf$gt_GT == "1/1" ,snp_data_vcf$gt_GT_alleles,  '<NA>')
```

```
snp_data_vcf_genotype<-reshape2::dcast( snp_data_vcf , CHROM + POS ~ Indiv, value.var="g
t_GT")
snp_data_vcf_nucleotide<-reshape2::dcast( snp_data_vcf , CHROM + POS ~ Indiv, value.var=
"gt_GT_alleles")

snp_data_vcf_genotype_chr19<-snp_data_vcf_genotype[rowSums(snp_data_vcf_genotype== "<NA
>") < 26,]
snp_data_vcf_nucleotide_chr19<-snp_data_vcf_nucleotide[rowSums(snp_data_vcf_nucleotide==
"<NA>") < 26,]

rm(snp_data_vcf,snp_data_vcf_genotype,snp_data_vcf_nucleotide)

snp_data_vcf <- read.vcfR("/mnt/storage/lab_folder/heifer_infertility/alignment_SNP/2021
_11_3_samtools_variant_filtering_chr20.vcf.gz", verbose = FALSE)
snp_data_vcf <- extract.indels(snp_data_vcf, return.indels = FALSE )
snp_data_vcf <- vcfR2tidy(snp_data_vcf, info_only = FALSE, single_frame = TRUE, toss_INF
O_column = TRUE)
snp_data_vcf<-as.data.frame(snp_data_vcf$dat)
snp_data_vcf$gt_GT <- ifelse(snp_data_vcf$gt_DP > 10, snp_data_vcf$gt_GT,  '<NA>' )
snp_data_vcf$gt_GT_alleles <- ifelse(snp_data_vcf$gt_DP > 10, snp_data_vcf$gt_GT_allele
s,  '<NA>' )
snp_data_vcf$gt_GT <- ifelse(snp_data_vcf$gt_GT == "0/0" | snp_data_vcf$gt_GT == "0/1" |
snp_data_vcf$gt_GT == "1/1" ,snp_data_vcf$gt_GT,  '<NA>')
snp_data_vcf$gt_GT_alleles <- ifelse(snp_data_vcf$gt_GT == "0/0" | snp_data_vcf$gt_GT ==
"0/1" | snp_data_vcf$gt_GT == "1/1" ,snp_data_vcf$gt_GT_alleles,  '<NA>')
snp_data_vcf_genotype<-reshape2::dcast( snp_data_vcf , CHROM + POS ~ Indiv, value.var="g
t_GT")
snp_data_vcf_nucleotide<-reshape2::dcast( snp_data_vcf , CHROM + POS ~ Indiv, value.var=
"gt_GT_alleles")

snp_data_vcf_genotype_chr20<-snp_data_vcf_genotype[rowSums(snp_data_vcf_genotype== "<NA
>") < 26,]
snp_data_vcf_nucleotide_chr20<-snp_data_vcf_nucleotide[rowSums(snp_data_vcf_nucleotide==
"<NA>") < 26,]

rm(snp_data_vcf,snp_data_vcf_genotype,snp_data_vcf_nucleotide)

snp_data_vcf <- read.vcfR("/mnt/storage/lab_folder/heifer_infertility/alignment_SNP/2021
_11_3_samtools_variant_filtering_chr21.vcf.gz", verbose = FALSE)
snp_data_vcf <- extract.indels(snp_data_vcf, return.indels = FALSE )
snp_data_vcf <- vcfR2tidy(snp_data_vcf, info_only = FALSE, single_frame = TRUE, toss_INF
O_column = TRUE)
snp_data_vcf<-as.data.frame(snp_data_vcf$dat)
snp_data_vcf$gt_GT <- ifelse(snp_data_vcf$gt_DP > 10, snp_data_vcf$gt_GT,  '<NA>' )
snp_data_vcf$gt_GT_alleles <- ifelse(snp_data_vcf$gt_DP > 10, snp_data_vcf$gt_GT_allele
s,  '<NA>' )
snp_data_vcf$gt_GT <- ifelse(snp_data_vcf$gt_GT == "0/0" | snp_data_vcf$gt_GT == "0/1" |
snp_data_vcf$gt_GT == "1/1" ,snp_data_vcf$gt_GT,  '<NA>')
snp_data_vcf$gt_GT_alleles <- ifelse(snp_data_vcf$gt_GT == "0/0" | snp_data_vcf$gt_GT ==
"0/1" | snp_data_vcf$gt_GT == "1/1" ,snp_data_vcf$gt_GT_alleles,  '<NA>')
snp_data_vcf_genotype<-reshape2::dcast( snp_data_vcf , CHROM + POS ~ Indiv, value.var="g
t_GT")
```

```
snp_data_vcf_nucleotide<-reshape2::dcast( snp_data_vcf , CHROM + POS ~ Indiv, value.var=
"gt_GT_alleles")

snp_data_vcf_genotype_chr21<-snp_data_vcf_genotype[rowSums(snp_data_vcf_genotype== "<NA
>") < 26,]
snp_data_vcf_nucleotide_chr21<-snp_data_vcf_nucleotide[rowSums(snp_data_vcf_nucleotide==
"<NA>") < 26,]

rm(snp_data_vcf,snp_data_vcf_genotype,snp_data_vcf_nucleotide)

snp_data_vcf <- read.vcfR("/mnt/storage/lab_folder/heifer_infertility/alignment_SNP/2021
_11_3_samtools_variant_filtering_chr22.vcf.gz", verbose = FALSE)
snp_data_vcf <- extract.indels(snp_data_vcf, return.indels = FALSE )
snp_data_vcf <- vcfR2tidy(snp_data_vcf, info_only = FALSE, single_frame = TRUE, toss_INF
O_column = TRUE)
snp_data_vcf<-as.data.frame(snp_data_vcf$dat)
snp_data_vcf$gt_GT <- ifelse(snp_data_vcf$gt_DP > 10, snp_data_vcf$gt_GT,  '<NA>' )
snp_data_vcf$gt_GT_alleles <- ifelse(snp_data_vcf$gt_DP > 10, snp_data_vcf$gt_GT_allele
s,  '<NA>' )
snp_data_vcf$gt_GT <- ifelse(snp_data_vcf$gt_GT == "0/0" | snp_data_vcf$gt_GT == "0/1" |
snp_data_vcf$gt_GT == "1/1" ,snp_data_vcf$gt_GT,  '<NA>')
snp_data_vcf$gt_GT_alleles <- ifelse(snp_data_vcf$gt_GT == "0/0" | snp_data_vcf$gt_GT ==
"0/1" | snp_data_vcf$gt_GT == "1/1" ,snp_data_vcf$gt_GT_alleles,  '<NA>')
snp_data_vcf_genotype<-reshape2::dcast( snp_data_vcf , CHROM + POS ~ Indiv, value.var="g
t_GT")
snp_data_vcf_nucleotide<-reshape2::dcast( snp_data_vcf , CHROM + POS ~ Indiv, value.var=
"gt_GT_alleles")

snp_data_vcf_genotype_chr22<-snp_data_vcf_genotype[rowSums(snp_data_vcf_genotype== "<NA
>") < 26,]
snp_data_vcf_nucleotide_chr22<-snp_data_vcf_nucleotide[rowSums(snp_data_vcf_nucleotide==
"<NA>") < 26,]

rm(snp_data_vcf,snp_data_vcf_genotype,snp_data_vcf_nucleotide)

snp_data_vcf <- read.vcfR("/mnt/storage/lab_folder/heifer_infertility/alignment_SNP/2021
_11_3_samtools_variant_filtering_chr23.vcf.gz", verbose = FALSE)
snp_data_vcf <- extract.indels(snp_data_vcf, return.indels = FALSE )
snp_data_vcf <- vcfR2tidy(snp_data_vcf, info_only = FALSE, single_frame = TRUE, toss_INF
O_column = TRUE)
snp_data_vcf<-as.data.frame(snp_data_vcf$dat)
snp_data_vcf$gt_GT <- ifelse(snp_data_vcf$gt_DP > 10, snp_data_vcf$gt_GT,  '<NA>' )
snp_data_vcf$gt_GT_alleles <- ifelse(snp_data_vcf$gt_DP > 10, snp_data_vcf$gt_GT_allele
s,  '<NA>' )
snp_data_vcf$gt_GT <- ifelse(snp_data_vcf$gt_GT == "0/0" | snp_data_vcf$gt_GT == "0/1" |
snp_data_vcf$gt_GT == "1/1" ,snp_data_vcf$gt_GT,  '<NA>')
snp_data_vcf$gt_GT_alleles <- ifelse(snp_data_vcf$gt_GT == "0/0" | snp_data_vcf$gt_GT ==
"0/1" | snp_data_vcf$gt_GT == "1/1" ,snp_data_vcf$gt_GT_alleles,  '<NA>')
snp_data_vcf_genotype<-reshape2::dcast( snp_data_vcf , CHROM + POS ~ Indiv, value.var="g
t_GT")
snp_data_vcf_nucleotide<-reshape2::dcast( snp_data_vcf , CHROM + POS ~ Indiv, value.var=
"gt_GT_alleles")
```

```
snp_data_vcf_genotype_chr23<-snp_data_vcf_genotype[rowSums(snp_data_vcf_genotype== "<NA
>") < 26,]
snp_data_vcf_nucleotide_chr23<-snp_data_vcf_nucleotide[rowSums(snp_data_vcf_nucleotide==
"<NA>") < 26,]

rm(snp_data_vcf,snp_data_vcf_genotype,snp_data_vcf_nucleotide)

snp_data_vcf <- read.vcfR("/mnt/storage/lab_folder/heifer_infertility/alignment_SNP/2021
_11_3_samtools_variant_filtering_chr24.vcf.gz", verbose = FALSE)
snp_data_vcf <- extract.indels(snp_data_vcf, return.indels = FALSE )
snp_data_vcf <- vcfR2tidy(snp_data_vcf, info_only = FALSE, single_frame = TRUE, toss_INF
O_column = TRUE)
snp_data_vcf<-as.data.frame(snp_data_vcf$dat)
snp_data_vcf$gt_GT <- ifelse(snp_data_vcf$gt_DP > 10, snp_data_vcf$gt_GT,  '<NA>' )
snp_data_vcf$gt_GT_alleles <- ifelse(snp_data_vcf$gt_DP > 10, snp_data_vcf$gt_GT_allele
s,  '<NA>' )
snp_data_vcf$gt_GT <- ifelse(snp_data_vcf$gt_GT == "0/0" | snp_data_vcf$gt_GT == "0/1" |
snp_data_vcf$gt_GT == "1/1" ,snp_data_vcf$gt_GT,  '<NA>')
snp_data_vcf$gt_GT_alleles <- ifelse(snp_data_vcf$gt_GT == "0/0" | snp_data_vcf$gt_GT ==
"0/1" | snp_data_vcf$gt_GT == "1/1" ,snp_data_vcf$gt_GT_alleles,  '<NA>')
snp_data_vcf_genotype<-reshape2::dcast( snp_data_vcf , CHROM + POS ~ Indiv, value.var="g
t_GT")
snp_data_vcf_nucleotide<-reshape2::dcast( snp_data_vcf , CHROM + POS ~ Indiv, value.var=
"gt_GT_alleles")

snp_data_vcf_genotype_chr24<-snp_data_vcf_genotype[rowSums(snp_data_vcf_genotype== "<NA
>") < 26,]
snp_data_vcf_nucleotide_chr24<-snp_data_vcf_nucleotide[rowSums(snp_data_vcf_nucleotide==
"<NA>") < 26,]

rm(snp_data_vcf,snp_data_vcf_genotype,snp_data_vcf_nucleotide)

snp_data_vcf <- read.vcfR("/mnt/storage/lab_folder/heifer_infertility/alignment_SNP/2021
_11_3_samtools_variant_filtering_chr25.vcf.gz", verbose = FALSE)
snp_data_vcf <- extract.indels(snp_data_vcf, return.indels = FALSE )
snp_data_vcf <- vcfR2tidy(snp_data_vcf, info_only = FALSE, single_frame = TRUE, toss_INF
O_column = TRUE)
snp_data_vcf<-as.data.frame(snp_data_vcf$dat)
snp_data_vcf$gt_GT <- ifelse(snp_data_vcf$gt_DP > 10, snp_data_vcf$gt_GT,  '<NA>' )
snp_data_vcf$gt_GT_alleles <- ifelse(snp_data_vcf$gt_DP > 10, snp_data_vcf$gt_GT_allele
s,  '<NA>' )
snp_data_vcf$gt_GT <- ifelse(snp_data_vcf$gt_GT == "0/0" | snp_data_vcf$gt_GT == "0/1" |
snp_data_vcf$gt_GT == "1/1" ,snp_data_vcf$gt_GT,  '<NA>')
snp_data_vcf$gt_GT_alleles <- ifelse(snp_data_vcf$gt_GT == "0/0" | snp_data_vcf$gt_GT ==
"0/1" | snp_data_vcf$gt_GT == "1/1" ,snp_data_vcf$gt_GT_alleles,  '<NA>')
snp_data_vcf_genotype<-reshape2::dcast( snp_data_vcf , CHROM + POS ~ Indiv, value.var="g
t_GT")
snp_data_vcf_nucleotide<-reshape2::dcast( snp_data_vcf , CHROM + POS ~ Indiv, value.var=
"gt_GT_alleles")

snp_data_vcf_genotype_chr25<-snp_data_vcf_genotype[rowSums(snp_data_vcf_genotype== "<NA
```

```
>") < 26,]
snp_data_vcf_nucleotide_chr25<-snp_data_vcf_nucleotide[rowSums(snp_data_vcf_nucleotide==
"<NA>") < 26,]

rm(snp_data_vcf,snp_data_vcf_genotype,snp_data_vcf_nucleotide)

snp_data_vcf <- read.vcfR("/mnt/storage/lab_folder/heifer_infertility/alignment_SNP/2021
_11_3_samtools_variant_filtering_chr26.vcf.gz", verbose = FALSE)
snp_data_vcf <- extract.indels(snp_data_vcf, return.indels = FALSE )
snp_data_vcf <- vcfR2tidy(snp_data_vcf, info_only = FALSE, single_frame = TRUE, toss_INF
O_column = TRUE)
snp_data_vcf<-as.data.frame(snp_data_vcf$dat)
snp_data_vcf$gt_GT <- ifelse(snp_data_vcf$gt_DP > 10, snp_data_vcf$gt_GT,  '<NA>' )
snp_data_vcf$gt_GT_alleles <- ifelse(snp_data_vcf$gt_DP > 10, snp_data_vcf$gt_GT_allele
s,  '<NA>' )
snp_data_vcf$gt_GT <- ifelse(snp_data_vcf$gt_GT == "0/0" | snp_data_vcf$gt_GT == "0/1" |
snp_data_vcf$gt_GT == "1/1" ,snp_data_vcf$gt_GT,  '<NA>')
snp_data_vcf$gt_GT_alleles <- ifelse(snp_data_vcf$gt_GT == "0/0" | snp_data_vcf$gt_GT ==
"0/1" | snp_data_vcf$gt_GT == "1/1" ,snp_data_vcf$gt_GT_alleles,  '<NA>')
snp_data_vcf_genotype<-reshape2::dcast( snp_data_vcf , CHROM + POS ~ Indiv, value.var="g
t_GT")
snp_data_vcf_nucleotide<-reshape2::dcast( snp_data_vcf , CHROM + POS ~ Indiv, value.var=
"gt_GT_alleles")

snp_data_vcf_genotype_chr26<-snp_data_vcf_genotype[rowSums(snp_data_vcf_genotype== "<NA
>") < 26,]
snp_data_vcf_nucleotide_chr26<-snp_data_vcf_nucleotide[rowSums(snp_data_vcf_nucleotide==
"<NA>") < 26,]

rm(snp_data_vcf,snp_data_vcf_genotype,snp_data_vcf_nucleotide)

snp_data_vcf <- read.vcfR("/mnt/storage/lab_folder/heifer_infertility/alignment_SNP/2021
_11_3_samtools_variant_filtering_chr27.vcf.gz", verbose = FALSE)
snp_data_vcf <- extract.indels(snp_data_vcf, return.indels = FALSE )
snp_data_vcf <- vcfR2tidy(snp_data_vcf, info_only = FALSE, single_frame = TRUE, toss_INF
O_column = TRUE)
snp_data_vcf<-as.data.frame(snp_data_vcf$dat)
snp_data_vcf$gt_GT <- ifelse(snp_data_vcf$gt_DP > 10, snp_data_vcf$gt_GT,  '<NA>' )
snp_data_vcf$gt_GT_alleles <- ifelse(snp_data_vcf$gt_DP > 10, snp_data_vcf$gt_GT_allele
s,  '<NA>' )
snp_data_vcf$gt_GT <- ifelse(snp_data_vcf$gt_GT == "0/0" | snp_data_vcf$gt_GT == "0/1" |
snp_data_vcf$gt_GT == "1/1" ,snp_data_vcf$gt_GT,  '<NA>')
snp_data_vcf$gt_GT_alleles <- ifelse(snp_data_vcf$gt_GT == "0/0" | snp_data_vcf$gt_GT ==
"0/1" | snp_data_vcf$gt_GT == "1/1" ,snp_data_vcf$gt_GT_alleles,  '<NA>')
snp_data_vcf_genotype<-reshape2::dcast( snp_data_vcf , CHROM + POS ~ Indiv, value.var="g
t_GT")
snp_data_vcf_nucleotide<-reshape2::dcast( snp_data_vcf , CHROM + POS ~ Indiv, value.var=
"gt_GT_alleles")

snp_data_vcf_genotype_chr27<-snp_data_vcf_genotype[rowSums(snp_data_vcf_genotype== "<NA
>") < 26,]
snp_data_vcf_nucleotide_chr27<-snp_data_vcf_nucleotide[rowSums(snp_data_vcf_nucleotide==
```

```
"<NA>") < 26,]

rm(snp_data_vcf,snp_data_vcf_genotype,snp_data_vcf_nucleotide)

snp_data_vcf <- read.vcfR("/mnt/storage/lab_folder/heifer_infertility/alignment_SNP/2021
_11_3_samtools_variant_filtering_chr28.vcf.gz", verbose = FALSE)
snp_data_vcf <- extract.indels(snp_data_vcf, return.indels = FALSE )
snp_data_vcf <- vcfR2tidy(snp_data_vcf, info_only = FALSE, single_frame = TRUE, toss_INF
O_column = TRUE)
snp_data_vcf<-as.data.frame(snp_data_vcf$dat)
snp_data_vcf$gt_GT <- ifelse(snp_data_vcf$gt_DP > 10, snp_data_vcf$gt_GT,  '<NA>' )
snp_data_vcf$gt_GT_alleles <- ifelse(snp_data_vcf$gt_DP > 10, snp_data_vcf$gt_GT_allele
s,  '<NA>' )
snp_data_vcf$gt_GT <- ifelse(snp_data_vcf$gt_GT == "0/0" | snp_data_vcf$gt_GT == "0/1" |
snp_data_vcf$gt_GT == "1/1" ,snp_data_vcf$gt_GT,  '<NA>')
snp_data_vcf$gt_GT_alleles <- ifelse(snp_data_vcf$gt_GT == "0/0" | snp_data_vcf$gt_GT ==
"0/1" | snp_data_vcf$gt_GT == "1/1" ,snp_data_vcf$gt_GT_alleles,  '<NA>')
snp_data_vcf_genotype<-reshape2::dcast( snp_data_vcf , CHROM + POS ~ Indiv, value.var="g
t_GT")
snp_data_vcf_nucleotide<-reshape2::dcast( snp_data_vcf , CHROM + POS ~ Indiv, value.var=
"gt_GT_alleles")

snp_data_vcf_genotype_chr28<-snp_data_vcf_genotype[rowSums(snp_data_vcf_genotype== "<NA
>") < 26,]
snp_data_vcf_nucleotide_chr28<-snp_data_vcf_nucleotide[rowSums(snp_data_vcf_nucleotide==
"<NA>") < 26,]

rm(snp_data_vcf,snp_data_vcf_genotype,snp_data_vcf_nucleotide)

snp_data_vcf <- read.vcfR("/mnt/storage/lab_folder/heifer_infertility/alignment_SNP/2021
_11_3_samtools_variant_filtering_chrX.vcf.gz", verbose = FALSE)
snp_data_vcf <- extract.indels(snp_data_vcf, return.indels = FALSE )
snp_data_vcf <- vcfR2tidy(snp_data_vcf, info_only = FALSE, single_frame = TRUE, toss_INF
O_column = TRUE)
snp_data_vcf<-as.data.frame(snp_data_vcf$dat)
snp_data_vcf$gt_GT <- ifelse(snp_data_vcf$gt_DP > 10, snp_data_vcf$gt_GT,  '<NA>' )
snp_data_vcf$gt_GT_alleles <- ifelse(snp_data_vcf$gt_DP > 10, snp_data_vcf$gt_GT_allele
s,  '<NA>' )
snp_data_vcf$gt_GT <- ifelse(snp_data_vcf$gt_GT == "0/0" | snp_data_vcf$gt_GT == "0/1" |
snp_data_vcf$gt_GT == "1/1" ,snp_data_vcf$gt_GT,  '<NA>')
snp_data_vcf$gt_GT_alleles <- ifelse(snp_data_vcf$gt_GT == "0/0" | snp_data_vcf$gt_GT ==
"0/1" | snp_data_vcf$gt_GT == "1/1" ,snp_data_vcf$gt_GT_alleles,  '<NA>')
snp_data_vcf_genotype<-reshape2::dcast( snp_data_vcf , CHROM + POS ~ Indiv, value.var="g
t_GT")
snp_data_vcf_nucleotide<-reshape2::dcast( snp_data_vcf , CHROM + POS ~ Indiv, value.var=
"gt_GT_alleles")

snp_data_vcf_genotype_chrx<-snp_data_vcf_genotype[rowSums(snp_data_vcf_genotype== "<NA>"
) < 26,]
snp_data_vcf_nucleotide_chrx<-snp_data_vcf_nucleotide[rowSums(snp_data_vcf_nucleotide==
"<NA>") < 26,]
```

```
rm(snp_data_vcf,snp_data_vcf_genotype,snp_data_vcf_nucleotide)

gc()

merged_SNPS<-rbind(snp_data_vcf_genotype_chr1,snp_data_vcf_genotype_chr2, snp_data_vcf_genotype_chr3, snp_data_vcf_genotype_chr4, snp_data_vcf_genotype_chr5, snp_data_vcf_genotype_chr6, snp_data_vcf_genotype_chr7, snp_data_vcf_genotype_chr8, snp_data_vcf_genotype_chr9, snp_data_vcf_genotype_chr10, snp_data_vcf_genotype_chr11, snp_data_vcf_genotype_chr12, snp_data_vcf_genotype_chr13, snp_data_vcf_genotype_chr14, snp_data_vcf_genotype_chr15, snp_data_vcf_genotype_chr16, snp_data_vcf_genotype_chr17, snp_data_vcf_genotype_chr18, snp_data_vcf_genotype_chr19, snp_data_vcf_genotype_chr20, snp_data_vcf_genotype_chr21, snp_data_vcf_genotype_chr22, snp_data_vcf_genotype_chr23, snp_data_vcf_genotype_chr24, snp_data_vcf_genotype_chr25, snp_data_vcf_genotype_chr26, snp_data_vcf_genotype_chr27, snp_data_vcf_genotype_chr28, snp_data_vcf_genotype_chr29, snp_data_vcf_genotype_chrx)

rm(snp_data_vcf_genotype_chr1,snp_data_vcf_genotype_chr2, snp_data_vcf_genotype_chr3, snp_data_vcf_genotype_chr4, snp_data_vcf_genotype_chr5, snp_data_vcf_genotype_chr6, snp_data_vcf_genotype_chr7, snp_data_vcf_genotype_chr8, snp_data_vcf_genotype_chr9, snp_data_vcf_genotype_chr10,
   snp_data_vcf_genotype_chr11, snp_data_vcf_genotype_chr12, snp_data_vcf_genotype_chr13, snp_data_vcf_genotype_chr14, snp_data_vcf_genotype_chr15, snp_data_vcf_genotype_chr16, snp_data_vcf_genotype_chr17, snp_data_vcf_genotype_chr18, snp_data_vcf_genotype_chr19, snp_data_vcf_genotype_chr20,
   snp_data_vcf_genotype_chr21, snp_data_vcf_genotype_chr22, snp_data_vcf_genotype_chr23, snp_data_vcf_genotype_chr24, snp_data_vcf_genotype_chr25, snp_data_vcf_genotype_chr26, snp_data_vcf_genotype_chr27, snp_data_vcf_genotype_chr28, snp_data_vcf_genotype_chr29, snp_data_vcf_genotype_chrx)

merged_SNPS_nucleotide<- rbind(snp_data_vcf_nucleotide_chr1, snp_data_vcf_nucleotide_chr2, snp_data_vcf_nucleotide_chr3, snp_data_vcf_nucleotide_chr4, snp_data_vcf_nucleotide_chr5, snp_data_vcf_nucleotide_chr6, snp_data_vcf_nucleotide_chr7, snp_data_vcf_nucleotide_chr8, snp_data_vcf_nucleotide_chr9, snp_data_vcf_nucleotide_chr10, snp_data_vcf_nucleotide_chr11, snp_data_vcf_nucleotide_chr12, snp_data_vcf_nucleotide_chr13, snp_data_vcf_nucleotide_chr14, snp_data_vcf_nucleotide_chr15, snp_data_vcf_nucleotide_chr16, snp_data_vcf_nucleotide_chr17, snp_data_vcf_nucleotide_chr18, snp_data_vcf_nucleotide_chr19, snp_data_vcf_nucleotide_chr20, snp_data_vcf_nucleotide_chr21, snp_data_vcf_nucleotide_chr22, snp_data_vcf_nucleotide_chr23, snp_data_vcf_nucleotide_chr24, snp_data_vcf_nucleotide_chr25, snp_data_vcf_nucleotide_chr26, snp_data_vcf_nucleotide_chr27, snp_data_vcf_nucleotide_chr28, snp_data_vcf_nucleotide_chr29, snp_data_vcf_nucleotide_chrx )

rm(snp_data_vcf_nucleotide_chr1, snp_data_vcf_nucleotide_chr2, snp_data_vcf_nucleotide_chr3, snp_data_vcf_nucleotide_chr4, snp_data_vcf_nucleotide_chr5, snp_data_vcf_nucleotide_chr6, snp_data_vcf_nucleotide_chr7, snp_data_vcf_nucleotide_chr8, snp_data_vcf_nucleotide_chr9, snp_data_vcf_nucleotide_chr10,
   snp_data_vcf_nucleotide_chr11, snp_data_vcf_nucleotide_chr12, snp_data_vcf_nucleotide_chr13, snp_data_vcf_nucleotide_chr14, snp_data_vcf_nucleotide_chr15, snp_data_vcf_nucleotide_chr16, snp_data_vcf_nucleotide_chr17, snp_data_vcf_nucleotide_chr18, snp_data_vcf_nucleotide_chr19, snp_data_vcf_nucleotide_chr20,
   snp_data_vcf_nucleotide_chr21, snp_data_vcf_nucleotide_chr22, snp_data_vcf_nucleotide_chr23, snp_data_vcf_nucleotide_chr24, snp_data_vcf_nucleotide_chr25, snp_data_vcf_nucleotide_chr26, snp_data_vcf_nucleotide_chr27, snp_data_vcf_nucleotide_chr28, snp_data_vcf_nucleotide_chr29, snp_data_vcf_nucleotide_chrx )
```

```
gc()
```

```
saveRDS(merged_SNPS,'/mnt/storage/lab_folder/shared_R_codes/fernando/SNP_eqtl/merged_SNP
S_2021_11_13.rds', compress=FALSE)
saveRDS(merged_SNPS_nucleotide,'/mnt/storage/lab_folder/shared_R_codes/fernando/SNP_eqt
l/merged_SNPS_nucleotide_2021_11_13.rds', compress=FALSE)
```

```
merged_SNPS<-readRDS('/mnt/storage/lab_folder/shared_R_codes/fernando/SNP_eqtl/merged_SN
PS_2021_11_13.rds')
merged_SNPS_nucleotide<-readRDS('/mnt/storage/lab_folder/shared_R_codes/fernando/SNP_eqt
l/merged_SNPS_nucleotide_2021_11_13.rds')
```

#read in expression data

```
sourcefile_path<-"/mnt/storage/auburn/heifer_pregnancy/proj_2018/analysis/resources"

load( file=paste(sourcefile_path,"resource_data_2019_06_28.RData", sep="/"), verbose =TR
UE)
```

```
## Loading objects:
##   count_miRNA_plasma
##   count_pwbc
##   count_pwbc_2017
##   gene.length
##   annotation.ensembl.symbol
##   annotation.GO.biomart
##   bta_miRWalk_CDS
##   bta_miRWalk_3UTR
##   bta_miRWalk_5UTR
```

```
rm(count_miRNA_plasma,annotation.GO.biomart,bta_miRWalk_CDS,bta_miRWalk_3UTR,bta_miRWalk
_5UTR)

count_pwbc<-count_pwbc[,-3]
#Filter to remove lowly expressed genes
keep<-rowSums( cpm(count_pwbc) >= 2 ) >= 5
count_pwbc<-count_pwbc[keep,]

count_pwbc<-merge(count_pwbc,annotation.ensembl.symbol, by.x="row.names", by.y="ensembl_
gene_id", all=FALSE)
count_pwbc<-count_pwbc[ which(count_pwbc$gene_biotype=='protein_coding'), ]
rownames(count_pwbc)<-count_pwbc$Row.names
count_pwbc<-count_pwbc[,2:18]
#count_pwbc<-count_pwbc[,c(10,4,16,18,8,5,9,15,11,2,6,12,3,13,14,7,17)]

#count_pwbc_2017<-count_pwbc_2017[,c(13,14,15,16,17,18,19,20,21,22,23,24)]
keep<-rowSums( cpm(count_pwbc_2017) >= 2 ) >= 6
count_pwbc_2017<-count_pwbc_2017[keep,]
dim(count_pwbc_2017) #12432
```

```
## [1] 12432    24
```

```
#Filter 2017 data to retain only protein coding genes
count_pwbc_2017<-merge(count_pwbc_2017,annotation.ensembl.symbol, by.x="row.names", by.y
="ensembl_gene_id", all=FALSE)
count_pwbc_2017<-count_pwbc_2017[ which(count_pwbc_2017$gene_biotype=='protein_coding'),
]
rownames(count_pwbc_2017)<-count_pwbc_2017[,1]
count_pwbc_2017<-count_pwbc_2017[,c(2:25)]

merged_datasets<-merge(count_pwbc,count_pwbc_2017, by="row.names", all=FALSE)
rownames(merged_datasets)<-merged_datasets[,1]
merged_datasets<-merged_datasets[,2:42]

gene_length_merged<-subset(gene.length, Geneid %in% rownames(merged_datasets))
gene_length_merged<-gene_length_merged[order(gene_length_merged$Geneid),]
merged_datasets<-merged_datasets[order(rownames(merged_datasets)),]
table(rownames(merged_datasets)==gene_length_merged$Geneid) #all True
```

```
##
##  TRUE
## 10332
```

```
cpm_merged_datasets<-cpm(merged_datasets,normalized.lib.sizes = TRUE,log = FALSE)

x<- merged_datasets/gene_length_merged$Length
tpm_merged_datasets<-t( t(x) * 1e6 / colSums(x) )
rm(x)
```

#convert to TMM counts per million and normal transform

```
lib_size <- base::colSums(merged_datasets)
norm_factors <- calcNormFactors(object = merged_datasets, lib.size = lib_size, method =
"TMM")
normalized_lib_size<-colSums(merged_datasets) * norm_factors
tmm_per_million_tmm_per_million_normalized_expression<-edgeR::cpm(merged_datasets, lib.s
ize = normalized_lib_size)

n <- ncol(tmm_per_million_tmm_per_million_normalized_expression)
zvalues <- qnorm(ppoints(n))
TMM_tmm_per_million_normalized_expression_a <- tmm_per_million_tmm_per_million_normalize
d_expression
for (i in 1:nrow(TMM_tmm_per_million_normalized_expression_a)) {TMM_tmm_per_million_norm
alized_expression_a[i,] <- RankNorm(as.numeric(TMM_tmm_per_million_normalized_expression
_a[i,]), k = 0.375)}

TMM_tmm_per_million_normalized_expression_a <- as.matrix(TMM_tmm_per_million_normalized_
expression_a)
```

#Supplementary figure 1

```
TMM_tmm_per_million_normalized_expression_for_plotting<-reshape2::melt(as.matrix(tmm_per
_million_tmm_per_million_normalized_expression[1:5,]))

Supp_fig_01_A<-ggplot(data=TMM_tmm_per_million_normalized_expression_for_plotting, aes(x
=value))+
geom_histogram()+
facet_wrap(~Var1, scale="free", ncol=1)+
scale_x_continuous(name="TMM")+
theme_classic()

TMM_normalized_transformed_expression_for_plotting<-reshape2::melt(as.matrix(TMM_tmm_per
_million_normalized_expression_a[1:5,]))


Supp_fig_01_B<-ggplot(data=TMM_normalized_transformed_expression_for_plotting, aes(x=val
ue))+
geom_histogram()+
facet_wrap(~Var1, scale="free", ncol=1)+
scale_x_continuous(name="TMM")+
theme_classic()

Supp_fig_01_C<-ggplot(data=TMM_normalized_transformed_expression_for_plotting, aes(sampl
e=value))+
stat_qq() + stat_qq_line()+
facet_wrap(~Var1, scale="free", ncol=1)+
theme_classic()

plot_grid(Supp_fig_01_A, Supp_fig_01_B, Supp_fig_01_C, ncol=3,labels=c("A", "B", "C"),la
bel_fontface = "plain",label_size = 12)
```

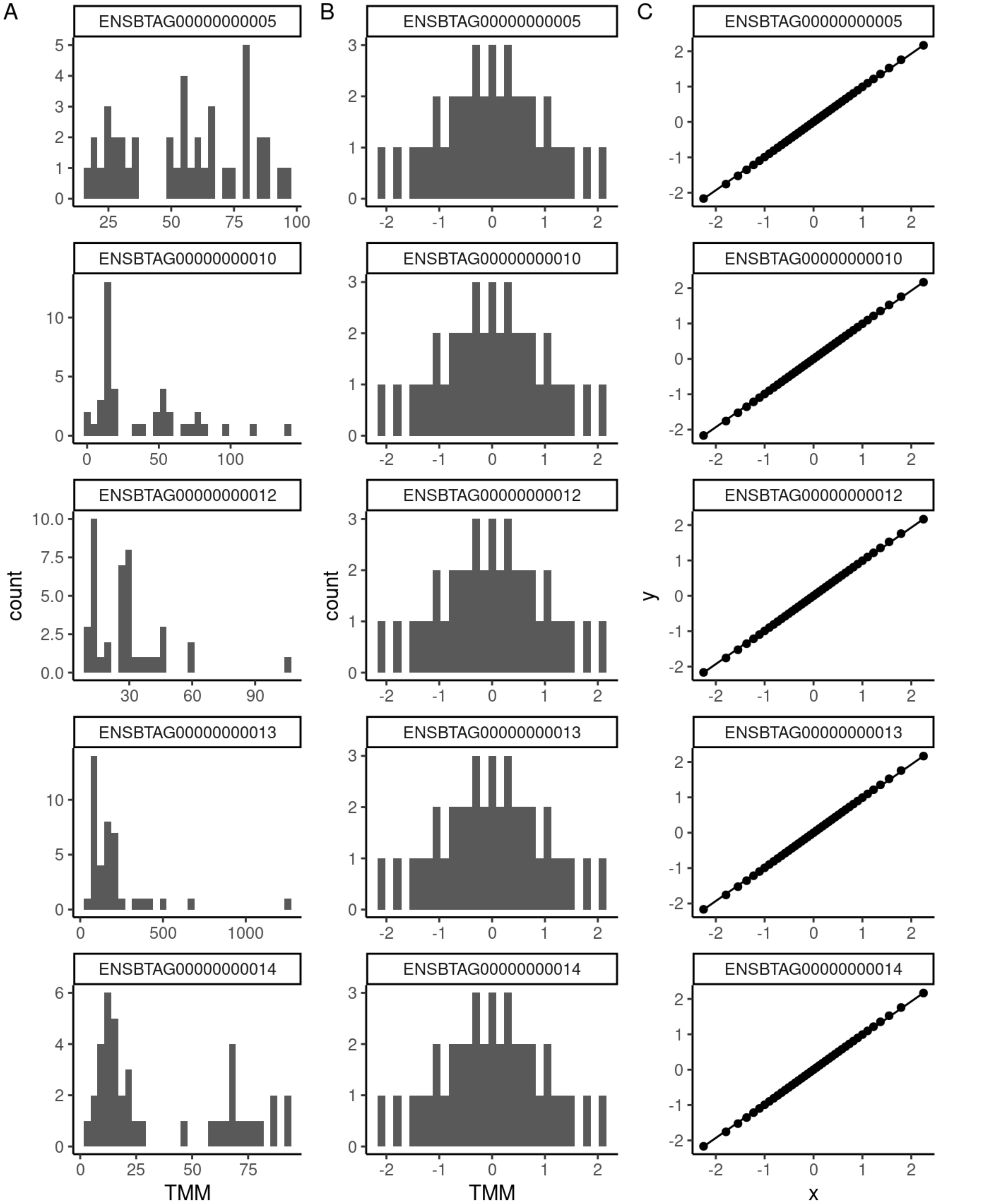


#set up genotype files for eQTL analysis

```
genotypes<-merged_SNPS[rowSums(is.na(merged_SNPS) )<20,]
rownames(genotypes) <- paste(genotypes$CHROM, genotypes$POS, sep=":")

genotypes[genotypes=="0/0"]<-as.numeric(0)
genotypes[genotypes=="0|0"]<-as.numeric(0)

genotypes[genotypes=="0/1"]<-as.numeric(1)
genotypes[genotypes=="0|1"]<-as.numeric(1)

genotypes[genotypes=="1/0"]<-as.numeric(1)
genotypes[genotypes=="1|0"]<-as.numeric(1)

genotypes[genotypes=="1/1"]<-as.numeric(2)
genotypes[genotypes=="1|1"]<-as.numeric(2)

genotypes_a<- genotypes[,3:44]
```

#Hardy Weinberg equlibrium

```
genot_counting<-data.frame(cbind(
                                 row_count(genotypes_a, count = 0, append = FALSE),
                                 row_count(genotypes_a, count = 1, append = FALSE),
                                 row_count(genotypes_a, count = 2, append = FALSE)))

genot_counting_filtered <- genot_counting[!(genot_counting$rowcount==0 & genot_counting
$rowcount.1==0 & genot_counting$rowcount.2>0),]
genot_counting_filtered <- genot_counting_filtered[!(genot_counting_filtered$rowcount>0
 & genot_counting_filtered$rowcount.1==0 & genot_counting_filtered$rowcount.2==0),]



genot_counting_filtered$alele_ref<-( (2 * genot_counting_filtered$rowcount) +  genot_cou
nting_filtered$rowcount.1) / (2* (genot_counting_filtered$rowcount + genot_counting_filt
ered$rowcount.1 + genot_counting_filtered$rowcount.2))

colnames(genot_counting_filtered)<-c("AA","AB","BB","alele_ref")

genot_counting_filtered$pvalue <- NA

for(index in 1:nrow(genot_counting_filtered)){
   genot_counting_filtered[index,5]<- HWExact(unlist(genot_counting_filtered[index,1:3
]), verbose = FALSE)$pval

}

genot_counting_filtered<-genot_counting_filtered[order( genot_counting_filtered$pvalu
e),]
genot_counting_filtered<-genot_counting_filtered[order( genot_counting_filtered$alele_re
f),]
genot_counting_filtered$HWB_sig_fdr<-p.adjust(genot_counting_filtered$pvalue, method="fd
r")
genot_counting_filtered$HWB_sig<-ifelse(genot_counting_filtered$HWB_sig_fdr < 0.01, "si
g", "not_sig")
genot_counting_filtered$HWB_sig<-factor(genot_counting_filtered$HWB_sig, levels= c( "not
_sig", "sig"))

genot_counting_filtered_a<-genot_counting_filtered[genot_counting_filtered$AA >=5 & geno
t_counting_filtered$AB >=5 & genot_counting_filtered$BB >=5,]

genot_counting_filtered_b<-genot_counting_filtered_a[genot_counting_filtered_a$HWB_sig==
"not_sig",]
```

#Supplementary figure 2

```
#set up genotype files for plink and pca analysis

#Ped file
rownames(merged_SNPS_nucleotide) <- paste(merged_SNPS_nucleotide$CHROM, merged_SNPS_nucl
eotide$POS, sep=":")
genotypes<-merged_SNPS_nucleotide[rownames(merged_SNPS_nucleotide) %in% rownames(genot_c
ounting_filtered_b),]
genotypes<-subset(genotypes, select=-c(SL297953))

genotypes_plink<-genotypes

genotypes_plink[genotypes_plink=="<NA>"]<-"0 0"
genotypes_plink<-t(genotypes_plink[ , 3:dim(genotypes_plink)[2]])
genotypes_plink <- as.data.frame(apply(genotypes_plink,2,function(x) gsub("/", " ", x)))
genotypes_plink <- as.data.frame(apply(genotypes_plink,2,function(x) gsub("|", " ", x)))

genotypes_plink<-cbind( data.frame( rownames(genotypes_plink), rownames(genotypes_plin
k), rep(0,41), rep(0,41), rep(2,41), rep(0,41)),genotypes_plink)

#Map file

map_file_plink<- data.frame(paste( "Chr", genotypes$CHROM, sep=""), paste(genotypes$CHRO
M,genotypes$POS, sep=":") , rep(0,dim(genotypes)[1]), genotypes$POS)

#write.table(genotypes_plink, "/mnt/storage/lab_folder/shared_R_codes/fernando/SNP_eqtl/
plink_files/genotypes_plink.ped", col.names = FALSE, row.names = FALSE, sep = " " , quot
e = FALSE)
#write.table(map_file_plink, "/mnt/storage/lab_folder/shared_R_codes/fernando/SNP_eqtl/p
link_files/genotypes_plink.map", col.names = FALSE, row.names = FALSE, sep = " " , quote
= FALSE)

#system("/home/fbiase/bioinfo/plink --file /mnt/storage/lab_folder/shared_R_codes/fernan
do/SNP_eqtl/plink_files/genotypes_plink --cow --make-bed --out /mnt/storage/lab_folder/s
hared_R_codes/fernando/SNP_eqtl/plink_files/genotypes_plink")

#system("/home/fbiase/bioinfo/plink --bfile /mnt/storage/lab_folder/shared_R_codes/ferna
ndo/SNP_eqtl/plink_files/genotypes_plink --cow   --pca --out /mnt/storage/lab_folder/sha
red_R_codes/fernando/SNP_eqtl/plink_files/genotypes_plink")
```

```
eval <- data.table::fread("/mnt/storage/lab_folder/shared_R_codes/fernando/SNP_eqtl/plin
k_files/genotypes_plink.eigenval", data.table = FALSE)
evec <- data.table::fread("/mnt/storage/lab_folder/shared_R_codes/fernando/SNP_eqtl/plin
k_files/genotypes_plink.eigenvec", data.table = FALSE)

percentage_PCA1<-round((eval$V1[1] / sum(eval$V1) )*100 ,2)
percentage_PCA2<-round((eval$V1[2] / sum(eval$V1) )*100 ,2)

  ggplot(evec) +
  geom_point(aes(V3, V4, shape=factor(rep(c("location 1","location 2","location 2"),c(12
,12,17)))), size=3  )+
  labs(x = paste("PC1: ",percentage_PCA1,"% variance", sep=""), y = paste("PC2: ",percen
tage_PCA2,"% variance", sep=""))+
  ggtitle("PCA SNPs")+
  theme_minimal(base_size = 15)+
 theme(
    axis.text = element_blank(),
    plot.margin=grid::unit(c(0,0,0,0), "mm"),
    legend.position = "bottom",
    legend.title = element_blank(),
    legend.text  = element_text(size=10),
    plot.title = element_text(hjust = 0.5, size=15)
  )
```

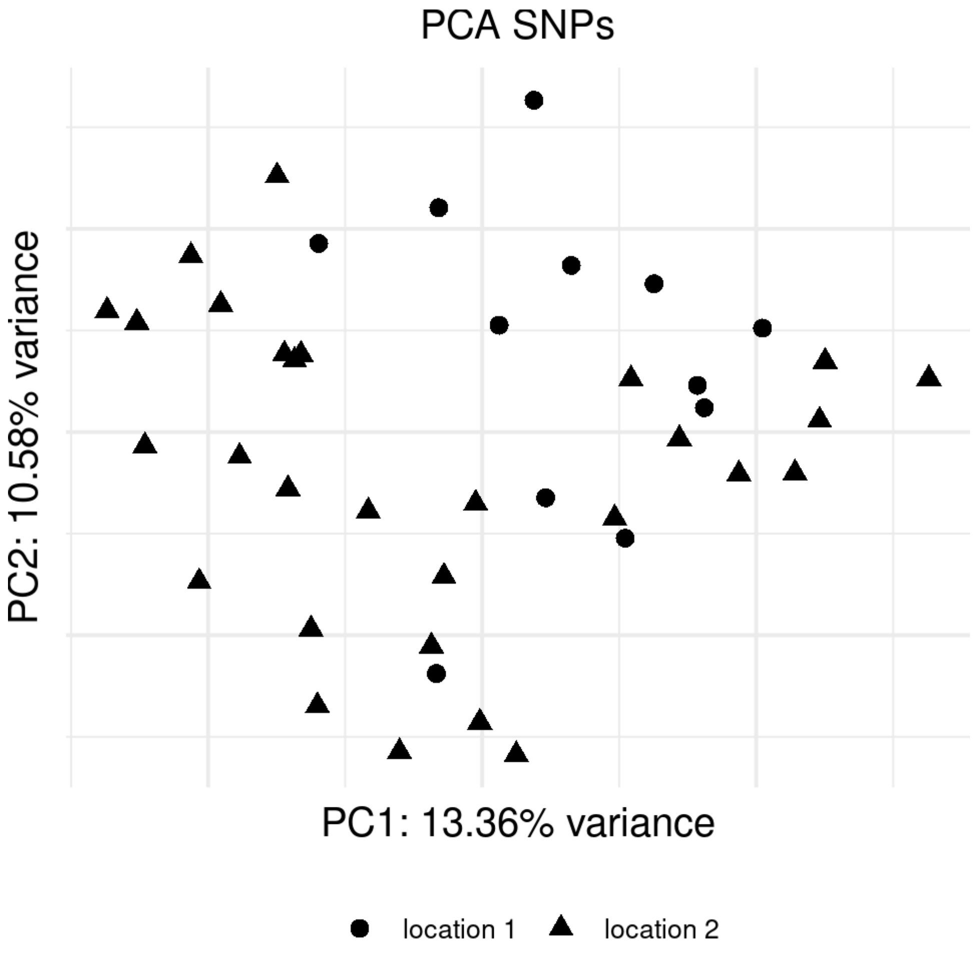


#Figure 1B and C

```
new_labels <- c("HW equilibrium", "not in HW equilibrium")
names(new_labels) <- c("not_sig", "sig")

figure_1_B<-ggplot(data=genot_counting_filtered, aes(y=alele_ref, 1:nrow(genot_counting_
filtered), color=HWB_sig))+
  geom_point()+
  facet_wrap(.~HWB_sig, labeller = labeller(HWB_sig = new_labels))+
  scale_y_continuous(name="Frequency reference allele")+
  scale_x_continuous(name="Single nucleotide polymorphisms")+
  theme_classic()+
  theme(
    legend.position = "none",
    axis.title = element_text(size=12, color="black"),
    axis.text.y = element_text(size=12, color="black"),
    axis.text.x = element_blank(),
    strip.text.x = element_text(size=10, color="black"),
  )

new_labels <- c("HW equilibrium, MAF>0.15 and >4 individuals in each genotype", "not in
 HW equilibrium")
names(new_labels) <- c("not_sig", "sig")

figure_1_C<-ggplot(data=genot_counting_filtered_b, aes(y=alele_ref, 1:nrow(genot_countin
g_filtered_b), color=HWB_sig))+
  geom_point()+
  facet_wrap(.~HWB_sig, labeller = labeller(HWB_sig = new_labels))+
  scale_y_continuous(name="Frequency reference allele")+
  scale_x_continuous(name="Single nucleotide polymorphisms")+
  theme_classic()+
  theme(
    legend.position = "none",
    axis.title = element_text(size=12, color="black"),
    axis.text.y = element_text(size=12, color="black"),
    axis.text.x = element_blank(),
    strip.text.x = element_text(size=10, color="black"),
  )

plot_grid(figure_1_B,NULL, figure_1_C, nrow=3, rel_heights=c(0.8,0.1,0.8))
```

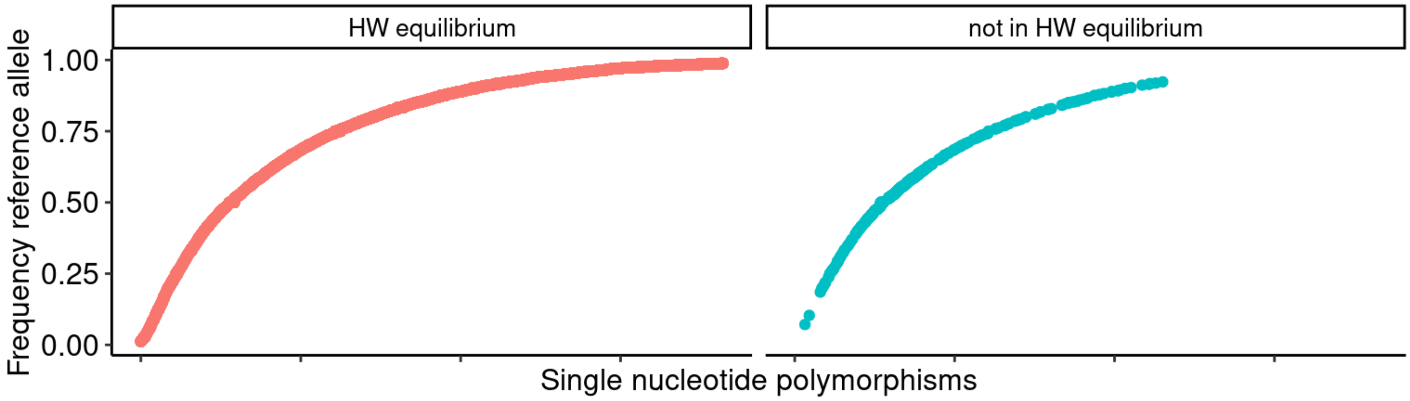


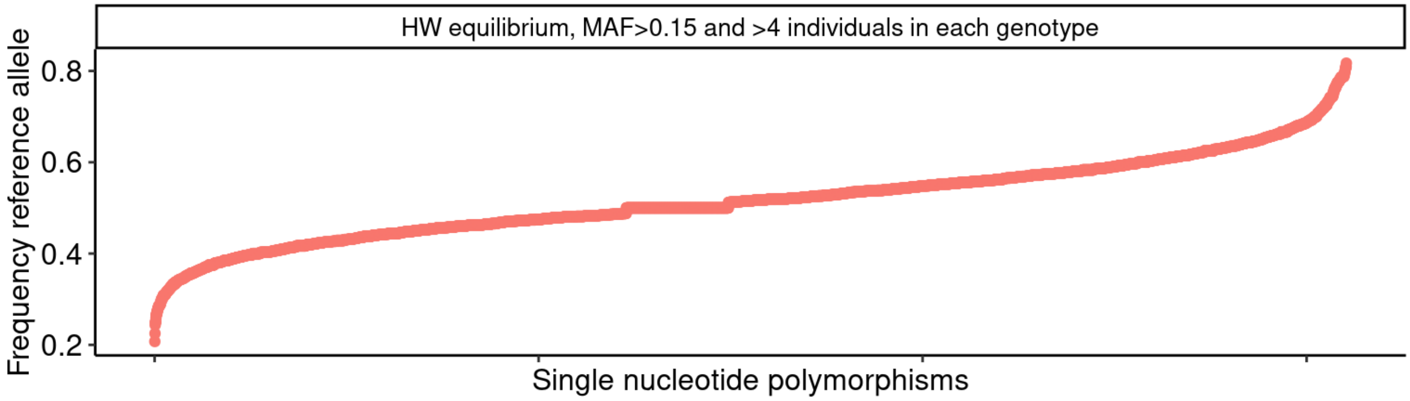


#Supplementary table 1

```
SNP_annotation_all<- read.table("/mnt/storage/lab_folder/shared_R_codes/fernando/SNP_eqt
l/xNEZMz1bkTufDelg.txt", header = TRUE, sep= "\t",comment.char="", stringsAsFactors = FA
LSE)

SNP_annotation_all<-SNP_annotation_all[,!(colnames(SNP_annotation_all) %in% c("Feature",
"EXON", "INTRON", "HGVSc", "HGVSp",  "FLAGS" , "cDNA_position" ,"CDS_position" , "Codon
s", "Protein_position", "Amino_acids", "SYMBOL_SOURCE" ,"HGNC_ID", "SIFT" ,"CLIN_SIG",
"SOMATIC", "PHENO","APPRIS", "MANE_SELECT", "MANE_PLUS_CLINICAL", "TSL"))]
SNP_annotation_all<-SNP_annotation_all[order(SNP_annotation_all$X.Uploaded_variation ,SN
P_annotation_all$Consequence,SNP_annotation_all$Gene),]
SNP_annotation_all<-SNP_annotation_all[!duplicated(SNP_annotation_all),]

SNP_annotation_all$SNPs<-ifelse(SNP_annotation_all$Existing_variation=="-","putative_ne
w", "SNPdb")
SNP_annotation_all_a <- SNP_annotation_all %>% separate(Location,c("SNP","Location2"), s
ep = "-")
#n snps in database
#length(unique(SNP_annotation_all[SNP_annotation_all$SNPs=="SNPdb",]$X.Uploaded_variatio
n))
#n snps not in database
#length(unique(SNP_annotation_all[SNP_annotation_all$SNPs=="putative_new",]$X.Uploaded_v
ariation))



#all
require(dplyr)

df1<-SNP_annotation_all %>% group_by(SNPs,Consequence) %>% summarize(n=n()) %>% arrange
(desc(SNPs),desc(n))
```

```
## `summarise()` has grouped output by 'SNPs'. You can override using the
## `.groups` argument.
```

```
n_total_size_SNPdb<-sum(df1$n[1:24])
n_total_size_new <- sum(df1$n[25:36])

df1$percentage <- c(round(df1$n[1:24]/n_total_size_SNPdb * 100, 2), round(df1$n[25:36]/n
_total_size_new * 100, 2))
```

#set up genotype files for eQTL analysis

```
genotypes_a<-genotypes_a[rowSums(genotypes_a=='1',na.rm=TRUE)>=5,]
genotypes_a<-genotypes_a[rowSums(genotypes_a=='2',na.rm=TRUE)>=5,]
genotypes_a<-genotypes_a[rowSums(genotypes_a=='0',na.rm=TRUE)>=5,]
genotypes_a1<-data.frame(lapply(genotypes_a,as.numeric))

genotypes_a2<-as.matrix(genotypes_a1,rownames = TRUE)
genotypes_c<-genotypes_a2[,-27]
rownames(genotypes_c)<- rownames(genotypes_a)

rm(genotypes,genotypes_a1,genotypes_a2)

genotypes_d<- genotypes_c[rownames(genotypes_c) %in% rownames(genot_counting_filtered_
b),]
genotypes_d <- genotypes_d[,intersect( colnames(TMM_tmm_per_million_normalized_expressio
n_a), colnames(genotypes_d))]

expression<-(TMM_tmm_per_million_normalized_expression_a[,intersect( colnames(TMM_tmm_pe
r_million_normalized_expression_a), colnames(genotypes_d))])
```

```
set.seed(321)

covariates = character()
snps = SlicedData$new()
snps$CreateFromMatrix(genotypes_d)

gene = SlicedData$new()
gene$CreateFromMatrix(expression)

cvrt = SlicedData$new()

eQTL_analysis_anova_all = "/mnt/storage/lab_folder/shared_R_codes/fernando/SNP_eqtl/resu
lts/eQTL_analysis_anova_filtered_2022_11_07.txt"

system.time(
Matrix_eQTL_main(
  snps = snps,
  gene = gene,
  cvrt = cvrt,
  useModel =modelANOVA,
  output_file_name = eQTL_analysis_anova_all,
  pvOutputThreshold = 1,
  verbose = TRUE,
  pvalue.hist = FALSE,
  min.pv.by.genesnp = FALSE,
  noFDRsaveMemory = TRUE)
)

#output system.time() when subsetting only for the results that printed at 5e-08
#    user   system  elapsed
#   3.761   1.085   4.699

#system("lbzip2  --compress -9 --quiet /mnt/storage/lab_folder/shared_R_codes/fernando/S
NP_eqtl/results/eQTL_analysis_anova_filtered_2022_11_07.txt")
```

#TMM anova annotation

```
genotypes_d1<- cbind(as.data.frame(genotypes_d), row_count(as.data.frame(genotypes_d), c
ount = 1, append = FALSE))
genotypes_d1<- cbind(genotypes_d1, row_count(genotypes_d1, count = 2, append = FALSE))
genotypes_d1<- cbind(genotypes_d1, row_count(genotypes_d1, count = 0, append = FALSE))

colnames(genotypes_d1)[42]<-"rowcounts_1"
colnames(genotypes_d1)[43]<-"rowcounts_2"
colnames(genotypes_d1)[44]<-"rowcounts_0"

eqtl_annotate_anova_TMM<- as.data.frame(fread("/mnt/storage/lab_folder/shared_R_codes/fe
rnando/SNP_eqtl/results/eQTL_analysis_anova_filtered_2022_11_07.txt.bz2", header = TRUE,
showProgress=FALSE))

eqtl_annotate_anova_TMM$q.values<-qvalue(eqtl_annotate_anova_TMM[,4])$qvalues
eqtl_annotate_anova_TMM$bonferroni<-p.adjust(eqtl_annotate_anova_TMM[,4],method="bonferr
oni")
eqtl_annotate_anova_TMM$FDR<-p.adjust(eqtl_annotate_anova_TMM[,4],method="fdr")
eqtl_annotate_anova_TMM1<-eqtl_annotate_anova_TMM[eqtl_annotate_anova_TMM[,4] < 5e-08,]
dim(eqtl_annotate_anova_TMM1)
```

```
## [1] 35  7
```

```
eqtl_annotate_anova_TMM1<-merge(eqtl_annotate_anova_TMM1, merged_SNPS_nucleotide, by.x=
c("SNP"), by.y= "row.names", all=FALSE)
eqtl_annotate_anova_TMM1<- merge(eqtl_annotate_anova_TMM1, genotypes_d1,by.x= c("SNP"),
 by.y= 'row.names', all=FALSE)
eqtl_annotate_anova_TMM1<-merge(eqtl_annotate_anova_TMM1, annotation.ensembl.symbol, by.
x="gene", by.y="ensembl_gene_id", all=FALSE)
eqtl_annotate_anova_TMM1$external_gene_name<-ifelse(eqtl_annotate_anova_TMM1$external_ge
ne_name=="", eqtl_annotate_anova_TMM1$gene, eqtl_annotate_anova_TMM1$external_gene_name)
eqtl_annotate_anova_TMM1<-eqtl_annotate_anova_TMM1[order(eqtl_annotate_anova_TMM1[,1]),]
```

#Figure 2

```
snps_sub2 = eqtl_annotate_anova_TMM1$SNP
genes_sub2 = eqtl_annotate_anova_TMM1$gene
gene_symbol = eqtl_annotate_anova_TMM1$external_gene_name

plot_list<-list()

for (index in seq(length(snps_sub2))){
  genotype_sub2 = unlist(merged_SNPS_nucleotide[snps_sub2[index],c(3:44)])
  expression_sub2 = tpm_merged_datasets[genes_sub2[index],]
  #lm_result = lm(expression_sub2 ~ genotype_sub2)
  genotype_sub2<-genotype_sub2[names(expression_sub2)]

  graph_data_frame<-data.frame(genotype_sub2=gsub("/","",genotype_sub2),expression_sub2,
snp=snps_sub2[index], gene_symbol=gene_symbol[index])
  graph_data_frame<-graph_data_frame[complete.cases(graph_data_frame),]
  graph_data_frame<-graph_data_frame[!(graph_data_frame$genotype_sub2 =="<NA>"),]

  plot_list[[index]]<-ggplot(data=graph_data_frame, aes(x=as.factor(genotype_sub2), y=ex
pression_sub2))+
    geom_boxplot(fill='transparent',outlier.shape = 4, outlier.color = "blue", size=0.1)
+
    geom_jitter(width=0.2, size=0.6)+
    scale_x_discrete(name= graph_data_frame$snp[1])+
    scale_y_continuous(name =  graph_data_frame$gene_symbol[1])+
    theme_classic(base_size = 7)+
    theme( axis.text=element_text(size=7,color="black"),
           axis.title=element_text(size=7,color="black"))
}
pdf(file="/mnt/storage/lab_folder/shared_R_codes/fernando/SNP_eqtl/results/Figure2.pdf",
width=7, height=6)
plot_grid( plotlist = plot_list, nrow = 5)
dev.off()
```

```
## png
##   2
```

```
eqtl_annotate_anova_TMM1<-merge(eqtl_annotate_anova_TMM1, SNP_annotation_all_a, by.x="SN
P", by.y="SNP", all=FALSE)
#eqtl_annotate_anova_TMM1<-eqtl_annotate_anova_TMM1[!duplicated(eqtl_annotate_anova_TMM1
$SNP,eqtl_annotate_anova_TMM1$gene),]
eqtl_annotate_anova_TMM1<-eqtl_annotate_anova_TMM1[order(eqtl_annotate_anova_TMM1[,4]),]

#write.table(eqtl_annotate_anova_TMM1,"/mnt/storage/lab_folder/shared_R_codes/fernando/S
NP_eqtl/results/Supplementary_table_2.txt", col.names = TRUE, row.names = FALSE, quote =
FALSE,sep ="\t")

dim(eqtl_annotate_anova_TMM1)
```

```
## [1]  40 113
```

#TMM linear eQTL analysis

```
set.seed(321)
eQTL_analysis_linear_all = "/mnt/storage/lab_folder/shared_R_codes/fernando/SNP_eqtl/res
ults/eQTL_analysis_linear_filtered_2022_11_07.txt"

system.time(
Matrix_eQTL_main(
  snps = snps,
  gene = gene,
  cvrt = cvrt,
  useModel =modelLINEAR,
  output_file_name = eQTL_analysis_linear_all,
  pvOutputThreshold = 1,
  verbose = TRUE,
  pvalue.hist = FALSE,
  min.pv.by.genesnp = FALSE,
  noFDRsaveMemory = TRUE) )

#output system.time() when subsetting only for the results that printed at 5e-08
# user  system elapsed
#  1.807   0.824   2.473

#  user   system  elapsed
# 263.628 2384.044 2123.196

system("lbzip2  --compress -9 --quiet /mnt/storage/lab_folder/shared_R_codes/fernando/SN
P_eqtl/results/eQTL_analysis_linear_filtered_2022_11_07.txt")
```

#linear annotation

```
eqtl_annotate_linear_TMM<- as.data.frame(fread("/mnt/storage/lab_folder/shared_R_codes/f
ernando/SNP_eqtl/results/eQTL_analysis_linear_filtered_2022_11_07.txt.bz2", header = TRU
E, showProgress=FALSE))

#eqtl_annotate_linear_TMM$q.values<-qvalue(eqtl_annotate_linear_TMM[,5])$qvalues
#eqtl_annotate_linear_TMM$bonferroni<-p.adjust(eqtl_annotate_linear_TMM[,5],method="bonf
erroni")
#eqtl_annotate_linear_TMM$FDR<-p.adjust(eqtl_annotate_linear_TMM[,5],method="fdr")
eqtl_annotate_linear_TMM1<-eqtl_annotate_linear_TMM[eqtl_annotate_linear_TMM[,5] < 5e-08
,]
dim(eqtl_annotate_linear_TMM1)
```

```
## [1] 39  5
```

```
eqtl_annotate_linear_TMM1<-merge(eqtl_annotate_linear_TMM1, merged_SNPS_nucleotide, by.x
= c("SNP"), by.y= "row.names", all=FALSE)
eqtl_annotate_linear_TMM1<- merge(eqtl_annotate_linear_TMM1, genotypes_d1,by.x= c("SNP"
), by.y= 'row.names', all=FALSE)
eqtl_annotate_linear_TMM1<-merge(eqtl_annotate_linear_TMM1, annotation.ensembl.symbol, b
y.x="gene", by.y="ensembl_gene_id", all=FALSE)
eqtl_annotate_linear_TMM1$external_gene_name<-ifelse(eqtl_annotate_linear_TMM1$external_
gene_name=="", eqtl_annotate_linear_TMM1$gene, eqtl_annotate_linear_TMM1$external_gene_n
ame)
eqtl_annotate_linear_TMM1<-eqtl_annotate_linear_TMM1[order(eqtl_annotate_linear_TMM1[,5
]),]
```

#Figure 3

```
snps_sub2 = eqtl_annotate_linear_TMM1$SNP
genes_sub2 = eqtl_annotate_linear_TMM1$gene
gene_symbol = eqtl_annotate_linear_TMM1$external_gene_name

plot_list<-list()

for (index in seq(length(snps_sub2))){
  genotype_sub2 = unlist(merged_SNPS_nucleotide[snps_sub2[index],c(3:44)])
  expression_sub2 = tpm_merged_datasets[genes_sub2[index],]
  #lm_result = lm(expression_sub2 ~ genotype_sub2)
  genotype_sub2<-genotype_sub2[names(expression_sub2)]

  graph_data_frame<-data.frame(genotype_sub2=gsub("/","",genotype_sub2),expression_sub2,
snp=snps_sub2[index], gene_symbol=gene_symbol[index])
  graph_data_frame<-graph_data_frame[complete.cases(graph_data_frame),]
  graph_data_frame<-graph_data_frame[!(graph_data_frame$genotype_sub2 =="<NA>"),]

  plot_list[[index]]<-ggplot(data=graph_data_frame, aes(x=as.factor(genotype_sub2), y=ex
pression_sub2))+
    geom_boxplot(fill='transparent',outlier.shape = 4, outlier.color = "blue", size=0.1)
+
    geom_jitter(width=0.2, size=0.6)+
    scale_x_discrete(name= NULL)+
    scale_x_discrete(name= graph_data_frame$snp[1])+
    scale_y_continuous(name =  graph_data_frame$gene_symbol[1])+
    theme_classic(base_size = 7)+
    theme( axis.text=element_text(size=7,color="black"),
           axis.title=element_text(size=7,color="black"))
}
#pdf(file="/mnt/storage/lab_folder/shared_R_codes/fernando/SNP_eqtl/results/Figure3.pd
f",width=7, height=6)
plot_grid( plotlist = plot_list, nrow = 5)
```

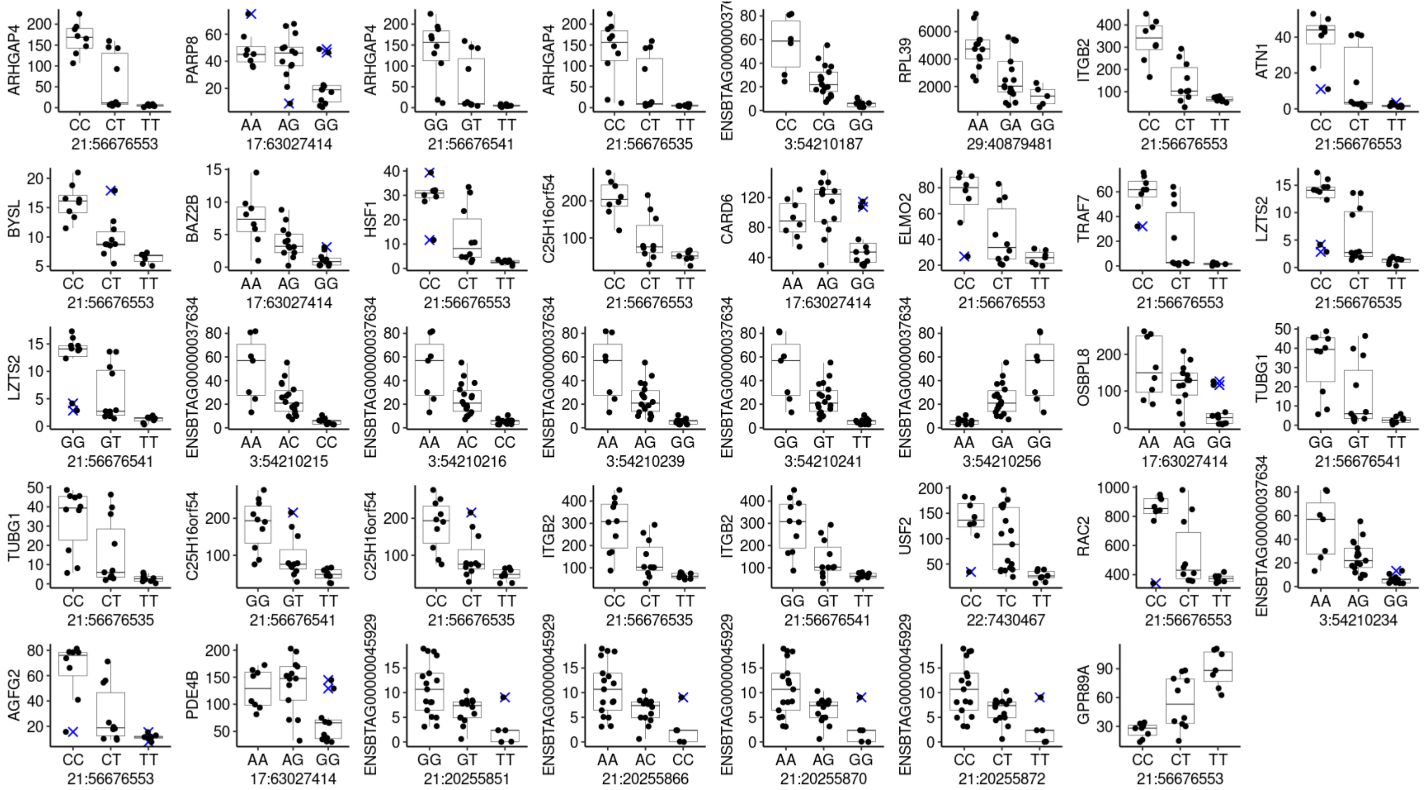


```
#dev.off()
```

```
eqtl_annotate_linear_TMM1<-merge(eqtl_annotate_linear_TMM1, SNP_annotation_all_a, by.x=
"SNP", by.y="SNP", all=FALSE)
eqtl_annotate_linear_TMM1<-eqtl_annotate_linear_TMM1[order(eqtl_annotate_linear_TMM1[,5
]),]

#write.table(eqtl_annotate_linear_TMM1, file= "/mnt/storage/lab_folder/shared_R_codes/fe
rnando/SNP_eqtl/results/Supplementary_table_3.txt", append = FALSE, quote = FALSE, sep =
"\t" ,row.names = FALSE)
eqtl_annotate_linear_TMM1<-eqtl_annotate_linear_TMM1[order(eqtl_annotate_linear_TMM1[,1
]),]
```

### EdgeR eQTL analysis

#### ANOVA contrasts

```
set.seed(321)
remove_outliers <- function(x, na.rm = TRUE, ...) {
  qnt <- quantile(x, probs=c(.25, .75), na.rm = na.rm, ...)
  H <- 3.5 * IQR(x, na.rm = na.rm)  #10 was good gave 28 eQTL
  y <- x
  #y[x < (qnt[1] - H)] <- NA
  y[x > (qnt[2] + H)] <- NA
  y
}

merged_datasets_outlier_removal_a<-data.frame()
merged_datasets_outlier_removal<- merged_datasets
for (i in 1:nrow(merged_datasets_outlier_removal)) {
  merged_datasets_outlier_removal_a <- rbind(merged_datasets_outlier_removal_a,remove_ou
tliers(merged_datasets_outlier_removal[i,]))
}

#dim(merged_datasets_outlier_removal_a[complete.cases(merged_datasets_outlier_removal_
a),])

merged_datasets_outlier_removal_a<-merged_datasets_outlier_removal_a[complete.cases(merg
ed_datasets_outlier_removal_a),]
```

```
rm(results_anova)
rm(results_zero_vs_one_two)
rm(results_zero_one_vs_two)
rm(results_one_vs_two)
rm(results_zero_vs_one)
rm(results_zero_vs_two)
```

```
cl <- makeCluster(34)
registerDoParallel(cl)

system.time (

results_anova<- foreach(i = seq(1:dim(genotypes_d)[1]) ,.combine = 'rbind', .inorder=FAL
SE , .errorhandling="remove", .packages="edgeR",.verbose=FALSE ) %dopar% {

  genotype<-as.character(genotypes_d[i,])
  genotype<-dplyr::recode(genotype, '0' = 'zero', '1' = 'one','2' = 'two')
  names(genotype)<-names(genotypes_d[i,])
  genotype<-genotype[!is.na(genotype)]
  genotype<-as.factor(genotype)
  merged_datasets_outlier_removal_b<-merged_datasets_outlier_removal_a[,names(genotype)]

  design<-model.matrix(~ 0 + genotype)
  colnames(design)<-c('zero','one','two')

  contrasts<-makeContrasts( zero - one,
                            zero - two,
                            one - two, levels=design)

  eqtl_edger_results<-DGEList(count=merged_datasets_outlier_removal_b, group=genotype, n
orm.factors = calcNormFactors(merged_datasets_outlier_removal_b, method = "TMM"))
  eqtl_edger_results<-estimateDisp(eqtl_edger_results,design, robust=TRUE)

  eqtl_edger_results_QLFit <- glmQLFit(eqtl_edger_results, design,robust=TRUE)
  eqtl_edger_results_QLFit <- glmQLFTest(eqtl_edger_results_QLFit , contrast=contrasts)
  eqtl_edger_results_QLFit_edgeR_results_QLF<- topTags(eqtl_edger_results_QLFit, adjust.
method = "none", n=Inf)$table
  eqtl_edger_results_QLFit_edgeR_results_QLF$snp<-rownames(genotypes_d)[i]
  eqtl_edger_results_QLFit_edgeR_results_QLF<-eqtl_edger_results_QLFit_edgeR_results_QLF
[eqtl_edger_results_QLFit_edgeR_results_QLF$PValue< 5e-08,]
  eqtl_edger_results_QLFit_edgeR_results_QLF$gene<-rownames(eqtl_edger_results_QLFit_edg
eR_results_QLF)

  rm(eqtl_edger_results_QLFit,keep,merged_datasets_outlier_removal_b)

  data.frame(eqtl_edger_results_QLFit_edgeR_results_QLF)
}

)

#user  system elapsed
#  8.354   3.266 355.865

stopCluster(cl)

cl <- makeCluster(34)
registerDoParallel(cl)

system.time (
```

```
results_zero_vs_one_two<- foreach(i = seq(1:dim(genotypes_d)[1]) ,.combine = 'rbind', .i
norder=FALSE , .errorhandling="remove", .packages="edgeR",.verbose=FALSE ) %dopar% {

  genotype<-as.character(genotypes_d[i,])
  genotype<-dplyr::recode(genotype, '0' = 'zero', '1' = 'one_two','2' = 'one_two')
  names(genotype)<-names(genotypes_d[i,])
  genotype<-genotype[!is.na(genotype)]
  genotype<-as.factor(genotype)
  merged_datasets_outlier_removal_b<-merged_datasets_outlier_removal_a[,names(genotype)]
  keep<-rowSums( cpm(merged_datasets_outlier_removal_b) >= 1 ) >= 5
  merged_datasets_outlier_removal_b<-merged_datasets_outlier_removal_b[keep,]

  design <- model.matrix(~ 0 + genotype)

  eqtl_edger<-DGEList(count=merged_datasets_outlier_removal_b, group=genotype, norm.fact
ors = calcNormFactors(merged_datasets_outlier_removal_b, method = "TMM"))
  eqtl_edger<-estimateDisp(eqtl_edger,design, robust=TRUE)
  eqtl_edger_QLFit <- glmQLFit(eqtl_edger, design,robust=TRUE)

  pairwise_contrast <- makeContrasts(contrast_1 = genotypezero - genotypeone_two, levels
=design)

  eqtl_edger_QLF <- glmQLFTest(eqtl_edger_QLFit, contrast= pairwise_contrast[,'contrast_
1'])
  eqtl_edger_results_QLF<- topTags(eqtl_edger_QLF, adjust.method = "none", n=Inf)$table
  eqtl_edger_results_QLF$snp<-rownames(genotypes_d)[i]
  eqtl_edger_results_QLF<-eqtl_edger_results_QLF[eqtl_edger_results_QLF$PValue< 5e-08,]
  eqtl_edger_results_QLF$gene<-rownames(eqtl_edger_results_QLF)

  rm(eqtl_edger_QLF,keep,merged_datasets_outlier_removal_b)

  data.frame(eqtl_edger_results_QLF)
}
)
stopCluster(cl)

#   user  system elapsed
#  8.855   4.074 352.567

cl <- makeCluster(34)
registerDoParallel(cl)

system.time (
results_zero_one_vs_two<- foreach(i = seq(1:dim(genotypes_d)[1]) ,.combine = 'rbind', .i
norder=FALSE , .errorhandling="remove", .packages="edgeR",.verbose=FALSE ) %dopar% {

  genotype<-as.character(genotypes_d[i,])
  genotype<-dplyr::recode(genotype, '0' = 'zero_one', '1' = 'zero_one','2' = 'two')
  names(genotype)<-names(genotypes_d[i,])
  genotype<-genotype[!is.na(genotype)]
  genotype<-as.factor(genotype)
  merged_datasets_outlier_removal_b<-merged_datasets_outlier_removal_a[,names(genotype)]
```

```
  keep<-rowSums( cpm(merged_datasets_outlier_removal_b) >= 1 ) >= 5
  merged_datasets_outlier_removal_b<-merged_datasets_outlier_removal_b[keep,]

  design <- model.matrix(~ 0 + genotype)

  eqtl_edger<-DGEList(count=merged_datasets_outlier_removal_b, group=genotype, norm.fact
ors = calcNormFactors(merged_datasets_outlier_removal_b, method = "TMM"))
  eqtl_edger<-estimateDisp(eqtl_edger,design, robust=TRUE)
  eqtl_edger_QLFit <- glmQLFit(eqtl_edger, design,robust=TRUE)

  pairwise_contrast <- makeContrasts(contrast_1 = genotypetwo - genotypezero_one, levels
=design)

  eqtl_edger_QLF <- glmQLFTest(eqtl_edger_QLFit, contrast= pairwise_contrast[,'contrast_
1'])
  eqtl_edger_results_QLF<- topTags(eqtl_edger_QLF, adjust.method = "none", n=Inf)$table
  eqtl_edger_results_QLF$snp<-rownames(genotypes_d)[i]
  eqtl_edger_results_QLF<-eqtl_edger_results_QLF[eqtl_edger_results_QLF$PValue< 5e-08,]
  eqtl_edger_results_QLF$gene<-rownames(eqtl_edger_results_QLF)

  rm(eqtl_edger_QLF,keep,merged_datasets_outlier_removal_b)

  data.frame(eqtl_edger_results_QLF)
}
)
stopCluster(cl)

#  user  system elapsed
#  7.625   2.028 353.194

cl <- makeCluster(34)
registerDoParallel(cl)
system.time (
results_one_vs_two<- foreach(i = seq(1:dim(genotypes_d)[1]) ,.combine = 'rbind', .inorde
r=FALSE , .errorhandling="remove", .packages="edgeR",.verbose=FALSE ) %dopar% {

  genotype<-as.character(genotypes_d[i,])
  genotype<-dplyr::recode(genotype, '0' = 'zero', '1' = 'one','2' = 'two')
  names(genotype)<-names(genotypes_d[i,])
  genotype<-genotype[!is.na(genotype)]
  genotype<-as.factor(genotype)
  merged_datasets_outlier_removal_b<-merged_datasets_outlier_removal_a[,names(genotype)]
  keep<-rowSums( cpm(merged_datasets_outlier_removal_b) >= 1 ) >= 5
  merged_datasets_outlier_removal_b<-merged_datasets_outlier_removal_b[keep,]

  design <- model.matrix(~ 0 + genotype)

  eqtl_edger<-DGEList(count=merged_datasets_outlier_removal_b, group=genotype, norm.fact
ors = calcNormFactors(merged_datasets_outlier_removal_b, method = "TMM"))
  eqtl_edger<-estimateDisp(eqtl_edger,design, robust=TRUE)
  eqtl_edger_QLFit <- glmQLFit(eqtl_edger, design,robust=TRUE)
```

```
  pairwise_contrast <- makeContrasts(contrast_1 = genotypeone - genotypetwo, levels=desi
gn)

  eqtl_edger_QLF <- glmQLFTest(eqtl_edger_QLFit, contrast= pairwise_contrast[,'contrast_
1'])
  eqtl_edger_results_QLF<- topTags(eqtl_edger_QLF, adjust.method = "none", n=Inf)$table
  eqtl_edger_results_QLF$snp<-rownames(genotypes_d)[i]
  eqtl_edger_results_QLF<-eqtl_edger_results_QLF[eqtl_edger_results_QLF$PValue< 5e-08,]
  eqtl_edger_results_QLF$gene<-rownames(eqtl_edger_results_QLF)

  rm(eqtl_edger_QLF,keep,merged_datasets_outlier_removal_b)

  data.frame(eqtl_edger_results_QLF)
}
)
stopCluster(cl)

#   user  system elapsed
#  7.548   2.138 360.063

cl <- makeCluster(34)
registerDoParallel(cl)
system.time (
results_zero_vs_one<- foreach(i = seq(1:dim(genotypes_d)[1]) ,.combine = 'rbind', .inord
er=FALSE , .errorhandling="remove", .packages="edgeR",.verbose=FALSE ) %dopar% {

  genotype<-as.character(genotypes_d[i,])
  genotype<-dplyr::recode(genotype, '0' = 'zero', '1' = 'one','2' = 'two')
  names(genotype)<-names(genotypes_d[i,])
  genotype<-genotype[!is.na(genotype)]
  genotype<-as.factor(genotype)
  merged_datasets_outlier_removal_b<-merged_datasets_outlier_removal_a[,names(genotype)]
  keep<-rowSums( cpm(merged_datasets_outlier_removal_b) >= 1 ) >= 5
  merged_datasets_outlier_removal_b<-merged_datasets_outlier_removal_b[keep,]

  design <- model.matrix(~ 0 + genotype)

  eqtl_edger<-DGEList(count=merged_datasets_outlier_removal_b, group=genotype, norm.fact
ors = calcNormFactors(merged_datasets_outlier_removal_b, method = "TMM"))
  eqtl_edger<-estimateDisp(eqtl_edger,design, robust=TRUE)
  eqtl_edger_QLFit <- glmQLFit(eqtl_edger, design,robust=TRUE)

  pairwise_contrast <- makeContrasts(contrast_1 = genotypezero - genotypeone, levels=des
ign)

  eqtl_edger_QLF <- glmQLFTest(eqtl_edger_QLFit, contrast= pairwise_contrast[,'contrast_
1'])
  eqtl_edger_results_QLF<- topTags(eqtl_edger_QLF, adjust.method = "none", n=Inf)$table
  eqtl_edger_results_QLF$snp<-rownames(genotypes_d)[i]
  eqtl_edger_results_QLF<-eqtl_edger_results_QLF[eqtl_edger_results_QLF$PValue< 5e-08,]
  eqtl_edger_results_QLF$gene<-rownames(eqtl_edger_results_QLF)
```

```
  rm(eqtl_edger_QLF,keep,merged_datasets_outlier_removal_b)

  data.frame(eqtl_edger_results_QLF)
}
)

#  user  system elapsed
#  6.889   2.027 358.571

stopCluster(cl)

cl <- makeCluster(34)
registerDoParallel(cl)
system.time (
results_zero_vs_two<- foreach(i = seq(1:dim(genotypes_d)[1]) ,.combine = 'rbind', .inord
er=FALSE , .errorhandling="remove", .packages="edgeR",.verbose=FALSE ) %dopar% {

  genotype<-as.character(genotypes_d[i,])
  genotype<-dplyr::recode(genotype, '0' = 'zero', '1' = 'one','2' = 'two')
  names(genotype)<-names(genotypes_d[i,])
  genotype<-genotype[!is.na(genotype)]
  genotype<-as.factor(genotype)
  merged_datasets_outlier_removal_b<-merged_datasets_outlier_removal_a[,names(genotype)]
  keep<-rowSums( cpm(merged_datasets_outlier_removal_b) >= 1 ) >= 5
  merged_datasets_outlier_removal_b<-merged_datasets_outlier_removal_b[keep,]

  design <- model.matrix(~ 0 + genotype)

  eqtl_edger<-DGEList(count=merged_datasets_outlier_removal_b, group=genotype, norm.fact
ors = calcNormFactors(merged_datasets_outlier_removal_b, method = "TMM"))
  eqtl_edger<-estimateDisp(eqtl_edger,design, robust=TRUE)
  eqtl_edger_QLFit <- glmQLFit(eqtl_edger, design,robust=TRUE)

  pairwise_contrast <- makeContrasts(contrast_1 = genotypezero - genotypetwo, levels=des
ign)

  eqtl_edger_QLF <- glmQLFTest(eqtl_edger_QLFit, contrast= pairwise_contrast[,'contrast_
1'])
  eqtl_edger_results_QLF<- topTags(eqtl_edger_QLF, adjust.method = "none", n=Inf)$table
  eqtl_edger_results_QLF$snp<-rownames(genotypes_d)[i]
  eqtl_edger_results_QLF<-eqtl_edger_results_QLF[eqtl_edger_results_QLF$PValue< 5e-08,]
  eqtl_edger_results_QLF$gene<-rownames(eqtl_edger_results_QLF)

  rm(eqtl_edger_QLF,keep,merged_datasets_outlier_removal_b)

  data.frame(eqtl_edger_results_QLF)
}
)
stopCluster(cl)

#   user  system elapsed
#  7.195   2.121 359.844
```

```
dim(dplyr::anti_join(results_zero_vs_one_two, results_one_vs_two, by=c("snp","gene")))

dim(dplyr::anti_join(results_zero_one_vs_two, results_zero_vs_one, by=c("snp","gene")))

results_zero_vs_one_two_a<-dplyr::anti_join(results_zero_vs_one_two, results_one_vs_two,
by=c("snp","gene"))

results_zero_one_vs_two_a<-dplyr::anti_join(results_zero_one_vs_two, results_zero_vs_on
e, by=c("snp","gene"))

results_dominance<-rbind(results_zero_vs_one_two_a,results_zero_one_vs_two_a)

results_anova_dominance <- merge(results_anova, results_dominance,  by.x=c("snp","gene"
), by.y= c("snp", "gene"), suffix=c(".anova",".contrast" ), all=FALSE)

#write.table(results_anova_dominance, file= "/mnt/storage/lab_folder/shared_R_codes/fern
ando/SNP_eqtl/results/2022_11_05_results_anova_dominance.txt", append = FALSE, quote = F
ALSE, sep = "\t" ,row.names = FALSE)
```

#combined dominance annotation

```
results_anova_dominance<-read.delim("/mnt/storage/lab_folder/shared_R_codes/fernando/SNP
_eqtl/results/2022_11_05_results_anova_dominance.txt", row.names=NULL, header = TRUE)
eqtl_annotate_anova_edgeR1 <- results_anova_dominance
#eqtl_annotate_anova_edgeR1<-eqtl_annotate_anova_edgeR[eqtl_annotate_anova_edgeR[,4] < 5
e-08,]
eqtl_annotate_anova_edgeR1<-merge(eqtl_annotate_anova_edgeR1, merged_SNPS_nucleotide, b
y.x= "snp", by.y= "row.names", all=FALSE)
eqtl_annotate_anova_edgeR1<-merge(eqtl_annotate_anova_edgeR1, genotypes_d1,by.x= c("snp"
), by.y= 'row.names', all=FALSE)
eqtl_annotate_anova_edgeR1<-merge(eqtl_annotate_anova_edgeR1, annotation.ensembl.symbol,
by.x="gene", by.y="ensembl_gene_id", all.x=TRUE, all.y=FALSE)
eqtl_annotate_anova_edgeR1<-merge(eqtl_annotate_anova_edgeR1, SNP_annotation_all_a, by.x
="snp", by.y="SNP", all=FALSE)
eqtl_annotate_anova_edgeR1<-eqtl_annotate_anova_edgeR1[order(eqtl_annotate_anova_edgeR1
$PValue.contrast ),]
eqtl_annotate_anova_edgeR1<-eqtl_annotate_anova_edgeR1[!duplicated(eqtl_annotate_anova_e
dgeR1[,c(1:2)]),]

#write.table(eqtl_annotate_anova_edgeR1, file= "/mnt/storage/lab_folder/shared_R_codes/f
ernando/SNP_eqtl/results/Supplementary_table_6.txt", append = FALSE, quote = FALSE, sep
 = "\t" ,row.names = FALSE)
```

```
eqtl_annotate_anova_edgeR1_taff15_a<- eqtl_annotate_anova_edgeR1[eqtl_annotate_anova_edg
eR1$snp=="19:14551828",]

snps_sub2 = eqtl_annotate_anova_edgeR1_taff15_a$snp[1:6]
genes_sub2 = eqtl_annotate_anova_edgeR1_taff15_a$gene[1:6]
gene_symbol = eqtl_annotate_anova_edgeR1_taff15_a$external_gene_name[1:6]


plot_list_1<-list()

for (index in seq(length(snps_sub2))){
  genotype_sub2 = unlist((merged_SNPS_nucleotide[snps_sub2[index],c(3:44)]))
  genotype_sub2 = genotype_sub2[!(genotype_sub2=="<NA>")]
  expression_sub2 = tpm_merged_datasets[genes_sub2[index],]
  expression_sub2 = expression_sub2[names(expression_sub2) %in% names(genotype_sub2) ]
  genotype_sub2<-genotype_sub2[names(expression_sub2)]

  graph_data_frame<-data.frame(genotype_sub2=gsub("/","",genotype_sub2),expression_sub2,
snp=snps_sub2[index], gene_symbol=gene_symbol[index])
  graph_data_frame<-graph_data_frame[complete.cases(graph_data_frame),]
  graph_data_frame<-graph_data_frame[!(graph_data_frame$genotype_sub2 =="<NA>"),]

  plot_list_1[[index]]<-ggplot(data=graph_data_frame, aes(x=as.factor(genotype_sub2), y=
expression_sub2))+
    geom_boxplot(fill='transparent',outlier.shape = 4, outlier.color = "blue", size=0.1)
+
    geom_jitter(width=0.2, size=1)+
    scale_x_discrete(name= graph_data_frame$snp[1])+
    scale_y_continuous(name =  graph_data_frame$gene_symbol[1])+
    theme_classic(base_size = 18)+
    theme( axis.text=element_text(size=18,color="black"),
           axis.title=element_text(size=18,color="black"))
}
Figure_6_B_a<-plot_grid( plotlist = plot_list_1, nrow = 2)


eqtl_annotate_anova_edgeR1_taff15_b<- eqtl_annotate_anova_edgeR1[eqtl_annotate_anova_edg
eR1$snp=="19:14554403",]

snps_sub2 = eqtl_annotate_anova_edgeR1_taff15_b$snp[1:6]
genes_sub2 = eqtl_annotate_anova_edgeR1_taff15_b$gene[1:6]
gene_symbol = eqtl_annotate_anova_edgeR1_taff15_b$external_gene_name[1:6]


plot_list_2<-list()

for (index in seq(length(snps_sub2))){
  genotype_sub2 = unlist((merged_SNPS_nucleotide[snps_sub2[index],c(3:44)]))
  genotype_sub2 = genotype_sub2[!(genotype_sub2=="<NA>")]
  expression_sub2 = tpm_merged_datasets[genes_sub2[index],]
  expression_sub2 = expression_sub2[names(expression_sub2) %in% names(genotype_sub2) ]
  genotype_sub2<-genotype_sub2[names(expression_sub2)]
```

```
  graph_data_frame<-data.frame(genotype_sub2=gsub("/","",genotype_sub2),expression_sub2,
snp=snps_sub2[index], gene_symbol=gene_symbol[index])
  graph_data_frame<-graph_data_frame[complete.cases(graph_data_frame),]
  graph_data_frame<-graph_data_frame[!(graph_data_frame$genotype_sub2 =="<NA>"),]

  plot_list_2[[index]]<-ggplot(data=graph_data_frame, aes(x=as.factor(genotype_sub2), y=
expression_sub2))+
    geom_boxplot(fill='transparent',outlier.shape = 4, outlier.color = "blue", size=0.1)
+
    geom_jitter(width=0.2, size=1)+
    scale_x_discrete(name= graph_data_frame$snp[1])+
    scale_y_continuous(name =  graph_data_frame$gene_symbol[1])+
    theme_classic(base_size = 18)+
    theme( axis.text=element_text(size=18,color="black"),
           axis.title=element_text(size=18,color="black"))
}
Figure_6_B_b<-plot_grid( plotlist = plot_list_2, nrow = 2)


plot_grid(Figure_6_B_a,NULL,Figure_6_B_b, nrow=1, rel_widths=c(1, 0.1, 1))
```

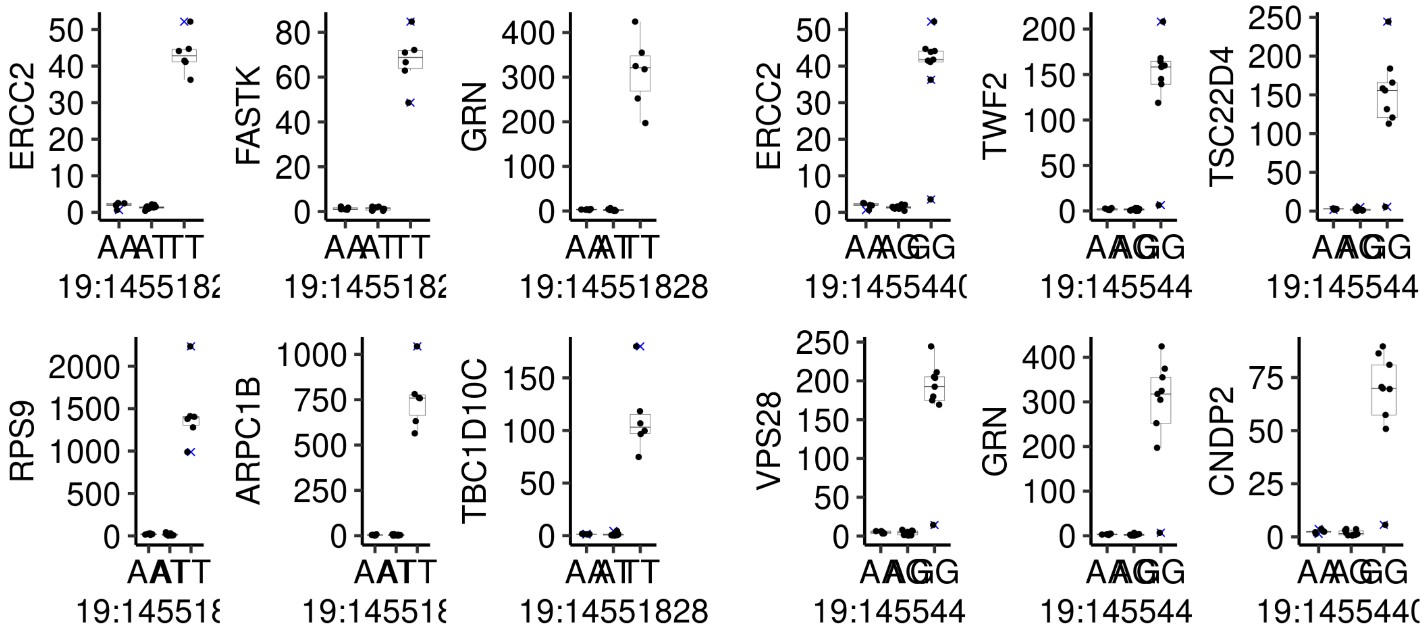


#Gene ontology enrichment analysis

```
annotation.ensembl.symbol<-read.delim("/mnt/storage/lab_folder/shared_R_codes/fernando/R
NA_degradation/resources/2021_02_19_annotation.ensembl.symbol.txt.bz2", header=TRUE, sep
= "\t",row.names=1, stringsAsFactors = FALSE)
gene.length<-read.delim("/mnt/storage/lab_folder/shared_R_codes/fernando/RNA_degradatio
n/resources/2021_02_19_gene.length.txt.bz2", header=TRUE, sep= "\t",row.names=1, strings
AsFactors = FALSE)
annotation.GO.biomart<-read.delim("/mnt/storage/lab_folder/shared_R_codes/fernando/RNA_d
egradation/resources/2021_02_19_annotation.GO.biomart.txt.bz2", header=TRUE, sep= "\t",r
ow.names=1, stringsAsFactors = FALSE)
annotation.ensembl.transcript<-read.delim("/mnt/storage/lab_folder/shared_R_codes/fernan
do/RNA_degradation/resources/2021_02_19_annotation.ensembl.transcript.txt.bz2", header=T
RUE, sep= "\t",row.names=1, stringsAsFactors = FALSE)


all_genes<-data.frame( gene=row.names(merged_datasets), stringsAsFactors=FALSE )
rownames(all_genes)<-all_genes$gene
N_expressed_genes<-length(all_genes$gene)

gene.length<-gene.length[gene.length$ensembl_gene_id %in% all_genes$gene,]

annotation.genelength.biomart_vector<-gene.length$transcript_length
names(annotation.genelength.biomart_vector)<-gene.length$ensembl_gene_id

annotation.GO.BP.biomart<-annotation.GO.biomart[annotation.GO.biomart$namespace_1003=="b
iological_process", c(1,3)]
annotation.GO.BP.biomart<-annotation.GO.BP.biomart[annotation.GO.BP.biomart$ensembl_gene
_id %in% rownames(all_genes),]
annotation.GO.MF.biomart<-annotation.GO.biomart[annotation.GO.biomart$namespace_1003=="m
olecular_function", c(1,3)]
annotation.GO.MF.biomart<-annotation.GO.MF.biomart[annotation.GO.MF.biomart$ensembl_gene
_id %in% rownames(all_genes),]

test.genes<-data.frame(a=unique(eqtl_annotate_anova_edgeR1[eqtl_annotate_anova_edgeR1$PV
alue.contrast < 1e-12,]$gene), stringsAsFactors=FALSE)

all_genes_numeric<-as.integer(all_genes$gene %in%test.genes$a)
names(all_genes_numeric)<-all_genes$gene

N_sig_genes<-length(test.genes$a)

#N_sig_genes

set.seed(9830)
pwf<-nullp(all_genes_numeric, bias.data=annotation.genelength.biomart_vector, plot.fit=F
ALSE )
GO_BP_Cats_raw_counts<-goseq(pwf,gene2cat=annotation.GO.BP.biomart, method ="Sampling",
 repcnt = 5000, use_genes_without_cat=FALSE)
GO_BP_Cats_raw_counts<-GO_BP_Cats_raw_counts[GO_BP_Cats_raw_counts$numDEInCat>3,]
GO_BP_Cats_raw_counts$FWER<-p.adjust(GO_BP_Cats_raw_counts$over_represented_pvalue, meth
od ="holm")
GO_BP_Cats_raw_counts<-GO_BP_Cats_raw_counts[with(GO_BP_Cats_raw_counts, order(FWER,over
_represented_pvalue, -numDEInCat)), ]
```

```
#head(GO_BP_Cats_raw_counts, n=20)

GO_BP_Cats_raw_counts$fold_enrichment<-(GO_BP_Cats_raw_counts$numDEInCat/N_sig_genes)/(GO_BP_Cats_raw_counts$numInCat/N_expressed_genes)
annotation.GO.BP.biomart_testgenes<-annotation.GO.BP.biomart[annotation.GO.BP.biomart$ensembl_gene_id %in% test.genes$a, ]
GO_BP_Cats_raw_counts<-merge(GO_BP_Cats_raw_counts,annotation.GO.BP.biomart_testgenes, by.x="category", by.y="go_id", all.x=TRUE, all.y=FALSE)
GO_BP_Cats_raw_counts<-merge(GO_BP_Cats_raw_counts, annotation.ensembl.symbol, by.x="ensembl_gene_id", by.y="ensembl_gene_id", all=FALSE, all.x=TRUE, all.y=FALSE)
GO_BP_Cats_raw_counts<-GO_BP_Cats_raw_counts[with(GO_BP_Cats_raw_counts, order(FWER,term)), ]
GO_BP_Cats_raw_counts<-GO_BP_Cats_raw_counts[GO_BP_Cats_raw_counts$FWER<0.1,]

#write.table(GO_BP_Cats_raw_counts, file= "/mnt/storage/lab_folder/shared_R_codes/fernando/SNP_eqtl/results/Supplementary_table_7.txt", append = FALSE, quote = FALSE, sep = "\t" ,row.names = FALSE)
```

#EdgeR linear eqtl analysis

```
#rm(linear_test)
cl <- makeCluster(34)
registerDoParallel(cl)
system.time(
linear_test<- foreach(i = seq(1:dim(genotypes_d)[1]) ,.combine = 'rbind', .inorder=FALSE
, .errorhandling="remove", .packages="edgeR",.verbose=FALSE ) %dopar% {

  genotype<-genotypes_d[i,]
  names(genotype)<-names(genotypes_d[i,])
  genotype<-genotype[!is.na(genotype)]
  merged_datasets_outlier_removal_b<-merged_datasets_outlier_removal_a[,names(genotype)]
  keep<-rowSums( cpm(merged_datasets_outlier_removal_b) >= 1 ) >= 5
  merged_datasets_outlier_removal_b<-merged_datasets_outlier_removal_b[keep,]

  design <- model.matrix(~ genotype)

  eqtl_edger<-DGEList(count=merged_datasets_outlier_removal_b, group=genotype, norm.fact
ors = calcNormFactors(merged_datasets_outlier_removal_b, method = "TMM"))
  eqtl_edger<-estimateDisp(eqtl_edger,design, robust=TRUE)
  eqtl_edger_QLFit <- glmQLFit(eqtl_edger, design,robust=TRUE)

  eqtl_edger_QLF <- glmQLFTest(eqtl_edger_QLFit, coef = 2)
  eqtl_edger_results_QLF<- topTags(eqtl_edger_QLF, adjust.method = "none", n=Inf)$table
  eqtl_edger_results_QLF$snp<-rownames(genotypes_d)[i]
  eqtl_edger_results_QLF<-eqtl_edger_results_QLF[eqtl_edger_results_QLF$PValue< 5e-08,]
  eqtl_edger_results_QLF$gene<-rownames(eqtl_edger_results_QLF)

  rm(eqtl_edger_QLF,keep,merged_datasets_outlier_removal_b)

  data.frame(eqtl_edger_results_QLF)
}
)
stopCluster(cl)

#   user  system elapsed
#  6.705   1.764 517.853

#write.table(linear_test, file= "/mnt/storage/lab_folder/shared_R_codes/fernando/SNP_eqt
l/results/2022_11_05_linear_test.txt", append = FALSE, quote = FALSE, sep = "\t" ,row.na
mes = FALSE)
```

#EdgeR linear annotation

```
linear_test<-read.delim("/mnt/storage/lab_folder/shared_R_codes/fernando/SNP_eqtl/results/2022_11_05_linear_test.txt", row.names=NULL, header = TRUE)

linear_test_a<-linear_test
linear_test_a<-linear_test_a[!(linear_test_a$snp %in% c("19:14554927","19:14551828", "17:63027414","16:2303900","19:23080893", "13:31660515","19:14554403", "26:49413229", "17:62945054")),]
eqtl_annotate_linear_edgeR1 <- linear_test_a
eqtl_annotate_linear_edgeR1<-merge(eqtl_annotate_linear_edgeR1, merged_SNPS_nucleotide, by.x= "snp", by.y= "row.names", all=FALSE)
eqtl_annotate_linear_edgeR1<-merge(eqtl_annotate_linear_edgeR1, genotypes_d1,by.x= c("snp"), by.y= 'row.names', all=FALSE)
eqtl_annotate_linear_edgeR1<-merge(eqtl_annotate_linear_edgeR1, annotation.ensembl.symbol, by.x="gene", by.y="ensembl_gene_id", all.x=TRUE, all.y=FALSE)
eqtl_annotate_linear_edgeR1<-merge(eqtl_annotate_linear_edgeR1, SNP_annotation_all_a, by.x="snp", by.y="SNP", all=FALSE)
eqtl_annotate_linear_edgeR1<-eqtl_annotate_linear_edgeR1[order(eqtl_annotate_linear_edgeR1$PValue ),]
eqtl_annotate_linear_edgeR1<-eqtl_annotate_linear_edgeR1[!duplicated(eqtl_annotate_linear_edgeR1[,c(1:2)]),]

#write.table(eqtl_annotate_linear_edgeR1, file= "/mnt/storage/lab_folder/shared_R_codes/fernando/SNP_eqtl/results/Supplementary_table_5.txt", append = FALSE, quote = FALSE, sep = "\t" ,row.names = FALSE)
```

#Figure 5A

```
eqtl_annotate_linear_edgeR1$external_gene_name[1]<-"SIGLEC14"
eqtl_annotate_linear_edgeR1$external_gene_name[2]<-"SIGLEC14"
eqtl_annotate_linear_edgeR1$external_gene_name[3]<-"SLC11A2"

eqtl_annotate_linear_edgeR1$external_gene_name<-ifelse(eqtl_annotate_linear_edgeR1$exter
nal_gene_name=="", eqtl_annotate_linear_edgeR1$gene,eqtl_annotate_linear_edgeR1$external
_gene_name)

snps_sub2 = eqtl_annotate_linear_edgeR1$snp
genes_sub2 = eqtl_annotate_linear_edgeR1$gene
gene_symbol = eqtl_annotate_linear_edgeR1$external_gene_name

plot_list<-list()

for (index in seq(length(snps_sub2))){
  genotype_sub2 = unlist((merged_SNPS_nucleotide[snps_sub2[index],c(3:44)]))
  genotype_sub2 = genotype_sub2[!(genotype_sub2=="<NA>")]
  expression_sub2 = tpm_merged_datasets[genes_sub2[index],]
  expression_sub2 = expression_sub2[names(expression_sub2) %in% names(genotype_sub2) ]
  genotype_sub2<-genotype_sub2[names(expression_sub2)]

  graph_data_frame<-data.frame(genotype_sub2=gsub("/","",genotype_sub2),expression_sub2,
snp=snps_sub2[index], gene_symbol=gene_symbol[index])
  graph_data_frame<-graph_data_frame[complete.cases(graph_data_frame),]
  graph_data_frame<-graph_data_frame[!(graph_data_frame$genotype_sub2 =="<NA>"),]

  plot_list[[index]]<-ggplot(data=graph_data_frame, aes(x=as.factor(genotype_sub2), y=ex
pression_sub2))+
    geom_boxplot(fill='transparent',outlier.shape = 4, outlier.color = "blue", size=0.1)
+
    geom_jitter(width=0.2, size=1)+
    scale_x_discrete(name= graph_data_frame$snp[1])+
    scale_y_continuous(name =  graph_data_frame$gene_symbol[1])+
    theme_classic(base_size = 18)+
    theme( axis.text=element_text(size=18,color="black"),
           axis.title=element_text(size=18,color="black"))
}
plot_grid( plotlist = plot_list , nrow = 4, ncol=2)
```

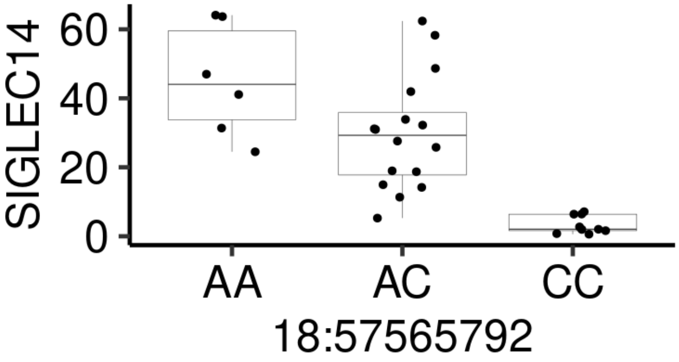
SIGLEC14
60
40
20
0
AA
AC
CC
18:57565792

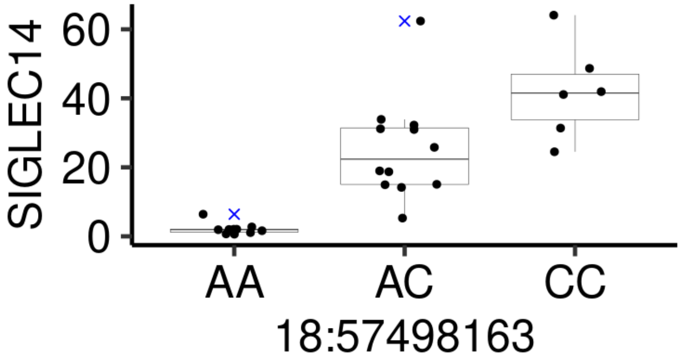
SIGLEC14
60
40
20
0
AA
AC
CC
18:57498163

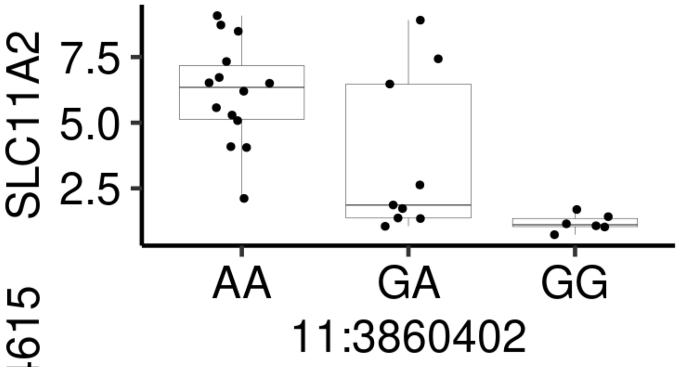
SLC11A2
7.5
5.0
2.5
AA
GA
GG
11:3860402

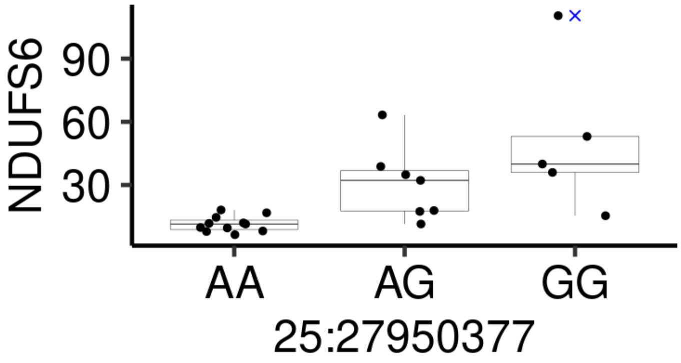
NDUFS6
90
60
30
AA
AG
GG
25:27950377

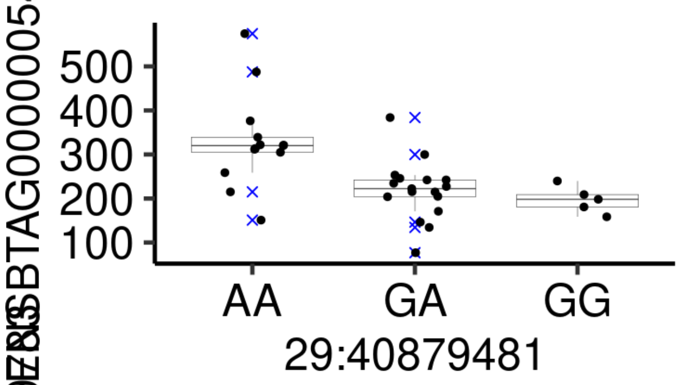
ENSBTAG00000054615
500
400
300
200
100
AA
GA
GG
29:40879481

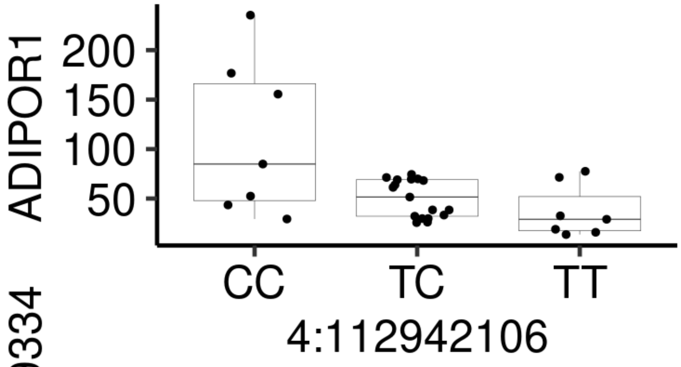
ADIPOR1
200
150
100
50
CC
TC
TT
4:112942106

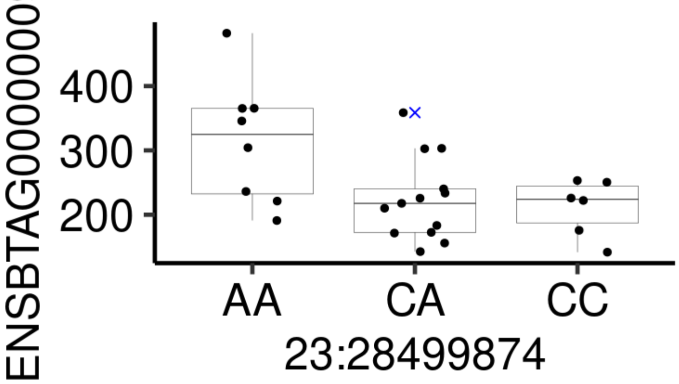
400
300
200
AA
CA
CC
23:28499874

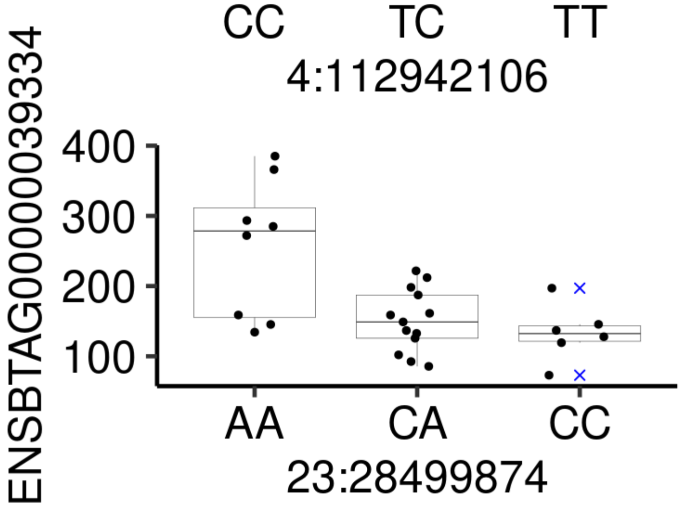
ENSBTAG00000039334
400
300
200
100
AA
CA
CC
23:28499874

```
results_anova_dominance<-dplyr::anti_join(results_anova_dominance, linear_test_a, by=c(
"snp","gene"))

results_anova_dominance_a<-results_anova_dominance[, c("logFC" ,"logCPM.contrast", "F.co
ntrast" ,"PValue.contrast", "snp","gene")]
colnames(results_anova_dominance_a)<-c("logFC","logCPM","F","PValue","snp","gene")
results_dominance_linear<-rbind(results_anova_dominance_a,linear_test_a)

eqtl_annotate_anova_linear_edgeR1 <- results_dominance_linear
#eqtl_annotate_anova_linear_edgeR1<-eqtl_annotate_anova_edgeR[eqtl_annotate_anova_edgeR
[,4] < 5e-08,]
eqtl_annotate_anova_linear_edgeR1<-merge(eqtl_annotate_anova_linear_edgeR1, merged_SNPS_
nucleotide, by.x= "snp", by.y= "row.names", all=FALSE)
eqtl_annotate_anova_linear_edgeR1<-merge(eqtl_annotate_anova_linear_edgeR1, genotypes_d
1,by.x= c("snp"), by.y= 'row.names', all=FALSE)
eqtl_annotate_anova_linear_edgeR1<-merge(eqtl_annotate_anova_linear_edgeR1, annotation.e
nsembl.symbol, by.x="gene", by.y="ensembl_gene_id", all.x=TRUE, all.y=FALSE)
eqtl_annotate_anova_linear_edgeR1<-merge(eqtl_annotate_anova_linear_edgeR1, SNP_annotati
on_all_a, by.x="snp", by.y="SNP", all=FALSE)
eqtl_annotate_anova_linear_edgeR1<-eqtl_annotate_anova_linear_edgeR1[!duplicated(eqtl_an
notate_anova_linear_edgeR1[,c(1:2)]),]
```

#gene network

```
for_network<-eqtl_annotate_anova_linear_edgeR1[,c("snp","external_gene_name")]
for_network<-for_network[!(for_network$external_gene_name==""),]
write.table(for_network,"/mnt/storage/lab_folder/shared_R_codes/fernando/SNP_eqtl/result
s/edger_network.txt", col.names = TRUE, row.names = TRUE, quote = FALSE,sep ="\t")
```

#Figure 4B

```
frequency_gene_SNP<-data.frame(table(eqtl_annotate_anova_linear_edgeR1$SYMBOL))
frequency_gene_SNP<-frequency_gene_SNP[!frequency_gene_SNP$Var1=="-",]
frequency_gene_SNP<-frequency_gene_SNP[order(frequency_gene_SNP$Freq, decreasing=TRUE),]
head(frequency_gene_SNP)
```

```
##       Var1 Freq
## 10   TAF15 2263
## 9     SMG6  213
## 12     VIM    6
## 5     FUT4    3
## 2    AHNAK    1
## 3  ALDH6A1    1
```

```
frequency_gene_SNP$Var1<-factor(frequency_gene_SNP$Var1, levels=frequency_gene_SNP$Var1)

p1 = ggplot(data=frequency_gene_SNP, aes(x=Var1,y=Freq))+
  geom_col(aes())+
  scale_x_discrete(name=NULL)+
  scale_y_continuous(name="Frequency")+
  #scale_fill_manual(values=c("black","#5c006f","#0072d3","#ff7a9b"), labels=c("other",
 "missense variant","3 prime UTR variant", "5 prime UTR variant" ))+
  theme_classic()+
  theme(
    axis.text.x = element_text(angle=90, color="black", face="italic", size=15, vjust=0.
5,hjust = 0),
    axis.text.y = element_text(color="black", size=15),
    axis.title=element_text(color="black", size=15),
    legend.position = c(0.6, 0.9),
    legend.title = element_blank(),
    legend.direction="horizontal"
  )+
  geom_rect(aes(xmin = "FUT4", xmax = "ZNF175", ymin = 0, ymax = 50), color = "black", a
lpha = 0)

frequency_gene_SNP_zoom<-frequency_gene_SNP[frequency_gene_SNP$Freq<5,]

p2 = ggplot(data=frequency_gene_SNP_zoom, aes(x=Var1,y=Freq))+
  geom_col(aes())+
  scale_x_discrete(name=NULL)+
  scale_y_continuous(name="Frequency")+
  #scale_fill_manual(values=c("black","#5c006f","#0072d3","#ff7a9b"))+
  labs(y = NULL, x = NULL) +
  theme_classic()+
  theme(
    axis.text.x = element_text(angle=90, color="black", face="italic", size=15, vjust=0.
5,hjust = 0),
    legend.position = "none")

p1 +
  annotation_custom(grob = ggplotGrob(p2), xmin = "FUT4", xmax = "ZNF175" ,ymin = 700, y
max = 2300)+
  geom_rect(aes(xmin = "FUT4", xmax = "ZNF175", ymin = 700, ymax = 2300), color='black',
linetype='dashed', alpha=0) +
  geom_path(aes(x,y,group=grp),
            data=data.frame(x = c("FUT4","FUT4","ZNF175","ZNF175"), y=c(20,700,20,700),g
rp=c(1,1,2,2)),
            linetype='dashed')
```

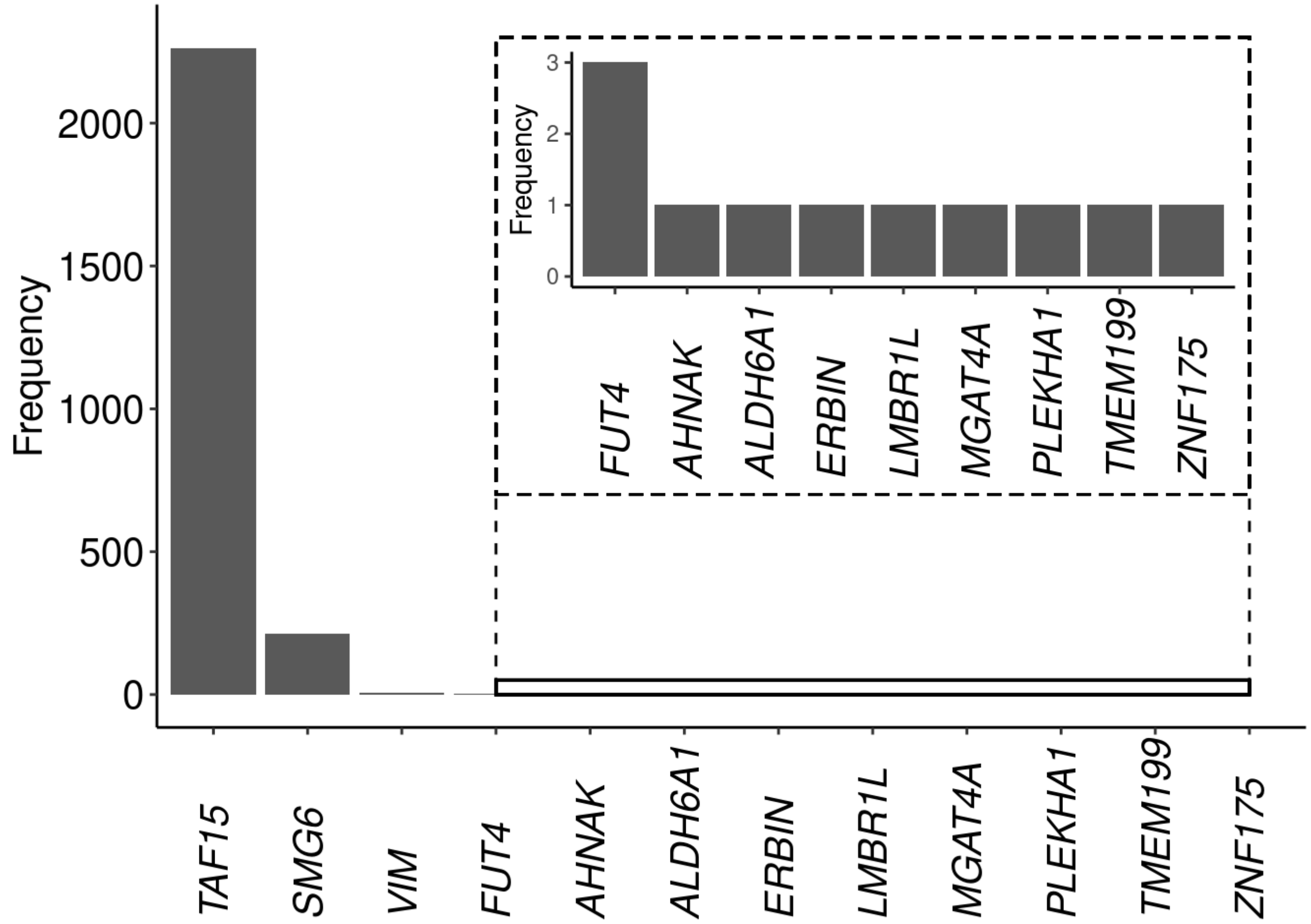


#Supplementary table 4

```
colnames(eqtl_annotate_anova_TMM)<-c("SNP","gene","F-test","pvalue" , "q.values", "bonfe
rroni","FDR")

head(eqtl_annotate_anova_edgeR1)
```

```
##             snp               gene logFC.zero...one logFC.zero...two
## 35  19:14551828 ENSBTAG00000002072        -3.567393      -0.26226496
## 464 19:14551828 ENSBTAG00000011228        -4.496188      -0.08387129
## 340 19:14551828 ENSBTAG00000018823        -5.734614      -0.42314519
## 69  19:14551828 ENSBTAG00000006487        -4.965559      -0.03418299
## 495 19:14551828 ENSBTAG00000046248        -6.185310       0.13041926
## 679 19:14551828 ENSBTAG00000013133        -5.304771      -0.28023463
##     logFC.one...two logCPM.anova  F.anova PValue.anova    logFC logCPM.contrast
## 35         3.305128     5.826073 195.0496 1.456671e-15 3.475309        5.825912
## 464        4.412317     5.234047 186.6708 2.976182e-15 4.466365        5.233777
## 340        5.311469     7.882550 187.9597 2.954420e-15 5.580285        7.882495
## 69         4.931376     9.103754 174.3559 6.808174e-15 4.954641        9.103731
## 495        6.315729     8.317532 173.2688 7.296904e-15 6.226633        8.317490
## 679        5.024536     6.417904 177.2604 5.668229e-15 5.200606        6.417764
##     F.contrast PValue.contrast CHROM      POS SL220764.x SL220765.x SL220766.x
## 35    382.2776    1.209050e-16    19 14551828        T/T       <NA>        T/T
## 464   387.7560    1.227991e-16    19 14551828        T/T       <NA>        T/T
## 340   366.2399    2.646750e-16    19 14551828        T/T       <NA>        T/T
## 69    362.3687    2.993080e-16    19 14551828        T/T       <NA>        T/T
## 495   358.5646    3.381598e-16    19 14551828        T/T       <NA>        T/T
## 679   358.2817    3.412592e-16    19 14551828        T/T       <NA>        T/T
##     SL220767.x SL220768.x SL220769.x SL220770.x SL220771.x SL220772.x
## 35        <NA>       <NA>        T/T       <NA>       <NA>       <NA>
## 464       <NA>       <NA>        T/T       <NA>       <NA>       <NA>
## 340       <NA>       <NA>        T/T       <NA>       <NA>       <NA>
## 69        <NA>       <NA>        T/T       <NA>       <NA>       <NA>
## 495       <NA>       <NA>        T/T       <NA>       <NA>       <NA>
## 679       <NA>       <NA>        T/T       <NA>       <NA>       <NA>
##     SL220773.x SL220774.x SL220775.x SL253803.x SL253804.x SL253805.x
## 35        <NA>       <NA>       <NA>        A/T        T/T        A/T
## 464       <NA>       <NA>       <NA>        A/T        T/T        A/T
## 340       <NA>       <NA>       <NA>        A/T        T/T        A/T
## 69        <NA>       <NA>       <NA>        A/T        T/T        A/T
## 495       <NA>       <NA>       <NA>        A/T        T/T        A/T
## 679       <NA>       <NA>       <NA>        A/T        T/T        A/T
##     SL253806.x SL253807.x SL253808.x SL253809.x SL253810.x SL253811.x
## 35         A/A        A/T        T/T        A/T        A/T        T/T
## 464        A/A        A/T        T/T        A/T        A/T        T/T
## 340        A/A        A/T        T/T        A/T        A/T        T/T
## 69         A/A        A/T        T/T        A/T        A/T        T/T
## 495        A/A        A/T        T/T        A/T        A/T        T/T
## 679        A/A        A/T        T/T        A/T        A/T        T/T
##     SL253812.x SL253813.x SL253814.x SL297951.x SL297952.x SL297953 SL297954.x
## 35         A/T        A/A       <NA>        A/A       <NA>     <NA>        A/A
## 464        A/T        A/A       <NA>        A/A       <NA>     <NA>        A/A
## 340        A/T        A/A       <NA>        A/A       <NA>     <NA>        A/A
## 69         A/T        A/A       <NA>        A/A       <NA>     <NA>        A/A
## 495        A/T        A/A       <NA>        A/A       <NA>     <NA>        A/A
## 679        A/T        A/A       <NA>        A/A       <NA>     <NA>        A/A
##     SL297955.x SL297956.x SL297957.x SL297958.x SL297959.x SL297960.x
## 35         A/T       <NA>       <NA>        A/T        A/T        A/T
## 464        A/T       <NA>       <NA>        A/T        A/T        A/T
```

```
## 340        A/T       <NA>       <NA>        A/T        A/T        A/T
## 69         A/T       <NA>       <NA>        A/T        A/T        A/T
## 495        A/T       <NA>       <NA>        A/T        A/T        A/T
## 679        A/T       <NA>       <NA>        A/T        A/T        A/T
##     SL297961.x SL297962.x SL297963.x SL297964.x SL297965.x SL297966.x
## 35        <NA>       <NA>        A/T       <NA>       <NA>       <NA>
## 464       <NA>       <NA>        A/T       <NA>       <NA>       <NA>
## 340       <NA>       <NA>        A/T       <NA>       <NA>       <NA>
## 69        <NA>       <NA>        A/T       <NA>       <NA>       <NA>
## 495       <NA>       <NA>        A/T       <NA>       <NA>       <NA>
## 679       <NA>       <NA>        A/T       <NA>       <NA>       <NA>
##     SL297967.x SL297968.x SL297951.y SL297952.y SL297954.y SL297955.y
## 35        <NA>        A/A          0         NA          0          1
## 464       <NA>        A/A          0         NA          0          1
## 340       <NA>        A/A          0         NA          0          1
## 69        <NA>        A/A          0         NA          0          1
## 495       <NA>        A/A          0         NA          0          1
## 679       <NA>        A/A          0         NA          0          1
##     SL297956.y SL297957.y SL297958.y SL297959.y SL297960.y SL297961.y
## 35          NA         NA          1          1          1         NA
## 464         NA         NA          1          1          1         NA
## 340         NA         NA          1          1          1         NA
## 69          NA         NA          1          1          1         NA
## 495         NA         NA          1          1          1         NA
## 679         NA         NA          1          1          1         NA
##     SL297962.y SL297963.y SL297964.y SL297965.y SL297966.y SL297967.y
## 35          NA          1         NA         NA         NA         NA
## 464         NA          1         NA         NA         NA         NA
## 340         NA          1         NA         NA         NA         NA
## 69          NA          1         NA         NA         NA         NA
## 495         NA          1         NA         NA         NA         NA
## 679         NA          1         NA         NA         NA         NA
##     SL297968.y SL220764.y SL220765.y SL220766.y SL220767.y SL220768.y
## 35           0          2         NA          2         NA         NA
## 464          0          2         NA          2         NA         NA
## 340          0          2         NA          2         NA         NA
## 69           0          2         NA          2         NA         NA
## 495          0          2         NA          2         NA         NA
## 679          0          2         NA          2         NA         NA
##     SL220769.y SL220770.y SL220771.y SL220772.y SL220773.y SL220774.y
## 35           2         NA         NA         NA         NA         NA
## 464          2         NA         NA         NA         NA         NA
## 340          2         NA         NA         NA         NA         NA
## 69           2         NA         NA         NA         NA         NA
## 495          2         NA         NA         NA         NA         NA
## 679          2         NA         NA         NA         NA         NA
##     SL220775.y SL253803.y SL253804.y SL253805.y SL253806.y SL253807.y
## 35          NA          1          2          1          0          1
## 464         NA          1          2          1          0          1
## 340         NA          1          2          1          0          1
## 69          NA          1          2          1          0          1
## 495         NA          1          2          1          0          1
```

```
## 679         NA          1          2          1          0          1
##     SL253808.y SL253809.y SL253810.y SL253811.y SL253812.y SL253813.y
## 35           2          1          1          2          1          0
## 464          2          1          1          2          1          0
## 340          2          1          1          2          1          0
## 69           2          1          1          2          1          0
## 495          2          1          1          2          1          0
## 679          2          1          1          2          1          0
##     SL253814.y rowcounts_1 rowcounts_2 rowcounts_0 external_gene_name
## 35          NA          11           6           5              ERCC2
## 464         NA          11           6           5              FASTK
## 340         NA          11           6           5                GRN
## 69          NA          11           6           5               RPS9
## 495         NA          11           6           5             ARPC1B
## 679         NA          11           6           5           TBC1D10C
##                                                                                        
description
## 35  Bos taurus ERCC excision repair 2, TFIIH core complex helicase subunit (ERCC2), m
RNA. [Source:RefSeq mRNA;Acc:NM_001103317]
## 464                      Bos taurus Fas activated serine/threonine kinase (FASTK), m
RNA. [Source:RefSeq mRNA;Acc:NM_001035077]
## 340                                                                     granulin pre
cursor [Source:VGNC Symbol;Acc:VGNC:29663]
## 69                                         Bos taurus ribosomal protein S9 (RPS9), m
RNA. [Source:RefSeq mRNA;Acc:NM_001101152]
## 495                                           actin related protein 2/3 complex subu
nit 1B [Source:VGNC Symbol;Acc:VGNC:26164]
## 679                                                             TBC1 domain family memb
er 10C [Source:VGNC Symbol;Acc:VGNC:35625]
##     hgnc_symbol   gene_biotype transcript_length X.Uploaded_variation Location2
## 35              protein_coding              3386      19_14551828_A/T  14551828
## 464             protein_coding              1799      19_14551828_A/T  14551828
## 340             protein_coding              2341      19_14551828_A/T  14551828
## 69              protein_coding              1063      19_14551828_A/T  14551828
## 495             protein_coding              1522      19_14551828_A/T  14551828
## 679             protein_coding              2608      19_14551828_A/T  14551828
##     Allele    Consequence   IMPACT SYMBOL               Gene Feature_type
## 35       T intron_variant MODIFIER  TAF15 ENSBTAG00000006916   Transcript
## 464      T intron_variant MODIFIER  TAF15 ENSBTAG00000006916   Transcript
## 340      T intron_variant MODIFIER  TAF15 ENSBTAG00000006916   Transcript
## 69       T intron_variant MODIFIER  TAF15 ENSBTAG00000006916   Transcript
## 495      T intron_variant MODIFIER  TAF15 ENSBTAG00000006916   Transcript
## 679      T intron_variant MODIFIER  TAF15 ENSBTAG00000006916   Transcript
##            BIOTYPE Existing_variation DISTANCE STRAND  SNPs
## 35  protein_coding        rs135469682        -     -1 SNPdb
## 464 protein_coding        rs135469682        -     -1 SNPdb
## 340 protein_coding        rs135469682        -     -1 SNPdb
## 69  protein_coding        rs135469682        -     -1 SNPdb
## 495 protein_coding        rs135469682        -     -1 SNPdb
## 679 protein_coding        rs135469682        -     -1 SNPdb
```

```
head(eqtl_annotate_anova_TMM)
```

```
##          SNP               gene    F-test    pvalue  q.values bonferroni
## 1 1:1001634 ENSBTAG00000000005 0.1030084 0.9023705 0.6920012          1
## 2 1:1001634 ENSBTAG00000000010 1.5399937 0.2274640 0.4636318          1
## 3 1:1001634 ENSBTAG00000000012 1.4933308 0.2375089 0.4696016          1
## 4 1:1001634 ENSBTAG00000000013 1.7126826 0.1940116 0.4422671          1
## 5 1:1001634 ENSBTAG00000000014 1.3170926 0.2798651 0.4929793          1
## 6 1:1001634 ENSBTAG00000000016 0.2963252 0.7452480 0.6547860          1
##         FDR
## 1 0.9709408
## 2 0.6505178
## 3 0.6588939
## 4 0.6205412
## 5 0.6916949
## 6 0.9187245
```

```
combined_eqtl_annotate_anova_edgeR1_eqtl_annotate_anova_TMM<-merge(eqtl_annotate_anova_e
dgeR1,eqtl_annotate_anova_TMM,by.x=c("snp","gene"),by.y=c("SNP","gene"), all.x=TRUE, al
l.y=FALSE)
#head(combined_eqtl_annotate_anova_edgeR1_eqtl_annotate_anova_TMM)

#write.table(combined_eqtl_annotate_anova_edgeR1_eqtl_annotate_anova_TMM, file= "/mnt/st
orage/lab_folder/shared_R_codes/fernando/SNP_eqtl/results/Supplementary_table_4.txt", ap
pend = FALSE, quote = FALSE, sep = "\t" ,row.names = FALSE)

head(eqtl_annotate_linear_edgeR1)
```

```
##           snp               gene     logFC   logCPM        F       PValue CHROM
## 3 18:57565792 ENSBTAG00000019227 -2.731406 6.675846 66.62621 1.869332e-09    18
## 2 18:57498163 ENSBTAG00000019227  2.634750 6.349960 61.68778 9.618908e-09    18
## 1  11:3860402 ENSBTAG00000032902  1.767669 4.640786 57.50868 1.556154e-08    11
## 6 25:27950377 ENSBTAG00000009914  1.399857 6.454892 65.61085 1.750989e-08    25
## 7 29:40879481 ENSBTAG00000054615  0.894256 7.589831 49.98329 2.310701e-08    29
## 9 4:112942106 ENSBTAG00000009727  1.648550 8.464031 50.96189 3.138249e-08     4
##         POS SL220764.x SL220765.x SL220766.x SL220767.x SL220768.x SL220769.x
## 3  57565792        C/C        A/C        C/C        A/C        A/C        A/C
## 2  57498163        A/A        A/C       <NA>        A/C        A/C       <NA>
## 1   3860402        G/A        G/G        G/G       <NA>        G/G        G/G
## 6  27950377        A/G       <NA>       <NA>       <NA>       <NA>        A/G
## 7  40879481        G/A        G/A        G/A        G/G        G/G        G/A
## 9 112942106        T/C        T/C        T/C       <NA>        T/C        T/C
##   SL220770.x SL220771.x SL220772.x SL220773.x SL220774.x SL220775.x SL253803.x
## 3        A/C        C/C        C/C        C/C        A/C        C/C       <NA>
## 2        C/C       <NA>        A/A        A/A        A/C       <NA>        A/A
## 1        G/A       <NA>        G/G       <NA>       <NA>        G/G        A/A
## 6       <NA>       <NA>       <NA>       <NA>       <NA>       <NA>        A/G
## 7        G/G        G/A        G/A        G/G        G/A        G/G        G/A
## 9       <NA>       <NA>       <NA>       <NA>       <NA>        T/T        T/C
##   SL253804.x SL253805.x SL253806.x SL253807.x SL253808.x SL253809.x SL253810.x
## 3        A/C       <NA>       <NA>        A/C        C/C        A/A        A/C
## 2        A/C        A/A       <NA>        A/C        A/A       <NA>        A/C
## 1        G/A        G/A        A/A        A/A        A/A        A/A        G/A
## 6        A/G        A/A        A/A        A/G       <NA>        A/A        A/A
## 7        G/A        G/A        A/A        G/A        A/A        A/A        A/A
## 9        T/C        T/T        T/C        T/C        T/C       <NA>        C/C
##   SL253811.x SL253812.x SL253813.x SL253814.x SL297951.x SL297952.x SL297953
## 3        C/C        A/C        A/A        A/C        A/C        A/C     <NA>
## 2        A/A        A/C        C/C       <NA>        C/C       <NA>     <NA>
## 1        G/A        G/A        A/A        A/A        A/A       <NA>     <NA>
## 6        A/G        A/A        A/A        A/A        A/A       <NA>     <NA>
## 7        G/A        A/A        G/A        A/A        G/A       <NA>     <NA>
## 9        T/T        T/T        T/C        T/T        T/C       <NA>     <NA>
##   SL297954.x SL297955.x SL297956.x SL297957.x SL297958.x SL297959.x SL297960.x
## 3        A/A        A/A       <NA>        C/C       <NA>        A/C        A/C
## 2        C/C        C/C       <NA>        A/A       <NA>        A/C        A/C
## 1        G/A        A/A       <NA>       <NA>        A/A        A/A        A/A
## 6        A/A        A/A        G/G       <NA>        G/G        G/G        A/G
## 7        G/A        A/A       <NA>        A/A       <NA>        G/A        A/A
## 9        T/T        T/T       <NA>        T/C        C/C        T/C        T/C
##   SL297961.x SL297962.x SL297963.x SL297964.x SL297965.x SL297966.x SL297967.x
## 3       <NA>       <NA>       <NA>        A/C       <NA>        A/A       <NA>
## 2        A/A       <NA>        A/A        A/C        A/C       <NA>       <NA>
## 1       <NA>       <NA>        G/A        A/A       <NA>       <NA>       <NA>
## 6       <NA>       <NA>        A/A       <NA>       <NA>        G/G       <NA>
## 7        A/A       <NA>        G/A        A/A        A/A       <NA>       <NA>
## 9        C/C        C/C        T/C        T/C        C/C       <NA>        C/C
##   SL297968.x SL297951.y SL297952.y SL297954.y SL297955.y SL297956.y SL297957.y
## 3        A/A          1          1          0          0         NA          2
## 2        C/C          2         NA          2          2         NA          0
```

```
## 1         A/A          2          NA          1          2          NA          NA
## 6         G/G          0          NA          0          0           2          NA
## 7         A/A          1          NA          1          2          NA           2
## 9         C/C          1          NA          0          0          NA           1
##   SL297958.y SL297959.y SL297960.y SL297961.y SL297962.y SL297963.y SL297964.y
## 3         NA          1          1         NA         NA         NA          1
## 2         NA          1          1          0         NA          0          1
## 1          2          2          2         NA         NA          1          2
## 6          2          2          1         NA         NA          0         NA
## 7         NA          1          2          2         NA          1          2
## 9          2          1          1          2          2          1          1
##   SL297965.y SL297966.y SL297967.y SL297968.y SL220764.y SL220765.y SL220766.y
## 3         NA          0         NA          0          2          1          2
## 2          1         NA         NA          2          0          1         NA
## 1         NA         NA         NA          2          1          0          0
## 6         NA          2         NA          2          1         NA         NA
## 7          2         NA         NA          2          1          1          1
## 9          2         NA          2          2          1          1          1
##   SL220767.y SL220768.y SL220769.y SL220770.y SL220771.y SL220772.y SL220773.y
## 3          1          1          1          1          2          2          2
## 2          1          1         NA          2         NA          0          0
## 1         NA          0          0          1         NA          0         NA
## 6         NA         NA          1         NA         NA         NA         NA
## 7          0          0          1          0          1          1          0
## 9         NA          1          1         NA         NA         NA         NA
##   SL220774.y SL220775.y SL253803.y SL253804.y SL253805.y SL253806.y SL253807.y
## 3          1          2         NA          1         NA         NA          1
## 2          1         NA          0          1          0         NA          1
## 1         NA          0          2          1          1          2          2
## 6         NA         NA          1          1          0          0          1
## 7          1          0          1          1          1          2          1
## 9         NA          0          1          1          0          1          1
##   SL253808.y SL253809.y SL253810.y SL253811.y SL253812.y SL253813.y SL253814.y
## 3          2          0          1          2          1          0          1
## 2          0         NA          1          0          1          2         NA
## 1          2          2          1          1          1          2          2
## 6         NA          0          0          1          0          0          0
## 7          2          2          2          1          2          1          2
## 9          1         NA          2          0          0          1          0
##   rowcounts_1 rowcounts_2 rowcounts_0 external_gene_name
## 3          16           9           6           SIGLEC14
## 2          12           6          10           SIGLEC14
## 1           9          14           6            SLC11A2
## 6           7           5          11             NDUFS6
## 7          17          13           5 ENSBTAG00000054615
## 9          17           7           7            ADIPOR1
##
description
## 3                                                     sialic acid-binding Ig-like lect
in 14 [Source:NCBI gene (formerly Entrezgene);Acc:789748]
## 2                                                     sialic acid-binding Ig-like lect
in 14 [Source:NCBI gene (formerly Entrezgene);Acc:789748]
```

```
## 1 solute carrier family 11 (proton-coupled divalent metal ion transporters), member 2
-like [Source:NCBI gene (formerly Entrezgene);Acc:512464]
## 6                                                                  NADH:ubiquinone oxid
oreductase subunit S6 [Source:VGNC Symbol;Acc:VGNC:31971]
## 7
## 9                                                                                     a
diponectin receptor 1 [Source:VGNC Symbol;Acc:VGNC:25678]
##   hgnc_symbol   gene_biotype transcript_length chromosome_name start_position
## 3             protein_coding              2011              18       57637619
## 2             protein_coding              2011              18       57637619
## 1             protein_coding              2201               5       28846446
## 6             protein_coding              1291              20       70839103
## 7             protein_coding               393              20        3252364
## 9             protein_coding              1984              16       79870488
##   end_position strand X.Uploaded_variation Location2 Allele
## 3     57641293     -1      18_57565792_A/C  57565792      C
## 2     57641293     -1      18_57498163_A/C  57498163      C
## 1     28868817      1       11_3860402_G/A   3860402      A
## 6     70844383     -1      25_27950377_A/G  27950377      G
## 7      3252756     -1      29_40879481_G/A  40879481      A
## 9     79883989      1      4_112942106_T/C 112942106      C
##              Consequence   IMPACT SYMBOL               Gene Feature_type
## 3        missense_variant MODERATE      - ENSBTAG00000045880   Transcript
## 2 downstream_gene_variant MODIFIER      - ENSBTAG00000047675   Transcript
## 1      synonymous_variant      LOW MGAT4A ENSBTAG00000010388   Transcript
## 6        missense_variant MODERATE      - ENSBTAG00000055247   Transcript
## 7      synonymous_variant      LOW  AHNAK ENSBTAG00000013468   Transcript
## 9          intron_variant MODIFIER      - ENSBTAG00000039588   Transcript
##          BIOTYPE Existing_variation DISTANCE STRAND  SNPs
## 3 protein_coding         rs41892216        -     -1 SNPdb
## 2 protein_coding        rs135008768      483     -1 SNPdb
## 1 protein_coding        rs110261991        -     -1 SNPdb
## 6 protein_coding        rs439610235        -      1 SNPdb
## 7 protein_coding         rs42188410        -     -1 SNPdb
## 9 protein_coding        rs450283929        -     -1 SNPdb
```

```
head(eqtl_annotate_linear_TMM)
```

```
##         SNP               gene         beta      t-stat   p-value
## 1 1:1001634 ENSBTAG00000000005 -0.046765337 -0.19300556 0.8479562
## 2 1:1001634 ENSBTAG00000000010  0.190227150  0.79098246 0.4337383
## 3 1:1001634 ENSBTAG00000000012 -0.006538202 -0.02697123 0.9786202
## 4 1:1001634 ENSBTAG00000000013 -0.116077517 -0.48024952 0.6337328
## 5 1:1001634 ENSBTAG00000000014  0.206918860  0.86165522 0.3941444
## 6 1:1001634 ENSBTAG00000000016 -0.003125273 -0.01289221 0.9897795
```

```
colnames(eqtl_annotate_linear_TMM)<-c("SNP","gene","beta","t-stat",  "Pvalue_linear")

combined_eqtl_eqtl_annotate_linear_edgeR1_eqtl_annotate_linear_TMM1<-merge(eqtl_annotate
_linear_edgeR1,eqtl_annotate_linear_TMM,by.x=c("snp","gene"),by.y=c("SNP","gene"), all =
FALSE)

#write.table(combined_eqtl_eqtl_annotate_linear_edgeR1_eqtl_annotate_linear_TMM1, file=
 "/mnt/storage/lab_folder/shared_R_codes/fernando/SNP_eqtl/results/Supplementary_table_4
_a.txt", append = FALSE, quote = FALSE, sep = "\t" ,row.names = FALSE)
```

#Supplementary figure 3

```
plot_1<-ggplot(data=combined_eqtl_annotate_anova_edgeR1_eqtl_annotate_anova_TMM, aes(x=-
log10(PValue.anova), y=-log10(pvalue)))+
  scale_x_continuous(name="-Log10(P value) DEG framework")+
  scale_y_continuous(name="-Log10(P value) standard framework", breaks=c(0:7))+
  geom_point()+
  ggtitle("ANOVA model")+
  theme_bw(base_size = 13)+
  theme(
    axis.text=element_text(size=13, color="black")
  )
plot_2<-ggplot(data=combined_eqtl_eqtl_annotate_linear_edgeR1_eqtl_annotate_linear_TMM1,
aes(x=-log10(PValue), y=-log10(Pvalue_linear)))+
  scale_x_continuous(name="-Log10(P value) DEG framework")+
  scale_y_continuous(name="-Log10(P value) standard framework", breaks=c(0:7))+
  geom_point()+
  ggtitle("Additive model")+
  theme_bw(base_size = 13)+
  theme(
    axis.text=element_text(size=13, color="black")
  )

cowplot::plot_grid( plot_1, plot_2, ncol = 2, labels = c("A", "B") ,label_size = 12)
```

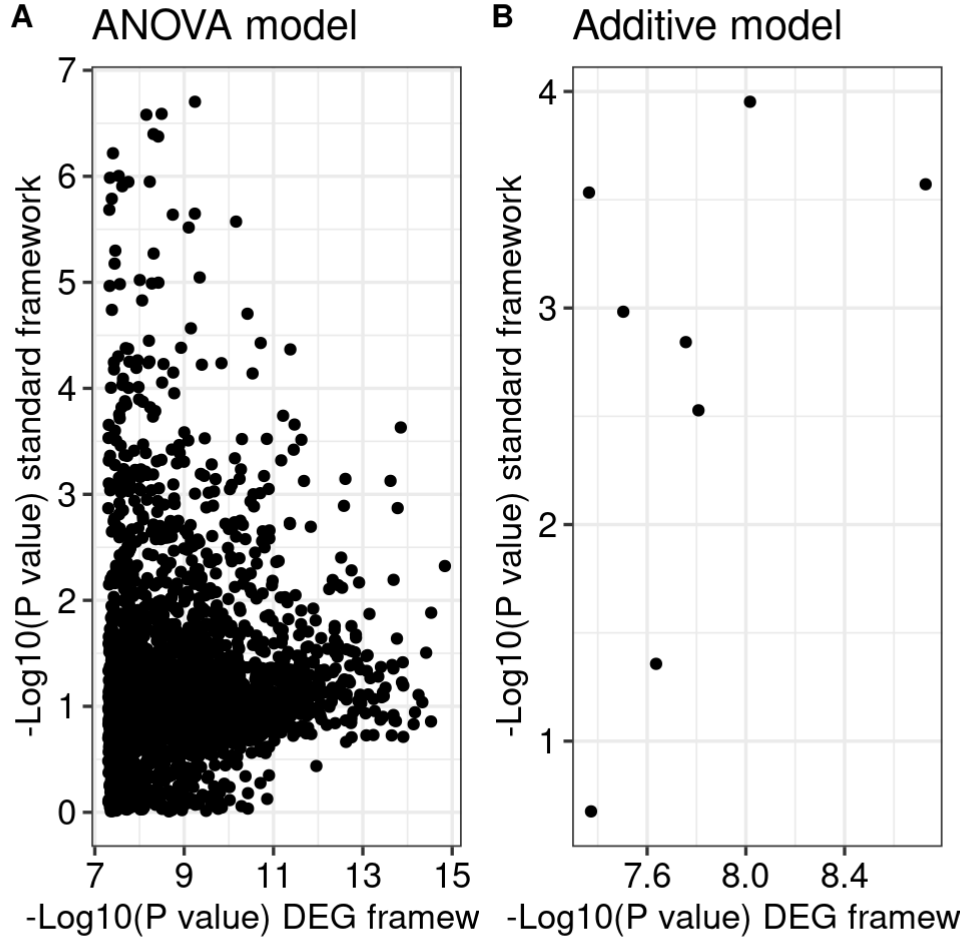


#Supplementary figure 4

```
font_size=12

snps_sub2 = eqtl_annotate_anova_edgeR1$snp[1:7]
genes_sub2 = eqtl_annotate_anova_edgeR1$gene[1:7]
gene_symbol = eqtl_annotate_anova_edgeR1$external_gene_name[1:7]

rm(graph_data_frame)

plot_list_1<-list()

for (index in seq(length(snps_sub2))){
  genotype_sub2 = unlist((merged_SNPS_nucleotide[snps_sub2[index],c(3:44)]))
  genotype_sub2 = genotype_sub2[!(genotype_sub2=="<NA>")]
  expression_sub2 = tpm_merged_datasets[genes_sub2[index],]
  expression_sub2 = expression_sub2[names(expression_sub2) %in% names(genotype_sub2) ]
  genotype_sub2<-genotype_sub2[names(expression_sub2)]

  graph_data_frame<-data.frame(genotype_sub2=gsub("/","",genotype_sub2),expression_sub2,
snp=snps_sub2[index], gene_symbol=gene_symbol[index])
  graph_data_frame<-graph_data_frame[complete.cases(graph_data_frame),]
  graph_data_frame<-graph_data_frame[!(graph_data_frame$genotype_sub2 =="<NA>"),]

  plot_list_1[[index]]<-ggplot(data=graph_data_frame, aes(x=as.factor(genotype_sub2), y=
expression_sub2))+
    geom_boxplot(fill='transparent',outlier.shape = 4, outlier.color = "blue", size=0.1)
+
    geom_jitter(width=0.2, size=1)+
    scale_x_discrete(name= graph_data_frame$snp[1])+
    scale_y_continuous(name =  graph_data_frame$gene_symbol[1])+
    theme_classic(base_size = font_size)+
    theme( axis.text=element_text(size=font_size,color="black"),
           axis.title=element_text(size=font_size,color="black"),
           plot.margin =unit(c(0.1,2,0.1,0.1), "cm"))
}

merged_datasets<-as.matrix(merged_datasets)

rm(graph_data_frame)

plot_list_2<-list()

for (index in seq(length(snps_sub2))){
  genotype_sub2 = unlist((merged_SNPS_nucleotide[snps_sub2[index],c(3:44)]))
  genotype_sub2 = genotype_sub2[!(genotype_sub2=="<NA>")]
  expression_sub2 = merged_datasets[genes_sub2[index],]
  expression_sub2 = expression_sub2[names(expression_sub2) %in% names(genotype_sub2) ]
  genotype_sub2<-genotype_sub2[names(expression_sub2)]

  graph_data_frame<-data.frame(genotype_sub2=gsub("/","",genotype_sub2),expression_sub2,
snp=snps_sub2[index], gene_symbol=gene_symbol[index])
  graph_data_frame<-graph_data_frame[complete.cases(graph_data_frame),]
  graph_data_frame<-graph_data_frame[!(graph_data_frame$genotype_sub2 =="<NA>"),]
```

```
  plot_list_2[[index]]<-ggplot(data=graph_data_frame, aes(x=as.factor(genotype_sub2), y=
expression_sub2))+
    geom_boxplot(fill='transparent',outlier.shape = 4, outlier.color = "blue", size=0.1)
+
    geom_jitter(width=0.2, size=1)+
    scale_x_discrete(name= graph_data_frame$snp[1])+
    scale_y_continuous(name =  graph_data_frame$gene_symbol[1])+
    theme_classic(base_size = font_size)+
    theme( axis.text=element_text(size=font_size,color="black"),
           axis.title=element_text(size=font_size,color="black"),
           plot.margin =unit(c(0.1,2,0.1,0.1), "cm"))
}

tmm_per_million_tmm_per_million_normalized_expression<-as.matrix(tmm_per_million_tmm_per
_million_normalized_expression)

rm(graph_data_frame)

plot_list_3<-list()

for (index in seq(length(snps_sub2))){
  genotype_sub2 = unlist((merged_SNPS_nucleotide[snps_sub2[index],c(3:44)]))
  genotype_sub2 = genotype_sub2[!(genotype_sub2=="<NA>")]
  expression_sub2 = tmm_per_million_tmm_per_million_normalized_expression[genes_sub2[ind
ex],]
  expression_sub2 = expression_sub2[names(expression_sub2) %in% names(genotype_sub2) ]
  genotype_sub2<-genotype_sub2[names(expression_sub2)]

  graph_data_frame<-data.frame(genotype_sub2=gsub("/","",genotype_sub2),expression_sub2,
snp=snps_sub2[index], gene_symbol=gene_symbol[index])
  graph_data_frame<-graph_data_frame[complete.cases(graph_data_frame),]
  graph_data_frame<-graph_data_frame[!(graph_data_frame$genotype_sub2 =="<NA>"),]

  plot_list_3[[index]]<-ggplot(data=graph_data_frame, aes(x=as.factor(genotype_sub2), y=
expression_sub2))+
    geom_boxplot(fill='transparent',outlier.shape = 4, outlier.color = "blue", size=0.1)
+
    geom_jitter(width=0.2, size=1)+
    scale_x_discrete(name= graph_data_frame$snp[1])+
    scale_y_continuous(name =  graph_data_frame$gene_symbol[1])+
    theme_classic(base_size = font_size)+
    theme( axis.text=element_text(size=font_size,color="black"),
           axis.title=element_text(size=font_size,color="black"),
           plot.margin =unit(c(0.1,2,0.1,0.1), "cm"))
}

rm(graph_data_frame)


plot_list_4<-list()
```

```
for (index in seq(length(snps_sub2))){
  genotype_sub2 = unlist((merged_SNPS_nucleotide[snps_sub2[index],c(3:44)]))
  genotype_sub2 = genotype_sub2[!(genotype_sub2=="<NA>")]
  expression_sub2 = TMM_tmm_per_million_normalized_expression_a[genes_sub2[index],]
  expression_sub2 = expression_sub2[names(expression_sub2) %in% names(genotype_sub2) ]
  genotype_sub2<-genotype_sub2[names(expression_sub2)]

  graph_data_frame<-data.frame(genotype_sub2=gsub("/","",genotype_sub2),expression_sub2,
snp=snps_sub2[index], gene_symbol=gene_symbol[index])
  graph_data_frame<-graph_data_frame[complete.cases(graph_data_frame),]
  graph_data_frame<-graph_data_frame[!(graph_data_frame$genotype_sub2 =="<NA>"),]

  plot_list_4[[index]]<-ggplot(data=graph_data_frame, aes(x=as.factor(genotype_sub2), y=
expression_sub2))+
    geom_boxplot(fill='transparent',outlier.shape = 4, outlier.color = "blue", size=0.1)
+
    geom_jitter(width=0.2, size=1)+
    scale_x_discrete(name= graph_data_frame$snp[1])+
    scale_y_continuous(name =  graph_data_frame$gene_symbol[1])+
    theme_classic(base_size = font_size)+
    theme( axis.text=element_text(size=font_size,color="black"),
           axis.title=element_text(size=font_size,color="black"),
           plot.margin =unit(c(0.1,2,0.1,0.1), "cm"))
}


y.grob.TPM <- textGrob("Transcript per million", gp=gpar(fontface="plain", col="black",
 fontsize=font_size), rot=90)

y.grob.count <- textGrob("Raw counts", gp=gpar(fontface="plain", col="black", fontsize=f
ont_size), rot=90)

y.grob.TMM <- textGrob("TMM normalized counts per million", gp=gpar(fontface="plain", co
l="black", fontsize=font_size), rot=90)

y.grob.TMM_normal <- textGrob("TMM normalized counts per million and normal transformed"
, gp=gpar(fontface="plain", col="black", fontsize=font_size), rot=90)


grid.arrange(arrangeGrob(cowplot::plot_grid(plotlist= plot_list_2  , ncol=1), left = y.g
rob.count),
             arrangeGrob(cowplot::plot_grid(plotlist= plot_list_1  , ncol=1), left = y.g
rob.TPM),
             arrangeGrob(cowplot::plot_grid(plotlist= plot_list_3  , ncol=1), left = y.g
rob.TMM),
             arrangeGrob(cowplot::plot_grid(plotlist= plot_list_4  , ncol=1), left = y.g
rob.TMM_normal),ncol=4)
```

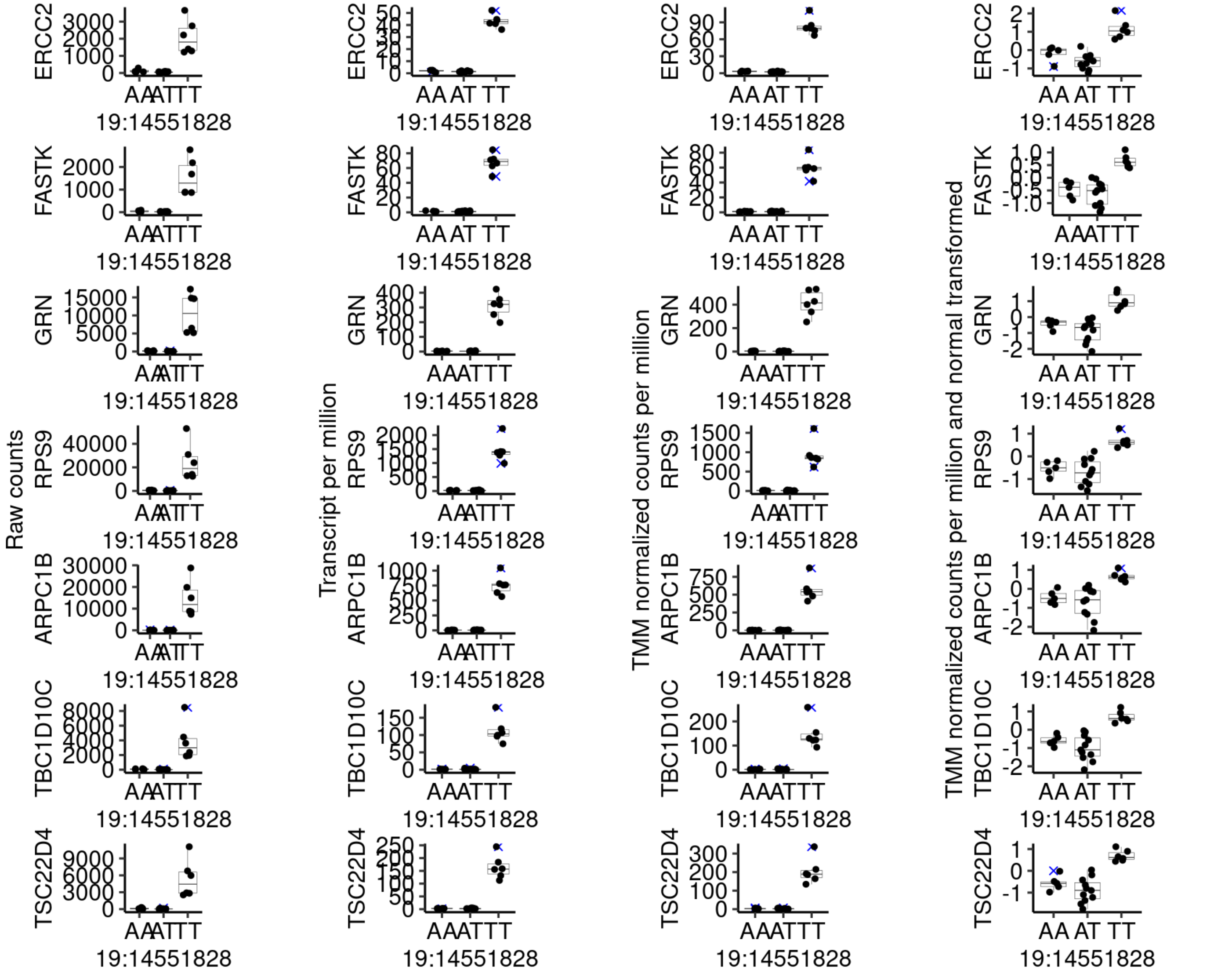


#Supplementary table 1 eqtl

```
# TMM

TMM_annotation <- SNP_annotation_all_a[(SNP_annotation_all_a$SNP %in% eqtl_annotate_anova_TMM1$SNP),]

TMM_annotation$SNPs<-ifelse(TMM_annotation$Existing_variation=="-","putative_new", "SNPdb")


df2= TMM_annotation %>% dplyr::group_by(SNPs, Consequence) %>% summarize(n=n()) %>% arrange(desc(SNPs),desc(n))
```

```
## `summarise()` has grouped output by 'SNPs'. You can override using the
## `.groups` argument.
```

```
n_total_size_SNPdb<-sum(df2$n[1:7])


df2$percentage <- round(df2$n[1:7]/n_total_size_SNPdb * 100, 2)

#raw counts
count_annotation <- SNP_annotation_all_a[(SNP_annotation_all_a$SNP %in% results_anova_do
minance$snp),]

count_annotation$SNPs<-ifelse(count_annotation$Existing_variation=="-","putative_new",
"SNPdb")

df3= count_annotation %>% dplyr::group_by(SNPs, Consequence) %>% summarize(n=n()) %>% ar
range(desc(SNPs),desc(n))
```

```
## `summarise()` has grouped output by 'SNPs'. You can override using the
## `.groups` argument.
```

```
n_total_size_SNPdb<-sum(df3$n[1:7])


df3$percentage <- round(df3$n[1:7]/n_total_size_SNPdb * 100, 2)

SNP_consequence_table_comparisons <- rbind(df1,df2,df3)

all_snps<-rbind(count_annotation,TMM_annotation)
all_snps<-all_snps[!duplicated(all_snps$SNP),]
#table(all_snps$SNPs)
#write.table(SNP_consequence_table_comparisons,"/mnt/storage/lab_folder/shared_R_codes/f
ernando/SNP_eqtl/results/SNP_consequence_table_comparisons.txt",col.names = TRUE, row.na
mes = TRUE, quote = FALSE,sep ="\t")
```

```
 sessionInfo()
```

```
## R version 4.2.2 Patched (2022-11-10 r83330)
## Platform: x86_64-pc-linux-gnu (64-bit)
## Running under: Ubuntu 20.04.5 LTS
##
## Matrix products: default
## BLAS:   /usr/lib/x86_64-linux-gnu/blas/libblas.so.3.9.0
## LAPACK: /usr/lib/x86_64-linux-gnu/lapack/liblapack.so.3.9.0
##
## locale:
##  [1] LC_CTYPE=en_US.UTF-8       LC_NUMERIC=C
##  [3] LC_TIME=en_US.UTF-8        LC_COLLATE=en_US.UTF-8
##  [5] LC_MONETARY=en_US.UTF-8    LC_MESSAGES=en_US.UTF-8
##  [7] LC_PAPER=en_US.UTF-8       LC_NAME=C
##  [9] LC_ADDRESS=C               LC_TELEPHONE=C
## [11] LC_MEASUREMENT=en_US.UTF-8 LC_IDENTIFICATION=C
##
## attached base packages:
##  [1] grid      stats4    parallel  stats     graphics  grDevices utils
##  [8] datasets  methods   base
##
## other attached packages:
##  [1] gridExtra_2.3                    HardyWeinberg_1.7.5
##  [3] nnet_7.3-18                      Rsolnp_1.16
##  [5] mice_3.14.0                      RNOmni_1.0.1
##  [7] dplyr_1.0.10                     tidyr_1.2.1
##  [9] ggforce_0.4.1                    goseq_1.48.0
## [11] geneLenDataBase_1.32.0           BiasedUrn_2.0.8
## [13] cowplot_1.1.1                    readxl_1.4.1
## [15] sjmisc_2.8.9                     ggplot2_3.3.6
## [17] edgeR_3.38.4                     limma_3.52.1
## [19] DESeq2_1.36.0                    SummarizedExperiment_1.26.1
## [21] MatrixGenerics_1.8.0             matrixStats_0.62.0
## [23] GenomicRanges_1.48.0             GenomeInfoDb_1.32.2
## [25] IRanges_2.30.0                   S4Vectors_0.34.0
## [27] GenomicTools_0.2.9.7             GenomicTools.fileHandler_0.1.5.9
## [29] data.table_1.14.4                gMWT_1.1.1
## [31] Rcpp_1.0.9                       clinfun_1.1.0
## [33] MatrixEQTL_2.3                   Biobase_2.56.0
## [35] BiocGenerics_0.42.0              doParallel_1.0.17
## [37] iterators_1.0.14                 foreach_1.5.2
## [39] qvalue_2.28.0                    stringr_1.4.1
## [41] reshape2_1.4.4                   R6_2.5.1
## [43] vcfR_1.13.0
##
## loaded via a namespace (and not attached):
##   [1] backports_1.4.1          circlize_0.4.15          BiocFileCache_2.4.0
##   [4] plyr_1.8.7               splines_4.2.2            BiocParallel_1.30.4
##   [7] digest_0.6.30            htmltools_0.5.3          GO.db_3.15.0
##  [10] fansi_1.0.3              magrittr_2.0.3           memoise_2.0.1
##  [13] cluster_2.1.4            Biostrings_2.64.1        annotate_1.74.0
##  [16] R.utils_2.12.0           prettyunits_1.1.1        colorspace_2.0-3
##  [19] rappdirs_0.3.3           blob_1.2.3               xfun_0.34
```

```
##  [22] crayon_1.5.2            RCurl_1.98-1.7          jsonlite_1.8.3
##  [25] genefilter_1.78.0       survival_3.3-1          ape_5.6-2
##  [28] glue_1.6.2              polyclip_1.10-4         gtable_0.3.1
##  [31] zlibbioc_1.42.0         XVector_0.36.0          DelayedArray_0.22.0
##  [34] shape_1.4.6             scales_1.2.1            mvtnorm_1.1-3
##  [37] DBI_1.1.3               viridisLite_0.4.1       xtable_1.8-4
##  [40] progress_1.2.2          bit_4.0.4               truncnorm_1.0-8
##  [43] httr_1.4.4              RColorBrewer_1.1-3      ellipsis_0.3.2
##  [46] R.methodsS3_1.8.2       farver_2.1.1            pkgconfig_2.0.3
##  [49] XML_3.99-0.11           sass_0.4.2              dbplyr_2.2.0
##  [52] locfit_1.5-9.6          utf8_1.2.2              labeling_0.4.2
##  [55] tidyselect_1.2.0        rlang_1.0.6             AnnotationDbi_1.58.0
##  [58] munsell_0.5.0           cellranger_1.1.0        tools_4.2.2
##  [61] cachem_1.0.6            cli_3.4.1               generics_0.1.3
##  [64] RSQLite_2.2.18          sjlabelled_1.2.0        broom_1.0.1
##  [67] evaluate_0.17           fastmap_1.1.0           yaml_2.3.6
##  [70] knitr_1.40              bit64_4.0.5             purrr_0.3.5
##  [73] KEGGREST_1.36.3         nlme_3.1-160            mime_0.12
##  [76] R.oo_1.25.0             xml2_1.3.3              biomaRt_2.52.0
##  [79] compiler_4.2.2          rstudioapi_0.14         filelock_1.0.2
##  [82] curl_4.3.3              png_0.1-7               tweenr_2.0.2
##  [85] tibble_3.1.8            geneplotter_1.74.0      bslib_0.4.0
##  [88] stringi_1.7.8           highr_0.9               GenomicFeatures_1.48.3
##  [91] lattice_0.20-45         Matrix_1.5-1            vegan_2.6-4
##  [94] permute_0.9-7           vctrs_0.5.0             pillar_1.8.1
##  [97] lifecycle_1.0.3         jquerylib_0.1.4         GlobalOptions_0.1.2
## [100] snpStats_1.46.0         bitops_1.0-7            insight_0.18.6
## [103] rtracklayer_1.56.0      BiocIO_1.6.0            codetools_0.2-18
## [106] MASS_7.3-58.1           assertthat_0.2.1        rjson_0.2.21
## [109] withr_2.5.0             pinfsc50_1.2.0          GenomicAlignments_1.32.0
## [112] Rsamtools_2.12.0        GenomeInfoDbData_1.2.8  mgcv_1.8-41
## [115] hms_1.1.2               rmarkdown_2.17          restfulr_0.0.14
```
