## Additional file 3 for "Robust identification of regulatory variants (eQTLs) using a differential expression framework developed for RNA-sequencing"

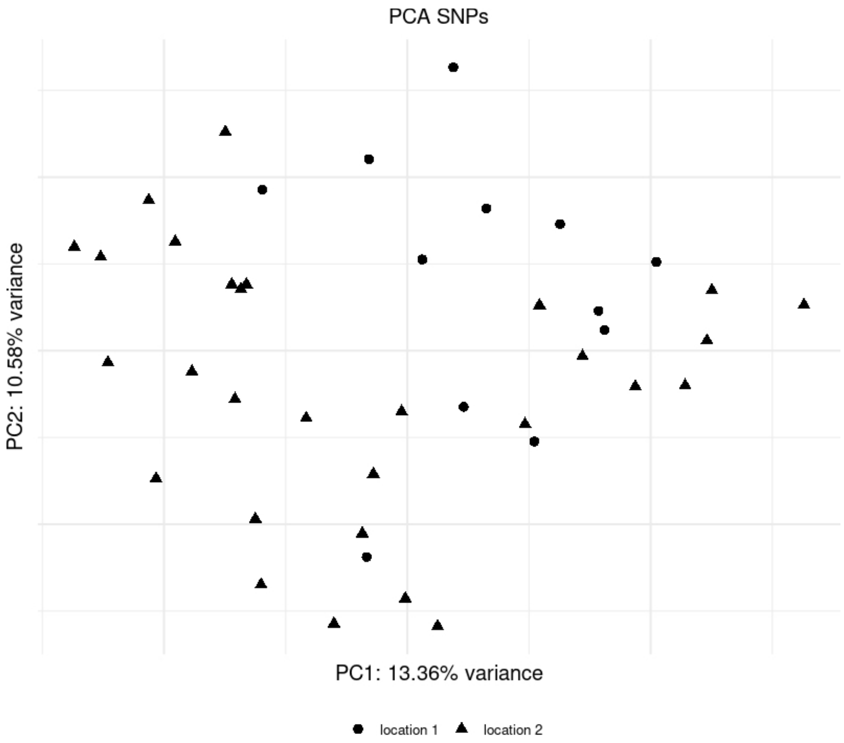


Figure S1. Principal component analysis of the samples based on the SNP data.

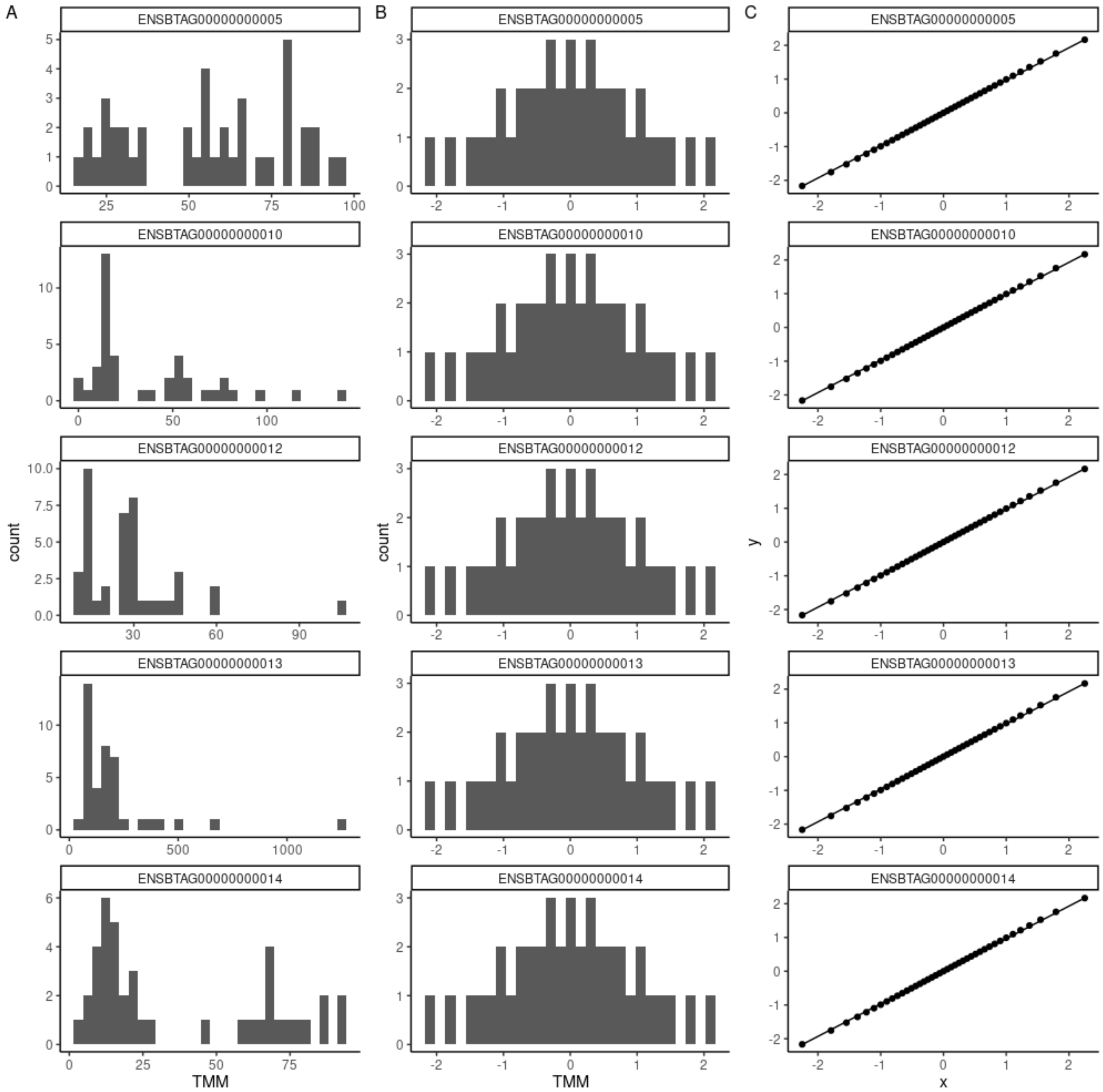


Figure S2. Representation of RNA-Sequencing data after (A) normalization of the count data with the TMM method and adjustment per million reads, and normalization as demonstrated by the (B) histogram and (C) qqplot.

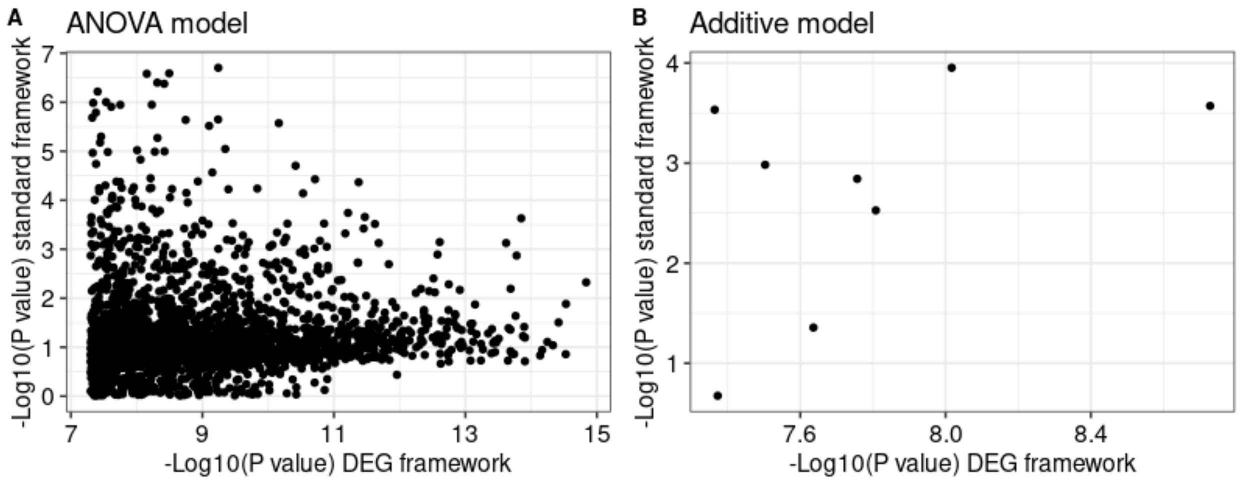


Supplementary figure 3. Scatterplot of the of the raw P values (-$Log_{10}$() transformed) for the eQTLs following the (A) ANOVA model, and (B) additive model.

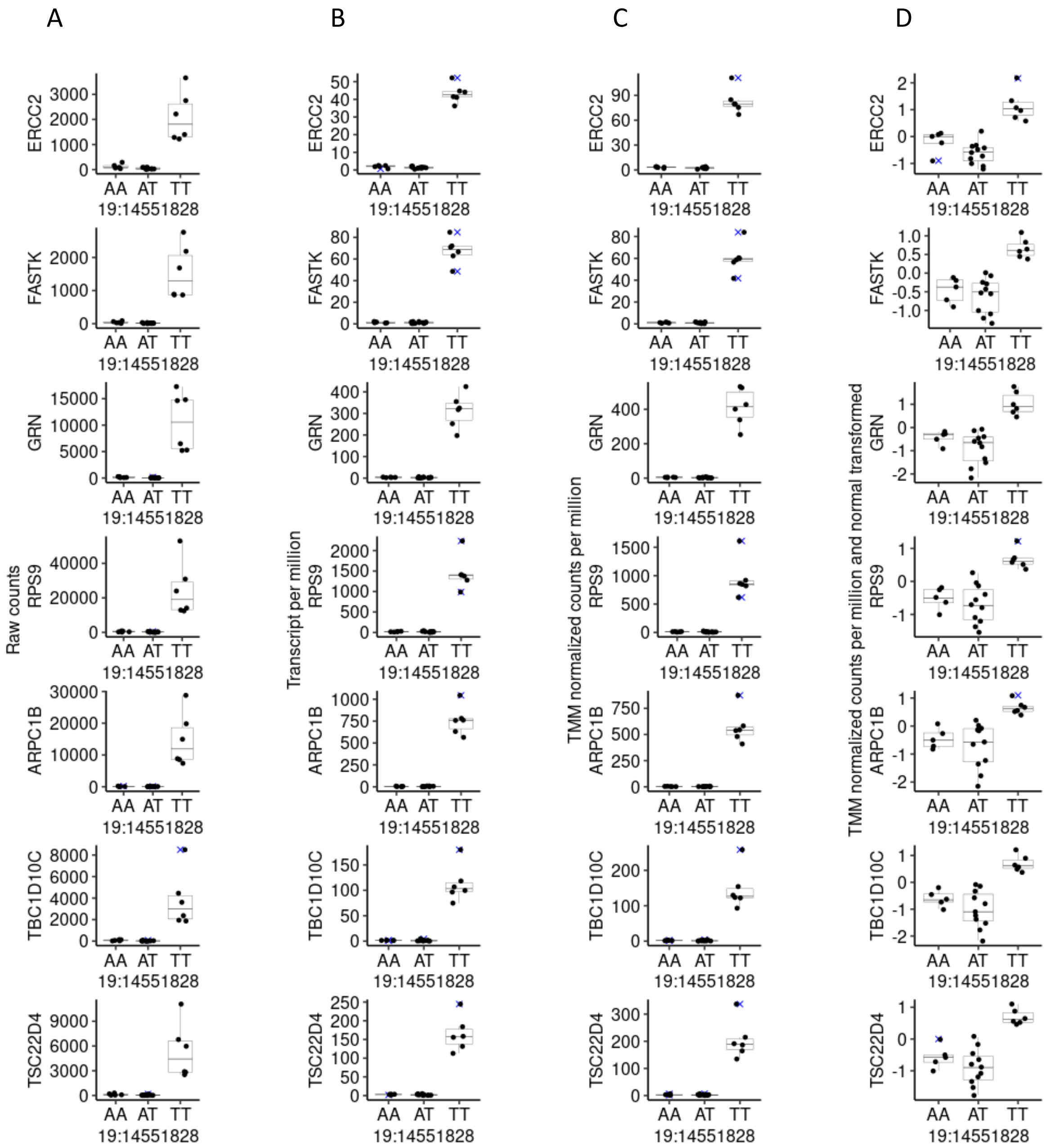


Supplementary figure 4. Plots of significant eQTLs following the dominance mode of allelic interaction identified by the DGE framework. (A) Raw counts (B) Transcript per million (C) TMM normalized counts per million. (D) TMM normalized counts per million and normal transformed.
